# Comparative atlas of genome-wide chromatin-associated protein co-occupancy

**DOI:** 10.1101/2024.12.17.628199

**Authors:** Shannon M. White, Belle A. Moyers, Tao Wang, Mark Mackiewicz, Annika K. Weimer, Fabian Grubert, Vivekanandan Ramalingam, Jay X. J. Luo, Lixia Jiang, Minyi Shi, Xinqiong Yang, Tristan Chou, Jie Zhai, Konor Von Kraut, Jessika Adrian, E. Christopher Partridge, Kristina Paul, Anshul Kundaje, Eric M. Mendenhall, Richard M. Myers, Michael P. Snyder

## Abstract

Accurate transcriptional regulation and chromatin dynamics requires the coordination and activity of chromatin-associated proteins (CAPs) at distinct loci. While the combinatorial activity of a select set of CAPs has been previously examined, these studies are limited by the underrepresentation of proteins and cell types explored, making it difficult to identify the global associations as well as the conservation of these associations across different cell types. Here, we performed 270 CAP chromatin immunoprecipitation followed by high-throughput sequencing (ChIP-Seq) experiments in both K562 and HepG2 cancer cell lines and explored the relationship between cell identity and CAP co-association using three distinct approaches. We employed a machine learning algorithm to organize the genome-wide binding profiles into 56 and 70 interpretable co-association modules for HepG2 and K562 cell lines, respectively. We found CAP co-association modules are mostly cell type-specific, however those present in both cell lines are largely comprised of TFs from a single TF family and anchor to unique loci via lineage-specific factors. While enhancer-associated co-binding modules were largely composed of cell type-specific CAPs, we found regulatory activity at promoter-enhancer module contacts to be enriched for chromatin remodeling proteins. Additionally, we used colocalization information derived from co-association models in conjunction with neural network models of transcription factor (TF) activity to identify high-confidence candidate TF cooperative pairs. Finally, through comparing CAP enrichment in high occupancy target (HOT) regions in K562 and HepG2 cell lines, we found cell type-specific HOT sites, but not common HOT sites, are selectively enriched at high copy number loci. Overall, this study uncovers principles of sequence-level and large-scale CAP genomic organization and demonstrates how this contributes to cell type-specific regulatory mechanisms and cellular functions.

## INTRODUCTION

Transcription factors (TFs), transcriptional cofactors, RNA-binding proteins (RBPs), histone-regulating proteins, and chromatin-remodeling enzymes comprise chromatin-associated proteins (CAPs) which collectively regulate cellular gene expression. TFs bind to short stretches of specific DNA sequences, known as motifs, at cis-regulatory regions and often operate in a combinatorial manner, recruiting several additional CAPs to regulate transcription. These cooperative transcriptional programs have been shown to be critical for several cellular functions including cell differentiation^1,2^, tissue development^3,4^, response to stimuli^5^, and disease progression^6^.

The ability to study complex CAP co-occupancy patterns has been facilitated by chromatin immunoprecipitation followed by high-throughput sequencing (ChIP-seq), which identifies CAP binding locations genome-wide^7,8^. Over the past decade, the Encyclopedia of DNA elements (ENCODE) consortium has published extensive ChIP-seq data across a handful of widely characterized cancer cell lines, which has led to significant insights regarding genome-wide DNA binding specificity and co-occupancy^9^. For example, the discovery and investigation of high occupancy target (HOT) loci^10,11^, the catalog of candidate cis-regulatory elements (cCREs)^12^, and several TF regulatory networks^13^ often utilized the comprehensive ENCODE ChIP-seq database. Additionally, we and others have utilized this expansive dataset to begin dissecting the complex co-binding dynamics that occur within and across cell types^14,15^.

Previous studies have used different techniques to computationally dissect the co-binding patterns of TFs in high dimensional datasets including focusing on all regions bound by a specific CAP and then analyzing all co-bound patterns^15^ or generating self-organizing maps (SOMs)^14^, which uses a hard-clustering algorithm to visualize the colocalization patterns of CAPs. Despite the many advances that have been made, co-occupancy studies have been difficult to comprehend across different cell types due to the highly complex and large number of co-binding patterns discovered and a paucity of CAP binding data using multiple cell types. Moreover, an extensive catalog of co-occupancy patterns across multiple cell lines is nearly impossible due to the sheer number of all possible comparisons. To address this, we have implemented the Regulatory Module Discovery (RMD) algorithm^16^ which uses topic modeling to systematically cluster co-occupied regions into regulatory modules defined by CAP co-occupancy. Importantly, this methodology results in orders of magnitude fewer regulatory modules than those obtained from SOMs, providing less specific but overall more interpretable co-binding patterns.

In order to understand the conserved and cell type-specific principles of CAP co-association, we identified regulatory modules from co-bound loci using 270 CAP ChIP-seq datasets that we generated across two cell lines, K562 and HepG2, as part of the ENCODE project. This extensive analysis in two highly characterized cell lines enabled the identification of common and cell type-specific co-occupancy patterns that are associated with specific chromatin state features, gene expression networks, and 3D chromatin interactions. We used regulatory modules to generate hypotheses about unknown TF cooperative associations, which we tested *in silico* using base-resolution neural network models of TF activity^17^. Finally, we expanded our analysis beyond regulatory modules to evaluate HOT site characteristics and dependencies across cell lines. These datasets and analyses reveal common and cell type-specific co-association modules and their potential functions across different cell types. Overall, these CAP datasets and co-occupancy maps provide a valuable resource for the scientific community.

## RESULTS

### Identification of chromatin-associated protein co-association modules in the extensively characterized K562 and HepG2 cell lines

In effort to conduct a large-scale comparison of the localization of chromatin-associated proteins (CAPs) across cell types, we generated 2,630 ChIP-seq datasets as part of the ENCODE Consortium. Among the 86 human cell lines with CAP ChIP-seq experiments, K562 and HepG2 cancer cell lines were the most extensively characterized, with 502 and 664 uniquely targeted experiments in total, respectively, and therefore were chosen to be the focus of our comparative analysis. These represent different tumor cell lines, as HepG2 was derived from a hepatocellular carcinoma tissue sample and K562 was derived from lymphoblast cells isolated from bone marrow of a patient with chronic myeloid leukemia^18,19^. To facilitate co-association comparisons, we restricted our analysis to the 270 CAPs with high quality ChIP-seq experiments conducted in both K562 and HepG2 cell lines **(Extended Data Fig. 1a-b)**. Among these CAPs, 191 (71%) are DNA-binding transcription factors (TFs) and 198 (73.3%) of the ChIP-seq datasets were generated in ENCODE phase 4 with the remainder from previous ENCODE efforts **(Fig. 1a and Extended Data Fig. 1c)**.

**Figure 1.**
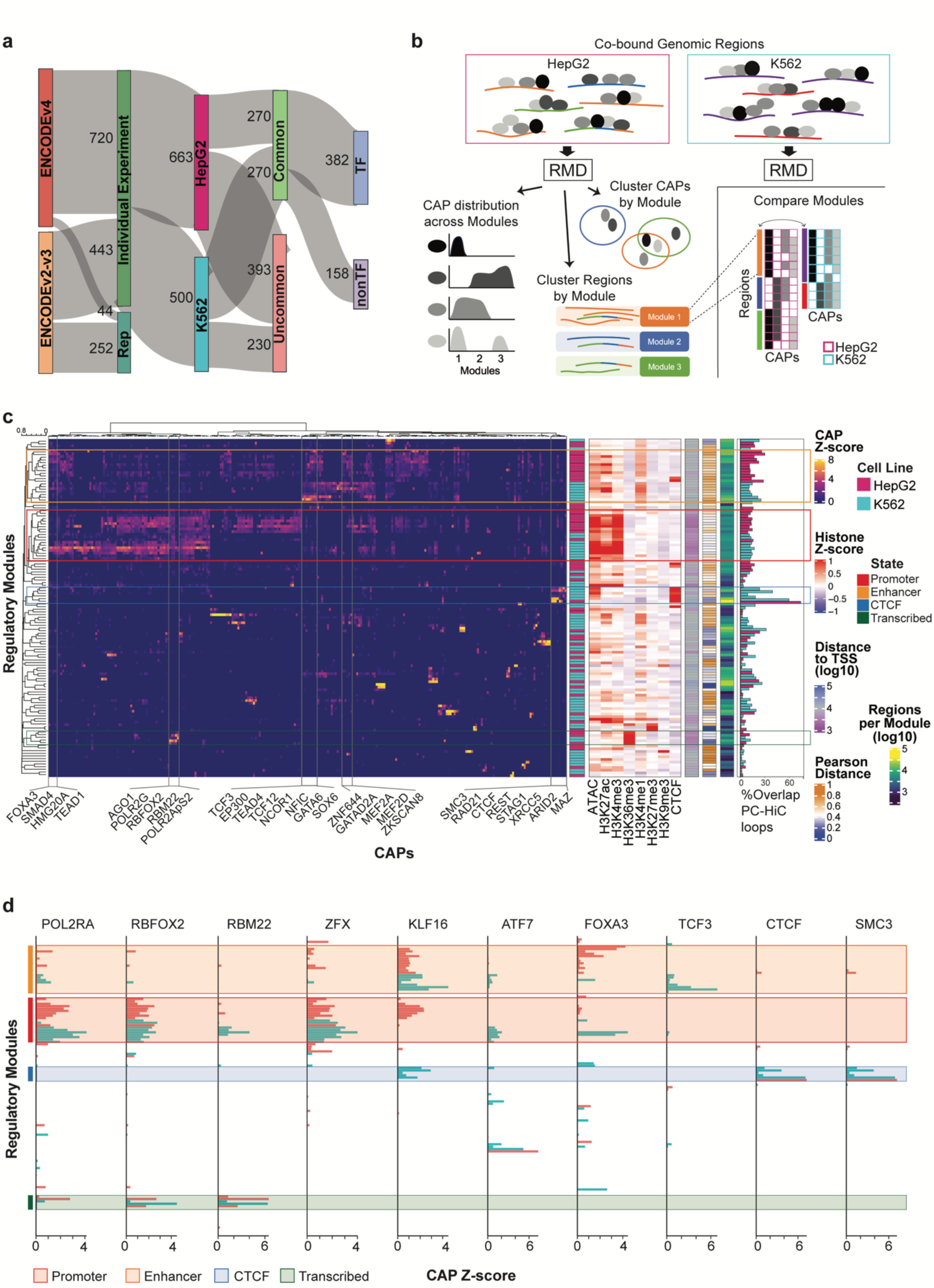
Identification of chromatin-associated protein co-binding modules in the extensively characterized K562 and HepG2 cell lines. a. Description of ChIP-seq datasets used in the comparative analysis. Sankey plot depicting the number of ChIP-seq targets experiments available from ENCODE phase 4 (v4) or earlier ENCODE releases. b. Schematic depicting the approach used to cluster ChIP-seq co-bound genomic regions into regulatory modules. For each cell line, a list of co-bound regions was derived using overlapping ChIP-seq peaks and organized into regulatory modules using a topic modeling algorithm for comparative analysis. The resulting co-association modules can be assessed for CAP involvement, region composition, and CAP co-association dynamics. c. Heatmap displaying the CAP z-scores for all 270 CAPs (left) along with the indicated regulatory metrics (right) across all co-binding modules identified in HepG2 and K562 cells. Rows and columns are clustered according to Pearson and Euclidean distance, respectively. Module clusters enriched for histone marks representing specific chromatin states are highlighted. d. Bar plots displaying individual CAP z-scores across all modules for the indicated CAPs. Modules are ordered the same as Fig. 1c. Module clusters enriched for chromatin states are as indicated.

To organize CAP peaks into interpretable co-binding groups, we first generated a CAP-region co-binding matrix representing all regions that contain at least three overlapping CAP ChIP-seq peaks, resulting in 174,136 and 146,999 co-bound regions in K562 and HepG2, respectively. Interestingly, despite having a similar distribution of peak numbers and genome coverage across all CAPs, K562 co-bound regions contain fewer CAPs compared to HepG2 (mean CAPs per region: 21.1 (HepG2) and 15.7 (K562), p-value < 2.2 x 10^-16^; **Extended Data Fig. 1d-f**). We then used these co-binding matrices as input into the Regulatory Module Discover (RMD)^16^ pipeline, which implements Bayesian non-parametric topic modeling to systematically cluster the co-bound regions into modules, whose total number is determined based on the complexity of the data **(Fig. 1b)**. Topic modeling^20^ has recently been explored as a tool to represent complex biological data in a low-dimensional setting in studies evaluating CAP co-binding^16^, transcriptional networks^21^, and biomarker discovery^22^. The RMD pipeline is a soft-clustering technique, meaning co-bound regions can be represented by multiple modules **(Fig. 1b)**. This feature allows for the final number of discovered modules to be in the tens as opposed to the thousands produced by hard-clustering self-organizing maps or mixture models. CAP distribution across modules represents their relative abundance at module regions **(Fig. 1b)**. As proof of principle, we utilized RMD on the ENCODE ChIP-seq datasets conducted on A549 cells following treatment with either vehicle control or dexamethasone, a cortisol mimic compound, for 12 hours^23^. Interestingly these results revealed CEBPB and NR3C1 associate with EP300 upon dexamethasone treatment but not in the untreated condition (**Extended Data Fig. 1g-h**), in agreement with existing knowledge that dexamethasone treatment activates NR3C1 and facilitates new enhancer interactions^23^.

After applying RMD to co-bound regions individually in K562 and HepG2 cell lines, we combined the results and generated a global combinatorial binding heatmap map consisting of 70 and 56 CAP co-binding modules, for K562 and HepG2, respectively **(Fig. 1c and Extended Data Table 3)**. In effort to standardize the data across CAPs, the number of regions bound by each module per CAP is displayed as a z-score, which was calculated individually for each cell line (see Methods). These data provide a simplified snapshot of the highly complex genome-wide CAP co-binding associations present across HepG2 and K562 cells.

### Co-association modules unbiasedly identify regulatory context for chromatin states

For each co-binding module, we first examined the relationship between CAP co-occupancy and chromatin regulatory state patterns associated with enhancers, promoters, CTCF, and repressed sites. Interestingly, despite the modules being clustered according to CAP co-occupancy z-scores, we found modules dominated by specific regulatory states grouped together, as determined using histone ChIP-seq signals across module regions. **(Fig. 1c and Extended Data Tables 4-5)**. A similar trend was observed when evaluating module IDEAS^24^ state and cCRE^12^ status distributions across all regions represented in each module **(Extended Data Fig. 2a)**. These results provide functional context to the modules and agree with previous studies that have suggested CAP co-occupancy as an indicator of regulatory state^9,16^. We further investigated modules within promoter, enhancer, transcribed, and CTCF-associated clusters. When evaluating CAP binding at each module, we found the co-occupancy correlation across cell lines to be highest at transcribed modules (Pearson correlation = 0.8, p-value = 2.12e^-8^) and the lowest at enhancer modules (Pearson correlation = 0.02, p-value = 0.8) **(Extended Data Fig. 2b)**. This is consistent with existing knowledge of enhancers representing cell type-specific regulatory elements^4,25^. Indeed, enhancer-associated modules exhibited cell type-specific co-occupancy of known lineage-specific factors including FOXA3 and GATAD2A for HepG2 cells and TCF3 for K562 cells **(Fig. 1c-d and Extended Data Fig. 2b)**. Modules associated with actively-transcribed genes (“Transcribed” IDEAS state) in both cell lines were largely depleted of DNA-binding TFs and instead were comprised of the co-occupancy of POL2 subunits POL2RA and POL2RG and the splicing-associated factors RBFOX2, RBM22, and AGO1, the latter indicative of the tight coupling of transcription and splicing. Meanwhile, promoter modules were comprised of known promoter-associated factors, such as POLR2A, POL2RG, ZFX, and RNA-binding proteins (RBPs) including RBFOX2 and HNRNPLL. Both HepG2 and K562 display cell type-specific CAP co-occupancy at promoter modules. For example, over 30% of HepG2 promoter module regions are bound by the TF KLF16, whereas K562 modules identify KLF16 co-occupancy at enhancer- and CTCF-associated modules **(Fig. 1c-d and Extended Data Fig. 2b)**. In contrast, over 35% of K562 and less than 10% of HepG2 promoter module regions are bound by ATF1 and ATF7 TFs **(Extended Data Fig. 2b-c)**. In agreement, the correlation strength between ATF1 or ATF7 ChIP-seq signal values with POLR2A signal is substantially higher in K562 than HepG2 across all co-bound regions **(Extended Data Fig. 2d)**. Notably, despite this differential occupancy at promoter modules, the well-established co-occupancy of ATF1, ATF7 and other AP-1 family members was identified in a separate module in both cell lines, suggesting the dynamics of TF co-binding include both conserved and cell type-specific patterns **(Fig. 1c-d and Extended Data Fig. 2c-d)**.

### Shared co-binding modules exhibit cell type-specific regulatory patterns

While co-regulatory epigenetic landscapes contain many cell type-specific features, we hypothesized that a subset of CAP co-associations would be conserved across cell types. Using Pearson correlation calculations of CAP z-scores for each HepG2 and K562 module pair, we identified 9 HepG2 co-binding modules that each have a highly comparable module in K562 **(Fig. 1c and Extended Data Table 6)**. For identifying similar modules, we found our strategy of using Pearson correlation of CAP z-scores was superior to simply identifying module pairs with the highest percent of shared CAPs, as Pearson correlation of CAP z-scores was more representative of the various co-binding patterns observed within a module (**Extended Data Fig. 3a**). While no module existed in its entirety in the opposing cell line, these shared regulatory modules (SRMs) contained “core” CAPs that were present across SRM pairs and enriched for CAPS from the same TF family (SRMs: ONR, ZF, CEBP/ATF4, bHLH, and sMAF), RNA-binding proteins (SRMs: pcRBP and POL2/RBP), and readers and writers of the same histone mark (SRM: HetChrom) **(Fig. 2a, Extended Data Fig. 3b, and Extended Data Table 3)**. Importantly, SRMs co-bound by many TFs showed enrichment for a specific TF family while devoid of co-occupancy of other TF family members **(Fig. 2b)**. Given that TFs from the same family recognize similar motifs, it is likely TF family-enriched SRMs represent genomic regions occupied by a subset of these TFs, especially considering that ONR and sMAF TFs bind DNA as dimers, as opposed to all constituents co-binding simultaneously. Furthermore, as our initial investigation found CTCF and transcribed regulatory cell states to contain highly conserved CAP co-association **(Extended Data Fig. 2b)**, it was expected that these modules would also arise as SRMs (CTCF/Cohesin and POL2/RBP, respectively). The complete list of SRM constituents can be found in **Extended Data Tables 3 and 7**.

**Figure 2.**
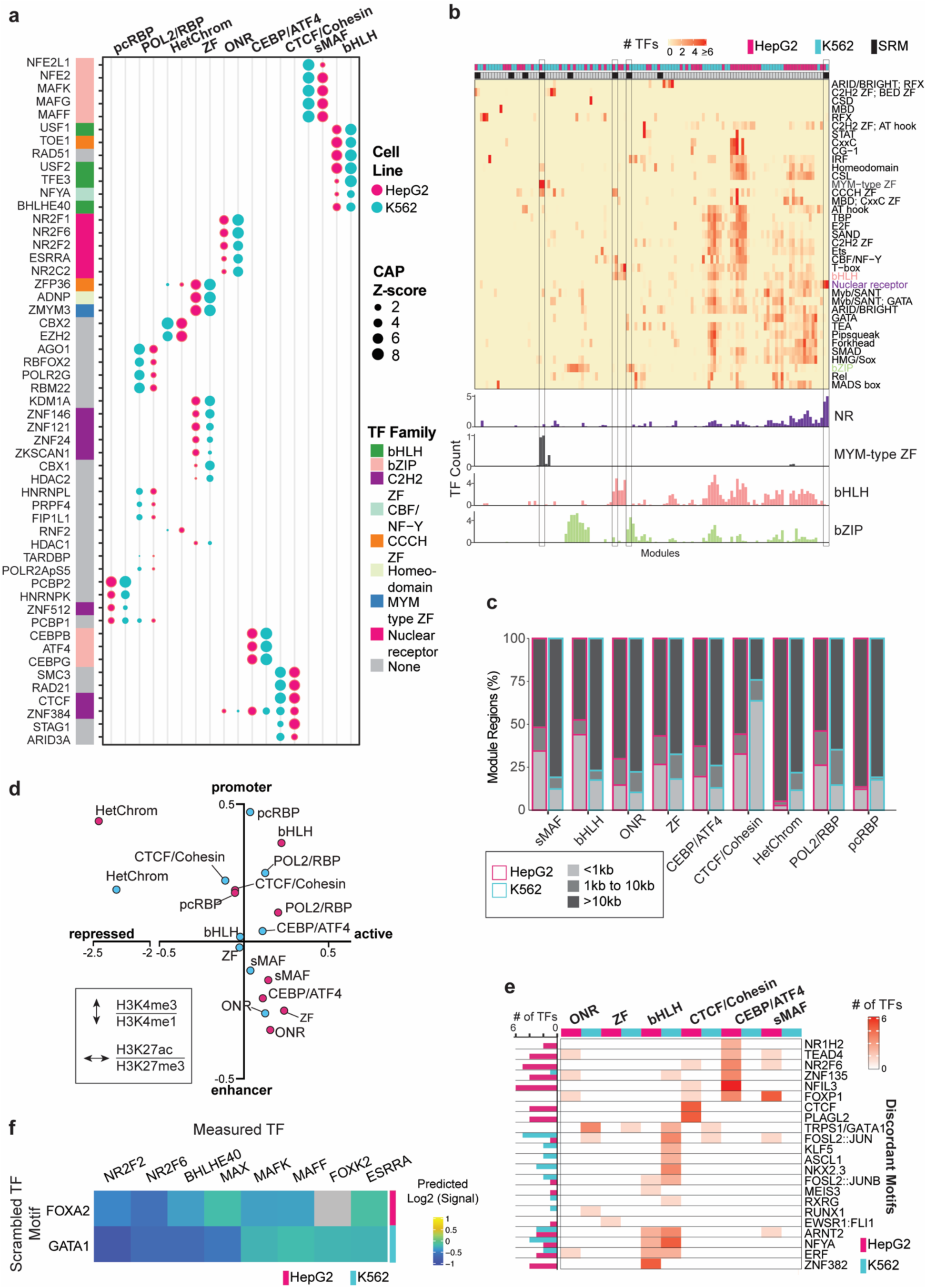
Shared co-binding complexes exhibit cell type-specific regulatory patterns. a. Dot plot displaying the CAP z-scores for the shared CAPs present at common regulatory modules. TF family annotation is indicated. b. Heatmap depicting the number of TFs within each module that belong to each TF family (top) and bar plots displaying the indicated TF family distribution across all modules (bottom). Note the highlighted SRMs are enriched for a single TF family. c. Percent of SRM module regions that are within the indicated distances from corresponding SRM in the opposite cell line. d. Scatterplot summarizing the regulatory state and activity of SRMs in each cell line. The x-axis corresponds to the ratio of H3K27ac to H3K27me3 signals and the y-axis corresponds to the ratio of H3K4me3 to H3K4me1 signals. e. Heatmap displaying the number of significant discordant Tomtom *de novo* motifs discovered from TF ChIP-seq peaks present within regions from each SRM. Discordant motifs were grouped based on their JASPAR motif cluster and the representative motif cluster “seed” motif is indicated as the row name. f. Heatmap depicting the log2 fold change of the predicted signal values at regions with scrambled FOXA2 or GATA1 motifs compared to sequences with in-tact motifs in the indicated cell lines. For each TF pair, co-bound regions where the scrambled TF motif was present were used for the predictive analysis.

We next explored if the regulatory status and genomic position of SRMs was maintained across cell types. Interestingly, despite having similar CAP composition, we found SRMs largely occupy distinct genomic regions, as the nearest corresponding module loci in the opposite cell line is over 10kb away, on average **(Fig. 2c)**. An exception is the CTCF/Cohesin module, where 63% of K562 module regions are within 1kb of HepG2 CTCF/Cohesin module regions and 32% of HepG2 module regions are within 1kb of K562 module regions, consistent with the existing literature suggesting CTCF-Cohesin-bound loops are highly conserved across cell types^26,27^. We then investigated if regions bound by SRMs display comparable regulatory activities. To do this we first calculated the signal z-score of the regulatory histone marks H3K27ac, H3K4me3, H3K4me1, and H3K27me3 at all CAP co-bound regions provided as input for the RMD pipeline. We then calculated the average signal z-score for each histone mark across module regions **(Extended Data Fig. 4a-b)** and used the difference of H3K4me3 to H3K4me1 or H3K27ac to H3K27me3 to determine the module regulatory state and activity, respectively. Active enhancers are bound by sMAF and ONR in both cell lines, while CTCF/Cohesin, POL2/RBP, HetChrom and pcRBP consistently co-occupy promoters **(Fig. 2d)**. In contrast, whereas HepG2 bHLH, CEBP/ATF4, and ZF modules bind promoter or enhancers, the corresponding K562 modules lack a regulatory state preference **(Fig. 2d and Extended Data Fig. 4a-b)**. Given that POL2/RBP SRMs are comprised of CAPs associated with transcription which bind to distinct promoters across cell lines **(Fig. 2a and 2c)**, we hypothesized that these SRMs would regulate cell type-specific genes. Indeed, we found genes regulated by POL2/RBP module in HepG2 showed significantly higher expression in HepG2 compared to K562 cells, while the converse was true for K562 regulated genes **(Extended Data Fig. 4c)**.

Given the low overlap of genomic regions bound by SRMs across cell lines, we hypothesized that SRMs would be associated with different regulatory pathways. Pathway enrichment of SRM-regulated genes showed similar regulatory networks are mediated by each SRM in HepG2 and K562 cells **(Extended Data Fig. 5a and Extended Data Table 8)**. We therefore investigated genes distally regulated by SRMs, identified using promoter capture Hi-C (PCHi-C), in both cell lines. We found a significant overlap between the target genes which interact with distally occupied genomic regions of ONR, bHLH, POL2/RBP, and CTCF/Cohesin SRMs compared to what would be expected by random chance **(Extended Data Fig. 5b-c)**, suggesting that while SRMs largely occupy distinct genomic regions in a cell type-specific manner, a subset of SRM regions maintain distal interactions with the same target genes.

Of note, while SRMs were identified based on their similar CAP composition, SRMs from each cell line also displayed cell type-specific CAPs, termed ancillary CAPs (aCAPs), independent of differential CAP expression levels across cell lines **(Extended Data Fig. 6a)**. For example, the HetChrom SRM contains four “core” CAPs present in both HepG2 and K562, which are the Polycomb-repressive complex subunits CBX2, EZH2, and RNF2, as well as ZFP36. Meanwhile, the HepG2 HetChrom module is also comprised of 15 aCAPs which are absent in K562 HetChrom module **(Extended Data Fig. 6b)**. These include TAF15, RBM39, HNRNPK, HNRNPLL, and FUS, of which all are RBPs, which frequently associated with sites of nascent transcription, and when considered together with the high H3K4me3 and H3K27me3 signal levels, suggest HepG2 HetChrom SRMs represent bivalent facultative chromatin regions **(Fig. 2d and Extended Data Fig. 6b-c)**. Notably, we found the HetChrom SRM was distinct from H3K9me3+ modules, a marker of constitutive heterochromatin, which displayed minimal CAP overlap between cell lines, warranting further investigation of CAP heterogeneity at highly inaccessible genomic regions **(Extended Data Fig. 6d)**. As facultative heterochromatin typically modulate developmental gene expression, we investigated H3K27me3 histone levels at HetChrom loci in embryonic stem cells that were either undifferentiated, differentiated to endoderm cells (liver lineage) or mesoderm cells (blood lineage). We found HetChrom H3K27me3 bimodal peaks identified in HepG2 cells were conserved in embryonic and endoderm but not mesoderm cells whereas K562 HetChrom loci demonstrated unimodal H3K27me3 peaks across all lineages **(Extended Data Fig. 6e)**. Together these results suggest that SRM genomic localization and regulatory activity may vary across cell lines, with aCAPs possibly contributing to cell type-specific SRM characteristics.

### Cell type-specific localization of shared co-binding modules correlates with CAP enrichment at discordant motifs

We next set out to understand what drives SRMs to localize to distinct loci in different cell types. We reasoned that module constituents could occupy different loci due to cell type-specific changes in chromatin accessibility. To test this, we calculated the chromatin accessibility signal for cell type-specific module loci in the opposing cell line. HepG2 SRM genomic regions exhibit significantly reduced K562 ATAC-seq signal compared to the corresponding HepG2 SRM loci **(Extended Data Fig. 7a)**. In contrast, as several K562 SRMs bind to regions with low accessibility in K562 cells, we observed comparable accessibility levels at corresponding HepG2 SRM regions **(Extended Data Fig. 7a)**. These results suggest that chromatin accessibility may play a role in differential module binding in HepG2 cells, but it does not fully explain cell type-specific localization of SRMs.

We next considered if common module CAP co-occupancy is facilitated through the co-association with other DNA-binding TFs in a cell type-specific manner. To investigate this, we assessed *de novo* motif enrichment of DNA-binding TFs present in SRMs across cell type-specific module regions and used peak centrality and enrichment scores to identify a set of high-confidence motifs for each TF. We compared these motifs to the JASPAR motif database^28^, kept motifs with high similarity to a known JASPAR motif, and grouped annotated motifs according to their JASPAR motif cluster to eliminate redundant motif calls. As expected, several enriched motifs matched the known TF motif **(Extended Data Fig. 7b)**. We reasoned that CAPs binding to loci enriched for discordant motifs, defined as known TF motifs that do not match either the canonical TF motif or a similar TF family motif, may explain cell type-specific SRM colocalization. Of the 28 and 35 DNA-binding TFs present in common modules, 14 and 28 TFs were enriched for discordant motifs in K562 and HepG2 cells, respectively, in addition to canonical or similar motifs **(Fig. 2e and Extended Data Fig. 7b-c)**. These cell type-specific motifs may be due to TFs indirectly binding DNA through recruitment via another TF which binds the discordant motif. Across all K562 shared modules, the most frequent discordant motifs discovered at common modules belonged to FOSL2::JUN (JASPAR cluster 1), and TRPS1/GATA1 (JASPAR cluster 11) motif identifiers. In contrast, discordant motifs identified in several HepG2 shared modules were enriched for ZNF135 (JASPAR cluster 65), FOXP1 (JASPAR cluster 10), NR2F6 (JASPAR cluster 4), and TEAD4 (JASPAR cluster 2) **(Fig. 2e)**. Therefore, at SRM loci, TFs are enriched for discordant motifs in a cell type-specific manner.

We further explored this finding through investigating the ONR module in detail. The ONR SRM is comprised of the core TFs, NR2F1, NR2F2, NR2F6, NR2C2, ESRRA, and ZNF384 in both HepG2 and K562 cells. Through our de novo motif discovery at peak sites within ONR module regions we found all 6 TFs to be enriched for motifs similar to their own motif **(Extended Data Fig. 8a)**. Additionally we found JASPAR clusters defined by TEAD4, FOXP1, ERF and ZNF135 motifs or TRPS1, FOSL1::JUN, and RUNX1 motifs as being enriched for 3/6 or 4/6 of the TFs at HepG2 and K562 peaks, respectively **(Extended Data Fig. 8a)**. Motif enrichment of ONR SRM regions co-bound in both cell lines or exclusively bound in one cell line revealed that regions bound in both cell lines were highly enriched for ONR motifs **(Extended Data Fig. 8b)**. In comparison, exclusively bound module regions were enriched for lineage-specific pioneer TFs^29–31^, with liver lineage FOX motifs enriched for HepG2-specific loci and hematopoietic lineage GATA motifs at K562-specific loci **(Extended Data Fig. 8b)**. Additionally, analysis of NR2C2 peak summit positions show enrichment at FOX or GATA motifs for HepG2 and K562 cells, respectively, in addition to TF-matching NR motifs **(Extended Data Fig. 8c-d)**. Moreover, FOXA3, an aCAP, displayed increased binding at HepG2-specific loci compared to shared module loci, whereas GATA2 showed increased binding at K562-specific loci compared to shared module loci **(Extended Data Fig. 8e)**. Finally, we conducted in silico TF cooperative analysis using neural network models of TF binding to generate ONR core and ancillary TF signal predictions following motif perturbation of GATA1 or FOXA3 for K562 and HepG2 cells, respectively (see Methods and **Fig. 6**). We found NR2F2 and NR2F6 to show significant dependency on GATA1 and FOXA3 motif presence in K562 and HepG2 cells, respectively, further supporting that these TFs bind indirectly to DNA via GATA and FOX TFs **(Fig. 2f)**. Together these data strongly suggests that loci consistently bound by members of the ONR module across cell lines is driven by direct DNA-recruitment to the NR motifs while recruitment to uniquely bound loci is facilitated, in part, by cell lineage TFs.

### Dynamics of CTCF-associated shared regulatory modules

HepG2 and K562 CTCF/Cohesin SRMs displayed high co-occupancy of CTCF with cohesin, a multi-subunit protein complex whose constituents include SMC1/3, RAD21, and STAG1/2^32^ **(Figs. 2a, 3a and Extended Data Fig. 2b)**. CTCF is a key regulator of chromatin architecture which interacts extensively with cohesin at loop anchors and topologically associating domain (TAD) boundaries, orchestrating DNA looping and facilitating distal interactions critical for gene regulation^33–37^. We found CTCF/Cohesin SRM regions to be comprised of ∼65%, ∼25%, and 4-14% of high-throughput chromosome conformation capture (Hi-C) loop anchors, intra-loop regions, and TAD boundaries, respectively, in both cell lines **(Fig. 3b)**. This accounted for 45% and 54% of total loops and 42% and 30% of total TADs present in HepG2 and K562, respectively **(Extended Data Fig. 9a-b).** Given the large number of chromatin loops represented by these module regions, we sought to determine how the other co-bound CAPs were distributed amongst Hi-C loops. Surprisingly, all co-bound CAPs, including core CAPs ARID3A and ZNF384, showed significantly elevated signal intensities at loop and TAD boundaries compared to intra-loop or -TAD regions **(Fig. 3c and Extended Data Fig. 9c)**. ARID3A is an ARID family TF that has been shown to promote HCC cell proliferation^38^ and stem cell gene expression^38–40^, whereas ZNF384 is a poorly characterized C2H2-type ZF TF that is primarily studied in the context of a fusion gene product in acute lymphoblastic leukemia^41^. While ARID family members such as ARID1/2 (SWI/SNF subunits) and JARID1 (histone demethylases) are well-known for chromatin remodeling, little is known about the role of ARID3A or ZNF384 in chromatin organization. We found that at module-bound loop anchors, 75% of ARID3A sites were positioned within 49 and 63 bp of CTCF summits, while 75% of ZNF384 sites fell within 54 and 81 bp in K562 and HepG2 cells, respectively, highlighting their close association with CTCF at loop anchors in both cell lines (**Fig. 3d-g**). Furthermore, ARID3A and ZNF384 displayed modest enrichment at cell type-specific chromatin loop ends compared to conserved loops **(Fig. 3h)**. All together, these data identify ARID3A and ZNF384 as novel CAPs which co-occupy a subset of CTCF- and cohesion-bound loop ends in a cell type-independent manner **(Fig. 3i)**.

**Figure 3.**
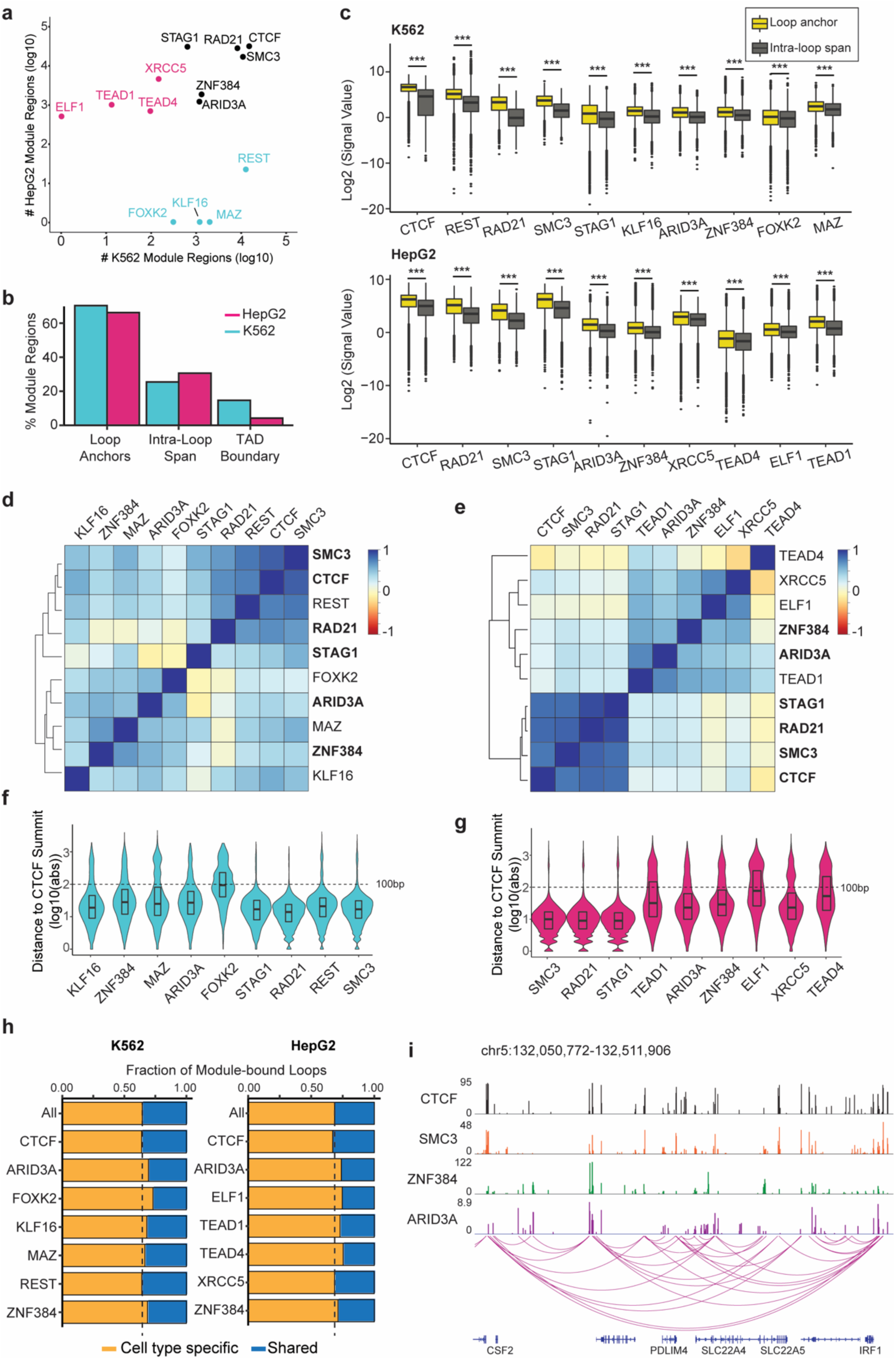
Dynamics of CTCF/Cohesin-associated co-binding modules. a) Scatterplot indicating the number of module regions bound by the indicated CAPs at Hepg2 and K562 CTCF/Cohesin SRMs (HepG2 module 44 and K562 module 2). CAPs are colored according to shared status (black = core, pink = HepG2 ancillary, blue = K562 ancillary). b) Bar plot displaying the number of CTCF/Cohesin SRM regions overlapping Hi-C loop anchors, intra-loop regions, or TAD boundaries in each cell line. c) Boxplots displaying the ChIP-seq signal value for all CAPs constituting K562 (top) or HepG2 (bottom) CTCF/Cohesin SRMs at module bound loop anchors or intra-loop regions. *** = p < 0.0005. d-e) Heatmap depicting ChIP-seq signal Spearman correlations at module-bound loop anchors for all CAPs co-associated in K562 (d) or HepG2 (e) CTCF/Cohesin SRMs. Core CAP names (present in both cell line SRMs) are bolded. f-g) Violin plots displaying the distance to CTCF peak summits for the indicated CAPs at module-bound loop anchors in K562 (f) or HepG2 (g) CTCF/Cohesin SRMs. h) Bar plot displaying the fraction of CAP-bound module loop anchors that are shared cross HepG2 and K562 cell lines or cell type-specific. Shared loops required both loop ends to be conserved between cell lines. All refers to all module-bound Hi-C loops. i) Snapshot of genomic region depicting Hi-C loops where loop anchors are co-bound by CTCF, SMC3, ZNF384, and ARID3A in K562 cells.

Ancillary CAPs also exhibited strong ChIP-seq correlations with CTCF and cohesin at module-bound loop anchors in both cell lines **(Fig. 3d-e)**. Notably, in K562 cells, MAZ and KLF16 bound in close proximity to CTCF at loop anchors and showed binding enrichment at H3K27ac+ (“active”) loop anchors compared to total loop anchors (**Fig. 3f and Extended Data Fig. 9d**). This is consistent with previous reports that identify MAZ as a cofactor of CTCF which regulates chromatin contacts within topologically associated domains (TADs) and modifies gene expression^42,43^. In agreement with previous studies, aggregate peak analysis (APA) illustrated higher contact strength at active loops with MAZ at bound compared to loops lacking either TF **(Extended Data Fig. 9e).** Furthermore, active K562 loops containing KLF16, a zinc finger TF previously unknown to associate with CTCF-bound regions, also exhibited higher contact strength than active loops without KLF16 bound **(Extended Data Fig. 9e)**. In HepG2 cells, TEAD1, a TEA family TF transactivated by the oncogenic YAP protein, exhibited a strong correlation with CTCF ChIP-seq signal levels, bound in close proximity to CTCF at loop anchors, and was enriched at active loops compared to all HepG2 Hi-C loops **(Fig. 3e, 3g, and Extended Data Fig. 9d and 9f)**. These results position core TFs ARID3A, ZNF384, and cell type-specific ancillary CAPs like MAZ, KLF16, and TEAD1 as potential components mediating the structural organization or regulatory activity at chromatin loops.

### CTCF-associated modules represent co-occupied regions of differing CAP compositions

Through our analysis thus far we unexpectedly found co-binding modules with similar CAP composition (SRMs) largely bound different genomic regions between cell lines **(Fig. 2)**. This led us to investigate co-binding modules which occupied the same genomic regions but were comprised of different CAPs. By comparing the both Pearson distance (measure of CAP similarity) and fraction of overlapping regions between HepG2 and K562 module pairs, we identified 4 differentially-bound module (DBMs) pairs **(Fig. 4a)**. Importantly, Pearson distance was not dependent on number of CAPs within modules, ensuring that our approach did not simply just identify modules with low numbers of CAPs **(Fig. 4a)**. Surprisingly, all DBM pairs included the HepG2 CTCF/Cohesin SRM (module 44(H)) paired with a distinct K562 module (**Fig. 4a)**. We found K562 DBMs were typically co-bound by many more CAPs and specifically enriched for TFs compared to HepG2 DBM **(Fig. 4b and Extended Data Fig. 10a-b)**. CTCF-bound peaks displayed comparable signal levels across K562 DBMs **(Extended Data Fig. 10c)**. Furthermore, DBM overlapping regions showed increased chromatin accessibility in K562 but not HepG2 cells compared all ATAC-seq peaks **(Fig. 4c)**. Due to the abundance of TFs present at the CTCF-associated K562 DBMs and lack thereof at HepG2 DBM regions, we hypothesized that these regions represent chromatin loop anchors which are involved in gene regulation in K562 cells. We found target genes regulated by interaction with distal regions in K562, identified through promoter-capture Hi-C, exhibited significantly higher expression in K562 cells compared to HepG2 cells whereas this pattern was not observed for HepG2 distally-regulated target genes **(Extended Data Fig. 10d)**. Furthermore, genes whose promoters overlapped DBM regions were found to have comparable expression levels between cell lines **(Extended Data Fig. 10d)**. We, therefore, hypothesized that DBM regions represent chromatin loops that function as either regulatory loops or structural loops in different cell lines. To investigate this, we performed PCHi-C) in both HepG2 and K562 cells, allowing us to generate a list of high confidence regulatory loops connecting distal loci to promoter regions. We then compared loop interaction scores at DBM distally regulated genes and all loop loci using PCHi-C and Hi-C datasets, the latter which identifies all chromatin loops regardless of function. We found PCHi-C loop interaction scores were significantly increased at K562 DBM regions compared to all interacting loops whereas HepG2 DBM regions showed no significant difference **(Fig. 4d)**. In contrast, HepG2 Hi-C interaction scores were significantly higher in DBM regions compared to all loops whereas K562 DBM Hi-C interaction scores were significantly reduced compared to all loops **(Fig. 4e)**. We next investigated CAP co-occupancy patterns at PCHi-C distal ends in K562 DBMs by evaluating the ChIP-seq signal correlations for all CAPs. We found many CAPs displayed a strong binding correlation at these regulatory distal ends **(Fig. 4f)**. For example, CAPs such as JUN, JUND, RCOR1 and TBL1XR1 all showed a high correlation with EP300, a CAP commonly associated with active enhancers **(Fig. 4f-g)**. Together this demonstrates that highly overlapping differentially bound modules are associated with CTCF chromatin loops and likely exhibit different regulatory roles based on cell type-specific co-binding differences which lead to differential regulation of distal gene targets.

**Figure 4.**
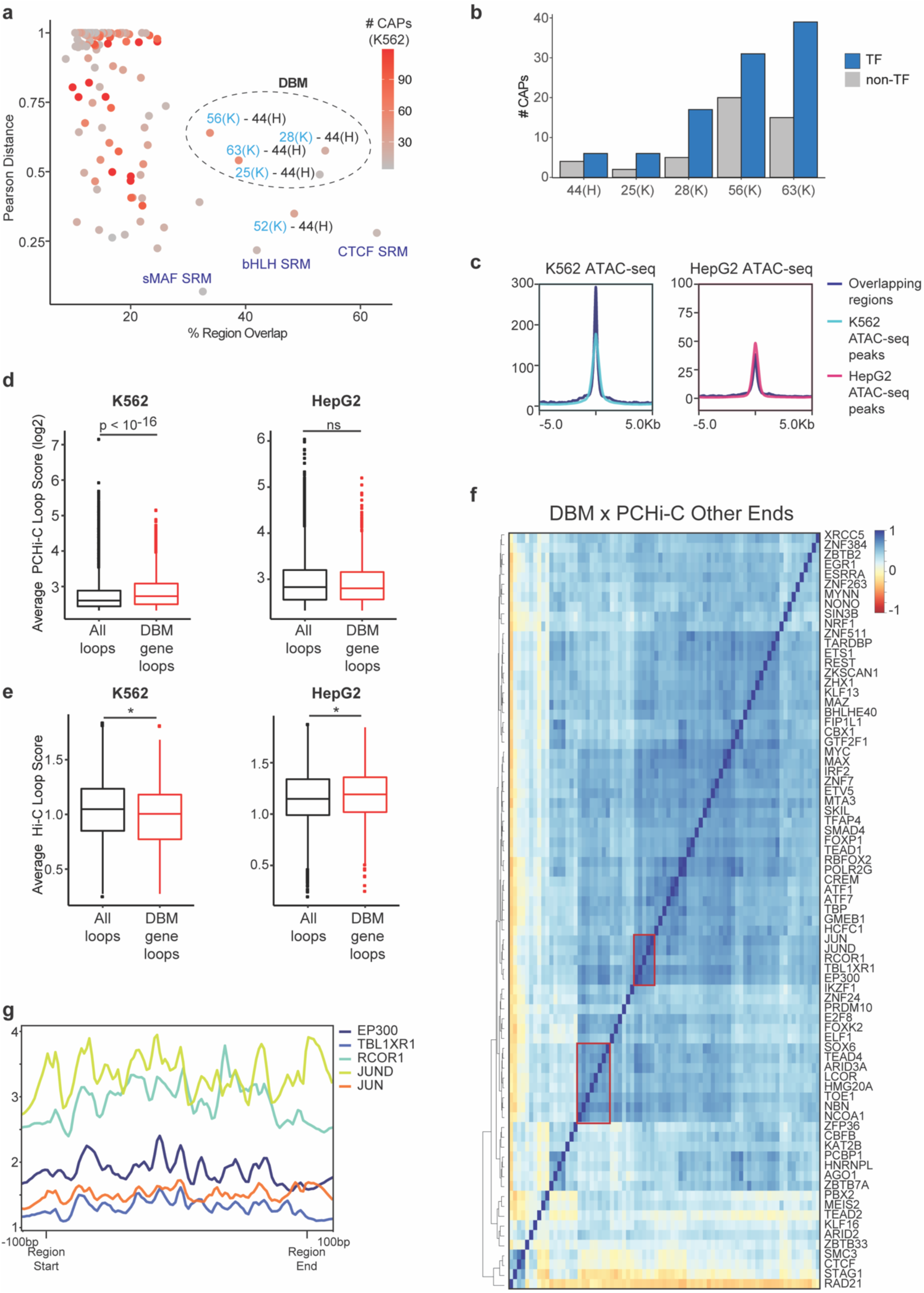
Module regions with differing CAP compositions across cell lines. a) Scatterplot depicting the strategy to determine differentially bound modules (DBMs). Pearson distance and fraction of region overlap was calculated for all possible module pairs. For each module, the module pair with the highest region overlap was kept and plotted. Module pairs with high Pearson distance metric (low CAP similarity) and high region overlap were classified as DBMs (circled). Points are colored based on the number of TFs present in the K562 module for each pair. Modules are referred to by their ID number followed by H or K for HepG2 and K562, respectively. b) Bar plot displaying the fraction of CAPs that are TFs and non-TFs for each DBM. Modules are referred to by their ID number followed by H or K for HepG2 and K562, respectively. c) K562 or HepG2 ATAC-seq signal profile plots at regions DBM overlapping or cell type-specific regions. d-e) PCHi-C (d) or Hi-C (e) interaction scores at all cell line loops or loops with genes distally regulated by DBM-bound other ends in K562 and HepG2 cells. * = p < 0.05; ns = not significant. All p*-*values determined using Wilcoxon rank sum test. f) Heatmap depicting ChIP-seq signal Spearman correlations at K562 DBM-bound PCHi-C “other” ends for all CAPs present in K562 DBMs. Specific CAP complexes with high correlations are highlighted. g) Profile plot of the indicated highly correlated CAPs across PCHi-C “other” end regions. PCHi-C regions are scaled to represent the same genomic distance.

### Co-binding module interactions at 3D chromatin regulatory loops

We next investigated the frequency of co-binding module combinations that are brought together by chromatin loops to regulate transcription. To do this, we used our PCHi-C datasets in both HepG2 and K562 cells which generated a list of high confidence regulatory loops connecting distal loci to promoter regions. We then conducted permutation testing to statistically test whether associations between two co-binding modules at interacting PCHi-C loops occur significantly more than expected by chance **(Fig. 5a)**. Surprisingly, in both HepG2 and K562 cells, over 50% of regulatory modules showed significant interactions with at least 20 other regulatory modules **(Extended Data Fig. 11a-b)**, suggesting there is little specificity among the majority of interactions between regulatory modules.

**Figure 5.**
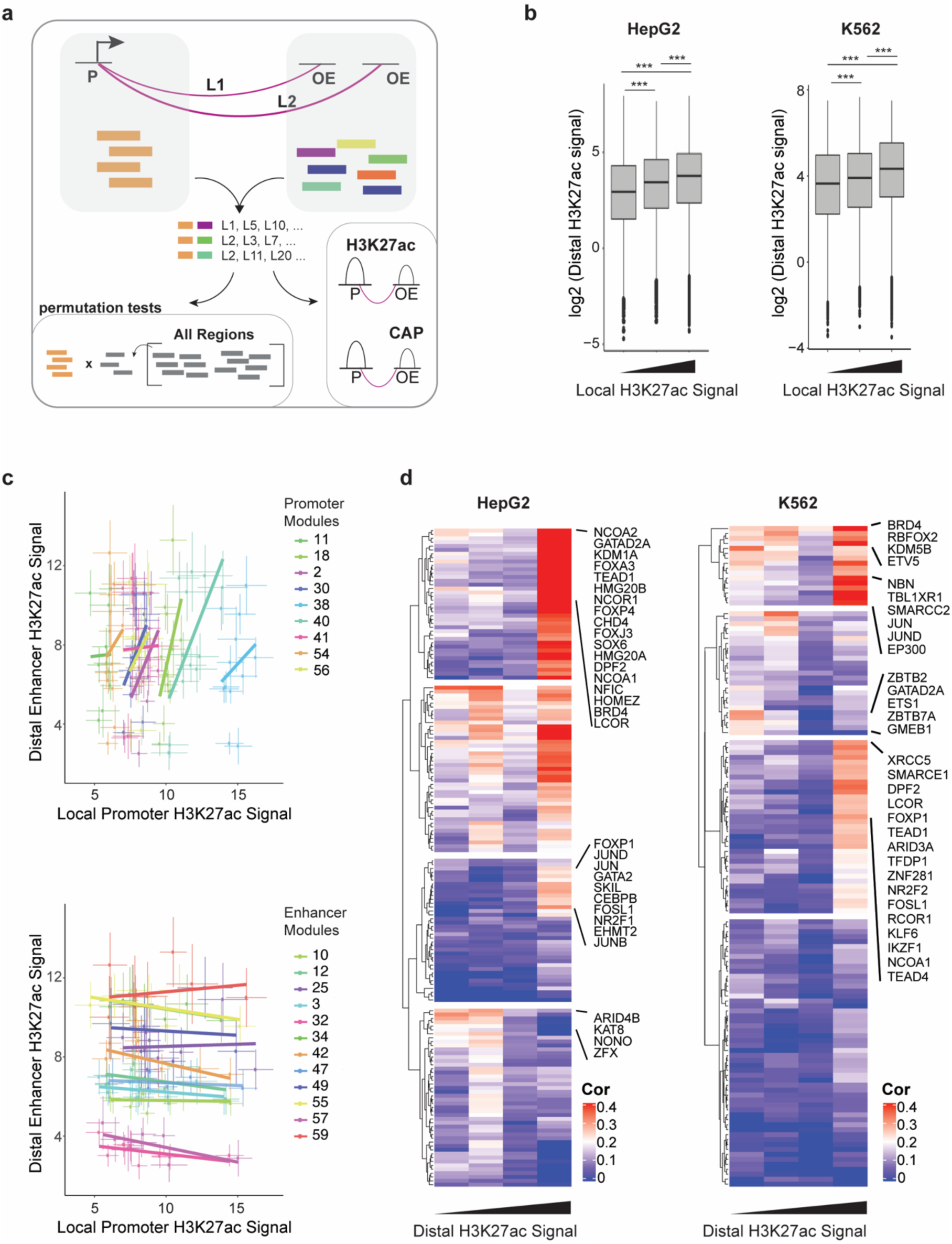
Module-module interactions at regulatory regions. a) Schematic describing analyses integrating PCHi-C data and regulatory modules. PCHi-C data was used to identify module-module interactions that occur more than expected by chance, carried out using permutation testing. For interacting loops (L), H3K27ac and CAP signal were assessed at promoter (P) and other end (OE) DNA regions. Color blocks refer to individual regulatory module regions. b) H3K27ac signal at enhancer-associated distal PCHi-C regions binned according to H3K27ac signal at promoter-associated bait regions. *** = p < 0.0005. All p-values determined using Wilcoxon rank sum test. c) Scatterplot of the HepG2 H3K27ac signal value at bait/promoter regions (x axis) and other end/distal regions (y-axis) for interacting module pairs. Lines indicate standard error values. Module interactions and linear trend lines are colored by promoter-associated modules (top) or enhancer-associated modules (bottom). d) CAP signal correlation with H3K27ac signal at enhancer-associated distal regions. Distal regions were binned according to H3K27ac average signal quartiles. Only CAPs present at enhancer-associated modules were investigated. CAPs known to be members of chromatin remodeling complexes are bolded and co-activators are indicated in red.

In an effort to explore the regulatory impact of module-module interactions, we intersected the PCHi-C loop calls with H3K27ac signal and compared H3K27ac signal levels interacting loop ends **(Fig. 5a)**. We then evaluated the local H3K27ac signal at module-bound promoter “bait” loop ends with the corresponding interacting distal module-bound loop ends amongst promoter-associated and enhancer-associated modules defined in **Figure 1**. Within individual promoter-associated modules, we found the local H3K27ac signal to be higher when interacting with a distal enhancer module that also displayed elevated H3K27ac signal, consistent with current models suggesting enhancer activity directly modulates promoter transcription levels^44^ **(Fig. 5b-c and Extended Data Fig. 12a-b)**. In contrast, when considering individual enhancer modules, we did not observe a strong correlation between H3K27ac signal at local and distal interacting module regions. Since individual enhancer modules appeared to correlate highly with distal H3K27ac signal, we next asked what CAPs are driving the activity at distal PCHi-C regions. To do this, we binned distal interactions with promoter-associated modules according to H3K27ac signal and then calculated the correlation between CAP signal and H3K27ac signal across all regions within each bin **(Fig. 5d)**. Despite the divergent CAPs that make up the enhancer-associated modules **(Extended Data Fig. 2b)**, we found similar CAPs are associated with high H3K27ac activity at promoter-enhancer module PCHi-C interactions. For example, we found chromatin remodeling components exhibited high signal to H3K7ac correlation at highly active distal regions in both HepG2 and K562 cell lines. This included mammalian SWI/SNF subunits SMARCE1, SMARCC2, and DPF, CoREST constituents HMG20A/B, PHF21A, and RCOR1, Polycomb repressor complex 2 members LCOR, EZH2, and KDM5B, and the nucleosome remodeling and deacetylase (NuRD) complex members KDM1A, CHD4, and GATAD2A. Beyond chromatin remodelers, TFs known to localize to enhancer regions such as FOXP1, TEAD1, TEAD4, JUN, JUND, and FOSL1 as well as TF co-activators such as BRD4 displayed high correlation with H3K27ac signal only at high activity distal regions in both cell lines **(Fig. 5d)**. Co-occupancy of chromatin remodeling complexes, co-repressor complexes, co-activators, and TFs suggest complex regulation of distal regions bound by enhancer modules. Together, these data suggest that promoter module activity appears to correlate with distal enhancer activity, whereas enhancers display more autonomous activity driven by chromatin remodeling complexes and CAPs with both co-activation and co-repressor activities.

### Co-binding modules as a means to identify TF-TF combinatorial DNA binding pairs

We then sought to utilize the co-binding modules to identify candidate combinatoric TF pairs. To do this we used our co-binding modules to filter the 18,145 total possible TF pairs into 4,684 potential TF pairs based on their presence in the same module **(Fig. 6a)**. As binding redundancy may exist among TFs that bind to the same motif, we removed all TF pairs where the TF motifs were highly similar using the motif clustering database provided by Viestra et al.^45^, resulting in 3,820 and 3,611 candidate TF pairs in HepG2 and K562 cells, respectively. For each TF pair, we then calculated the signal correlations at co-bound loci and removed any TF pairs with negative correlations **(Extended Data Fig. 13a)**. This excluded 664 (17.4%) and 723 (20%) HepG2 and K562 candidate TF pairs, respectively, which likely reflect TFs that are anti-cooperative. Of the remaining filtered TF pairs, 1,468 were identified in both cell lines whereas 1,688 and 1,406 TF pairs were HepG2 and K562, respectively, specific due to either cell type-specific module co-occupancy or peak signal positive correlations **(Extended Data Fig. 13b)**. Out of these filtered pairs, 9.4% are previously known to interact via protein-protein interactions (PPIs) whereas the remaining 90.6% have no known physical associations **(Extended Data Fig. 13c)**. This first-pass analysis created a list of candidate combinatoric TF pairs that show highly correlated binding strength at co-bound regions.

In an effort to validate these candidate TF combinatoric pairs *in silico*, we used base-resolution deep learning models of TF binding derived from ENCODE ChIP-seq data^17,46,47^ **(Extended Data Table 9)**. These TF prediction models can predict the local TF binding profile shape and signal across input DNA sequence. To explore the cooperative effects of TF_A_ and TF_B_, we compared TF binding signal predictions at unmodified or motif perturbed loci to infer TF dependency and cooperativity. To do this we retrieved all co-bound DNA sequences where both Motif_A_ and Motif_B_ are present and generated three conditions from these sequences: 1) no modifications to the sequences (control sequences), 2) Motif_B_ is scrambled, and 3) Motif_A_ is scrambled **(Fig. 6a)**. We provided these 3 conditions as input into the trained TF binding prediction models (listed in **Extended Data Table 9**) and compared the predicted signal profile at sequences lacking the cooperative motif to the control sequences. For example, to identify if TF_A_ and TF_B_ exhibit cooperativity, we measured the predicted TF_A_ profile at sequences with a scrambled Motif_B_ compared to control sequences, and we measured the predicted TF_B_ profile at sequences with a scrambled Motif_A_ compared to control sequences **(Fig. 6a)**. Any identified significant TF pair was investigated in the other cell line regardless of module co-occupancy or signal correlation status. Of the 4,878 total TF pairs investigated using prediction models with passing quality metrics, 14% (702 TF pairs) were predicted to cooperate significantly in at least one cell line **(Fig. 6b and Extended Data Fig. 13d)**. Interestingly, 60 HepG2 and 37 K562 pairs showed bidirectional cooperation (i.e. Motif_A_ impacts TF_B_ binding and Motif_B_ impacts TF_A_ binding) **(Extended Data Fig. 13e-f)**. TF pairs with similar motifs but members of different motif clusters^45^ served as positive controls validating this approach. For example, ATF2 and JUN both exhibit a significant drop in predicted TF signal when the other’s motif is scrambled, which is expected as their motifs differ by one nucleotide^48^ **(Extended Data Fig. 13e)**. To provide a more stringent cutoff criteria, we excluded all TF pairs with the same DNA binding domain (DBD) from future analyses. Out of the 592 significant TF pairs with differing DBDs, 537 (91%) have no previously known physical PPI **(Fig. 6b)**. Importantly, we identified novel TF pairs with bidirectional (motif scrambling of either motif impacted the opposite’s predicted binding signal) and unidirectional (motif scrambling of only one motif impacted the opposite’s predicted binding signal) cooperative features **(Extended Data Fig. 13e-g)**. Using this approach on HepG2 prediction models identified several novel TF-TF cooperators including the liver pioneer TF FOXA1 to cooperate bidirectionally with CEBPG and unidirectionally with TCF7 and TEAD1 **(Fig. 6b, Extended Data Fig. 13g)**. Meanwhile, analysis of K562 datasets discovered GATA2 to cooperate with MAFF, MAFK, and FOSL1 **(Fig. 6b and Extended Data Fig. 13g)**. Additionally, both HepG2 and K562 showed USF1/2 dependencies on ATF3 and CREB3 for binding **(Extended Data Fig. 13g)**. A complete list of all significant cooperative TFs can be found in **Extended Data Table 10**. Together these data compile a list of TF pairs with high-confidence cooperative DNA binding, mediated through their own motifs.

We then further investigated which TF family motifs displayed high cooperation with other TFs. We found TFs which are members of the AP1 and CREB TF motif families had high cooperation in both cell lines, suggesting that the presence of an AP1 and/or CREB TF assists with recruitment of additional TFs across cell types **(Extended Data Fig. 14a-b)**. Additionally, FOX/4 TF motifs, which includes FOXA TFs, exhibited the largest number of cooperative TFs in HepG2 cells, consistent with their role as pioneer TFs specifically in hepatic lineages. In contrast, we found TFs in K562 cells were highly dependent on MAF and ETS/1 motifs, with over 30 different TFs displaying *in silico* dependency on MAF or ETS/1 motif presence for binding. In both cases, the TFs dependent on FOX/4 or MAF motifs included a diverse array of TFs across several DNA-binding domain TF families **(Extended Data Fig. 14c)**. Among the TFs dependent on intact FOX motifs, TEAD1 and TEAD4 exhibited significant reductions in predicted binding at sequences with FOX motif perturbation in HepG2 but not K562 cells **(Fig. 6c-d)**. Interestingly, the TEAD1 DNA footprint, which is present in control sequences, is absent when FOXA3 motif is scrambled in HepG2 cells, suggesting that at these TEAD1-FOXA3 co-bound sequences, either TEAD1 is binding DNA indirectly via FOXA3 or FOXA3 pioneer properties are important for TEAD1 binding **(Fig. 6c)**. TEAD TFs have been demonstrated to cooperate with liver TFs FOXA3 and HNF4a during embryonic development^49–51^. Our results using deep learning base-resolution ChIP-seq prediction models suggest that this cooperation may continue in hepatocellular carcinoma cell lines, such as HepG2. Together, we demonstrated how utilizing machine learning models of TF co-binding in combination TF base pair binding can be used to identify high confidence TF cooperations.

**Figure 6.**
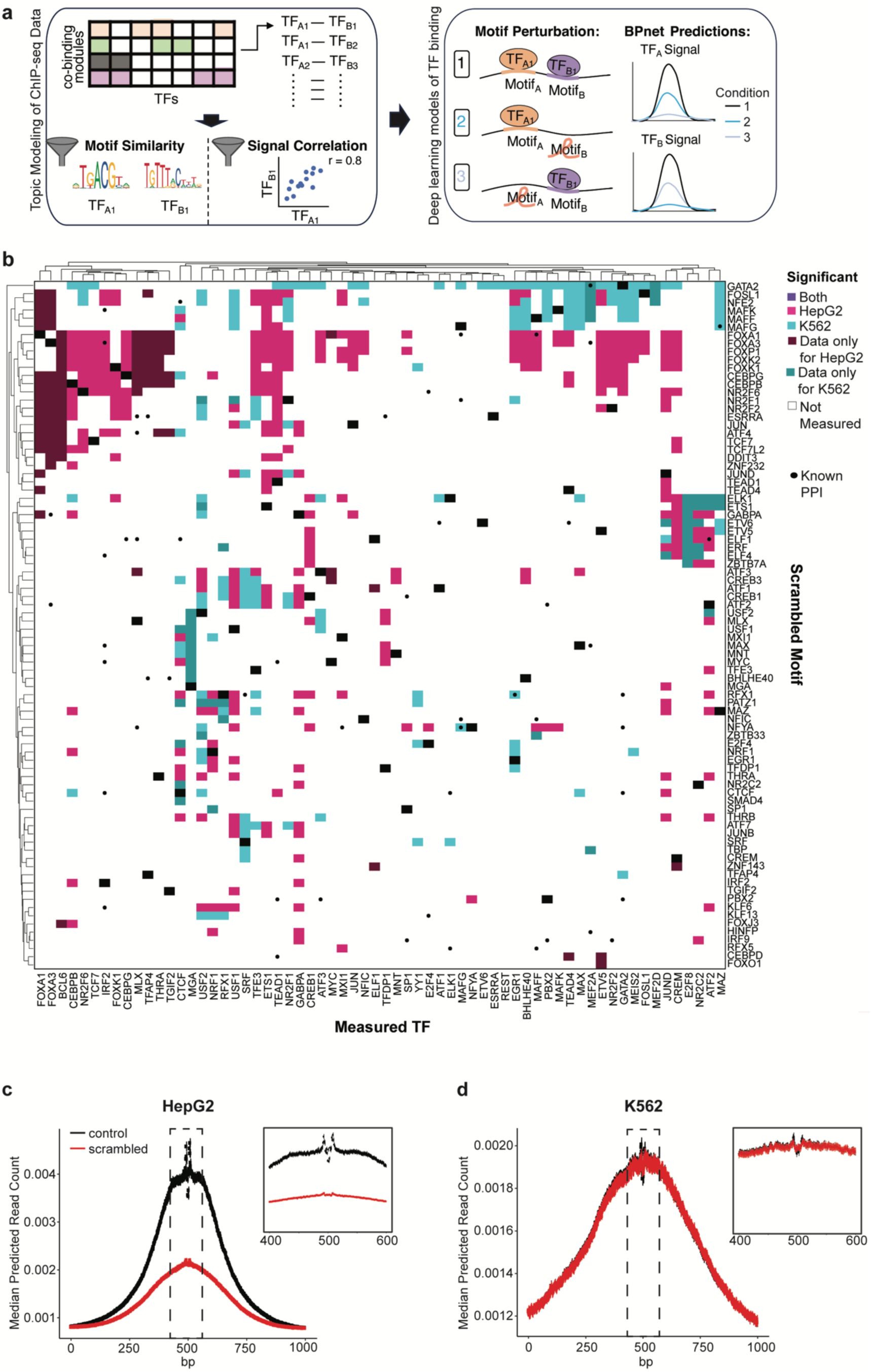
Assessment of candidate protein-protein cooperation using regulatory modules and base pair model of TF binding. a) Schematic depicting the strategy to identify TF cooperation. TF pairs were identified using results from co-binding modules and filtered for TFs with differing motifs and positively correlative ChIP-seq binding signals at co-bound regions. For each TF pair, regions with overlapping peaks that contained both TF motifs were either unmodified (control sequences) or modified to scramble either TF motif. Control and perturbed sequences were provided as input to base pair resolution models which predicted TF signal across all input sequences. b) Heatmap depicting the significant (see Methods) TF pairs identified from base pair deep learning prediction models colored according to significance in one or both cell lines. As co-binding module compositions was a prerequisite for this analysis, not all TF pairs are analyzed. When data for a specific cell line is missing, it is either because the model was unavailable or it failed quality control. c-d) Predicted signal plots of TEAD1 ChIP-seq at control or FOXA3 motif scrambled sequences in HepG2 (c) or K562 (d) models. The highlighted region is shown in magnification on the top right of each plot. Note in (c) the TEAD1 footprint is absent in FOXA3 scrambled sequences compared to control sequences, suggesting that TEAD1 is binding DNA indirectly through FOXA3.

### HOT site characteristics across cell lines

Genomic loci bound by up to hundreds of CAPs, termed high occupancy target (HOT) sites, have been shown by us and others to functionally regulate the transcription of highly expressed genes through both cell type-specific and ubiquitous mechanisms^9–11,52^. Using ChIP-seq CAP datasets available in both HepG2 and K562 cells that passed quality metrics (see Methods), we identified 9,353 and 6,538 HOT sites, respectively. Notably, this substantial difference in HOT site number persisted when using all available ENCODE ChIP-seq data, with 9,590 HepG2 and 5,461 K562 HOT sites, suggesting it is not due to the specific datasets used in the analysis. Moreover, regulatory modules did not show substantial enrichment for HOT sites **(Extended Data Fig. 15a)**, suggesting comparison of HOT sites should be performed directly. Out of the HOT sites identified from common ChIP-seq datasets, 39.4% of HepG2 and 67% of K562 HOT sites overlapped between the two cell types. Despite the difference in number of HOT sites, the distribution of HOT sites across cCREs does not differ substantially between cell types (**Fig. 7a).** This was found to be true independent of HOT site cutoff and TF set used for HOT site identification (**Extended Data Fig. 15b**). Interestingly, shared HOT sites were largely promoter-like (PLS) and proximal enhancer-like (pELS) regions, whereas cell type-specific HOT sites were dominated by distal enhancer-like (dELS) sequences, demonstrating that cell type-specific regulation of enhancer regions extends to HOT sites (**Extended Data Fig. 15c)**.

**Figure 7.**
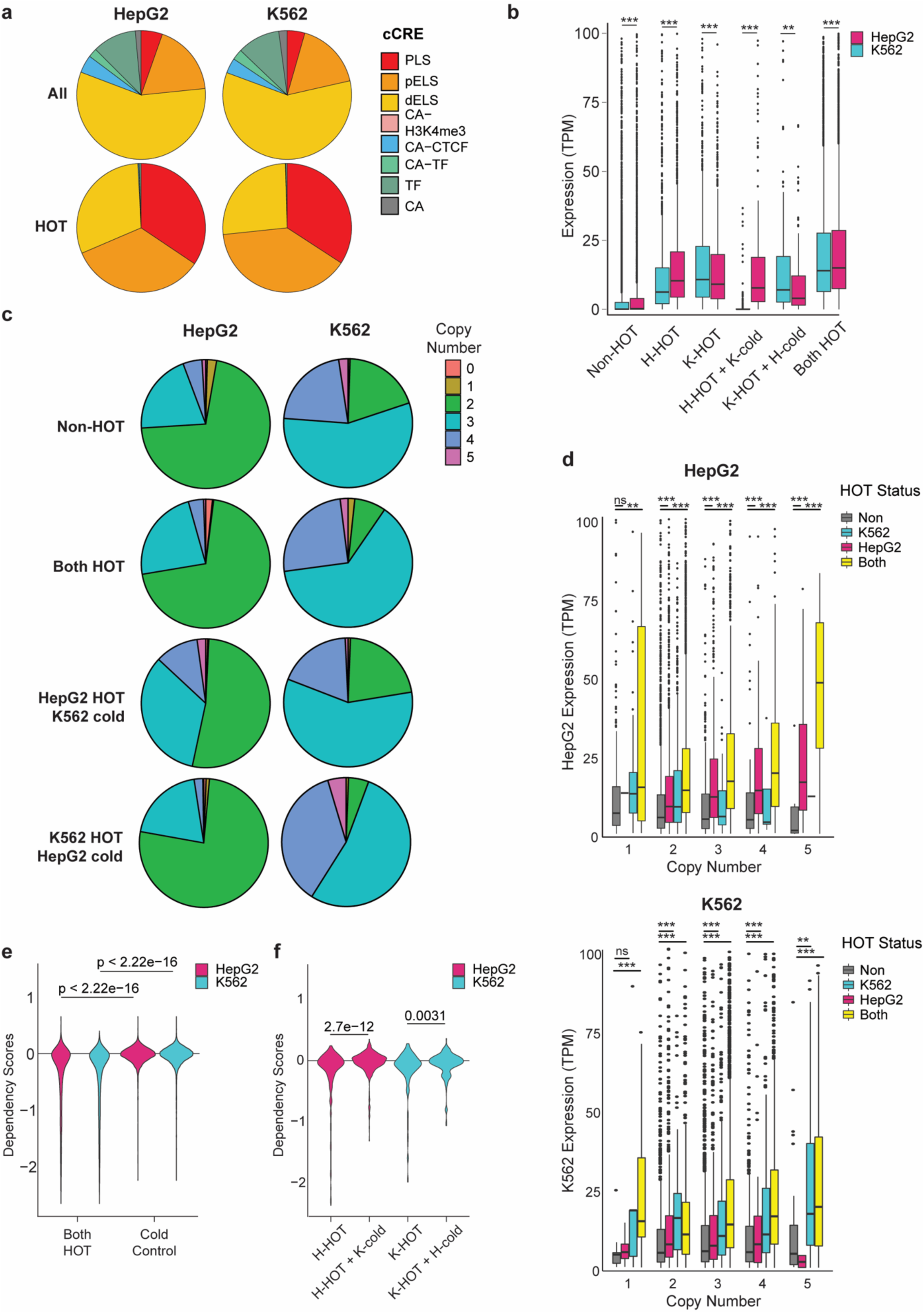
HOT site Characteristics across cell lines. a) Pie charts displaying the cCRE segmentation of all co-bound (top) or HOT (bottom) sites in each cell line. All co-bound regions and HOT sites were derived using common TF experiments. b) Boxplot showing gene expression levels in HepG2 and K562 for promoters regulated by the indicated HOT site classifications. ** = p < 0.005. *** = p < 0.0005. All p-values were determined using the paired Kruskal-Wallis test. c) Pie charts displaying the HepG2 and K562 copy number distribution for the indicated HOT site categories. d) RNA-seq expression levels for HepG2 (top) and K562 (bottom) genes whose promoters are HOT in either cell type, both, or neither, binned by copy number. ns = not significant; ** = p < 0.005; *** = p < 0.0005. All p-values were determined using the Wilcoxon rank sum test. e) HepG2 and K562 dependency scores for promoters which are HOT in both cell types compared to expression-matched cold genes. f) HepG2 and K562 dependency scores for genes which are HOT in the cell type of interest and either warm or cold in the alternate cell type. H-HOT: HepG2 HOT; K-HOT: K562 HOT; H-cold: HepG2 cold; K-cold: K562 cold

We also noted that approximately half of the HOT sites classified as cell-type specific were also highly bound in the alternate cell type, but did not pass the HOT cutoff and therefore were not considered a HOT region **(Extended Data Fig. 15d)**. We then calculated CAP enrichment in the HOT sites of one cell type versus the other by comparing HOT versus “cold” sites (bound by less than 10 CAPs) in each cell type (**Extended Data Fig. 16a)**. We found that most CAPs were significantly enriched for the HOT sites of one cell type. Moreover, this enrichment was consistent across PLS, pELS, and dELS HOT subtypes and was not associated with CAP expression differences between cell types (**Extended Data Fig. 16b and Extended Data Tables 11-14**). This suggests that TFs display cell-type-specific affinity for HOT site binding and regulation, rather than universally broad binding of highly open chromatin regions.

Because HOT sites are largely concentrated around PLS and pELS regions, we mapped all promoter-overlapping HOT sites to genes and explored expression in each cell type. We found genes regulated by HOT sites in either cell line to display higher expression levels compared to non-HOT genes **(Fig. 7b).** When we compared genes regulated by HOT sites in one cell line which are “cold” in the other cell line, we found they were significantly higher expressed in the HOT cell line compared to the “cold” cell line, suggesting this class of HOT sites are enriched for cell type-specific genes **(Fig. 7b)**. GO analysis revealed K562 HOT HepG2 “cold” genes were enriched for cytokine production while HepG2 HOT K562 “cold” genes showed enrichment for several liver-related functions, including Cholesterol Transport and HDL particle remodeling **(Extended Data Tables 15-16**), as might be expected. Together, these results demonstrate that cell type-specific HOT sites display distinct CAP co-occupancy which regulate highly expressed genes governing cell type-specific functions.

### HOT site ploidy and functional importance across cell lines

Since HOT sites are determined by a large number of peaks in the same region, which is a function of reads mapped to a location in a given ChIP-seq experiment, it is possible loci of higher copy number are more likely to produce HOT sites than loci of lower copy number. To address this potential bias, we investigated the relationship between chromosomal ploidy and HOT site occurrence. We found the copy number distribution at HOT sites which are common among cell types was similar to genomic baseline **(Fig. 7c)**. In contrast, cell-type-specific HOT sites were located at higher copy number loci compared to genome baseline, regardless of CAP occupancy level in the other cell type **(Fig. 7c and Extended Data Fig. 17a)**. Additionally, for cell type-specific HOT loci in the alternate, non-HOT cell type, the same genomic regions are biased towards having a lower copy number than the genomic baseline **(Fig. 7c)**. To confirm certain HOT site enrichment at high copy number loci was not simply an artefact in ChIP-seq peak calling, we compared the relative number of reads mapped across ploidy bins for ChIP-seq input datasets and CAP experimental datasets. Importantly, input control and experimental read numbers were comparable across all ploidy bins **(Extended Data Fig. 17b)**, confirming there is no enrichment in experimental reads compared to input control at higher copy number loci. Despite the differences observed in copy number distribution across HOT site categories, we found that genes regulated by HOT sites across all categories exhibited increased expression compared to genes lacking HOT regions (“non-HOT regions”), independent of ploidy **(Fig. 7d)**. In agreement, HOT sites exhibited significantly higher STARR-seq activity compared to non-HOT regions (**Extended Data Fig. 17c**), suggesting that HOT-site activity is a function of CAP co-occupancy rather than open chromatin status.

Given the high expression level amongst HOT-regulated genes and the increase in copy number associated with cell type-specific HOT loci, we investigated the dependency of HOT-regulated genes on cell growth using the Cancer Dependency Map^53^. We found genes regulated by HOT sites in both cell types are consistently more important for cell survival in both cell types compared to “cold”, expression-matched genes in the same cell type (see Methods) (**Fig. 7e-f**). Meanwhile, genes regulated by HOT sites that are HOT in one cell type but “cold” in the other were not as critical for cell survival (**Fig. 7e-f**). Taken together, these findings suggest that genes with HOT promoters across cell types are not located at higher copy number regions but do regulate housekeeping genes critical for survival, whereas cell type-specific HOT sites are selectively enriched at high copy number regions and instead regulate lineage-specific genes.

## DISCUSSION

CAP co-occupancy is known to play a critical role in chromatin dynamics and transcription regulation, as several TFs bind DNA as dimers and many CAPs function as members of a larger protein complex. While many studies have explored the functions associated with small-scale CAP co-occupancies, such as TF and TF co-activators, few studies have examined large-scale co-occupancies genome-wide and most have relied on computational approaches due to a lack of direct binding data in multiple cell types. Due to the high number of combinatorial possibilities when investigating a large number of CAPs, studies have implemented computational tools to parse through the complex data including a “one vs. all” approach where the co-binding amongst one CAP is assessed across all other CAPs^15^, or an “all vs. all” approach such as SOMs^14^ or other integrative analyses^54^. However, these approaches, despite utilizing dimensionality reduction techniques, arguably do not result in reasonably interpretable findings, with the resulting groupings remaining highly complex. In this paper, we have generated a large set of binding data across multiple cell types and implemented topic modeling, which allows for flexible grouping and results in dimensionality reduction of about 30 orders of magnitude, going from ∼150,000 co-bound regions to ∼50 regulatory modules, compared to about 4 orders of magnitudes reduced by SOM in previous work^14^. We implemented topic modeling for over 150 additional CAP ChIP-seq datasets and provided substantial examples of the types of results that can be generated from this analysis. In effort to compare results, CAP z-scores representing the relative abundance across modules was used to identify CAP module enrichment and compare enrichment across cell line modules. It is worth noting that this is a binary approach (bound versus not bound) and does not incorporate CAP signal at each loci. In many instances we investigated CAP signal for specific module cases after module generation (**Figs. 3, 4, and 6, and Extended Data Figs. 2, 6, and 8)** in lieu of incorporating signal into the module generation, as it is currently unclear how to do so given the magnitude of this study. In future studies investigating a smaller number of CAPs, it would be interesting to test approaches incorporating CAP signal such as binning CAP peaks by signal values and reclassifying those peaks as an independent variables to input into RMD or a similar algorithm.

Defining the chromatin state of specific loci has evolved over time, with the most recent annotations being cCREs developed by ENCODE and SCREEN^12^. Recently, a handful of papers have noted the correlation between CAP co-occupancy and chromatin state^9,16^. In our analysis, we found unsupervised hierarchical clustering of CAP z-score values across modules identified module clusters that corresponded to specific chromatin states. Interestingly, this was even true for enhancers that are known to exhibit cell type-specificity. However, within state-defined clusters, modules of each cell line tended to cluster with each other, suggesting that while there is conserved CAP co-occupancy, there is still cell type-specific variability amongst CAP co-occupancy at chromatin states. As the majority of the CAPs analyzed here have similar expression levels in both cell lines, it would be worthwhile to explore additional mechanisms that explain the cell type-specific co-occupancies.

By comparing regulatory modules derived across K562 and HepG2 cell lines, we discovered 9 modules that were comprised of highly similar constituents, termed SRMs. While many of the SRM constituents have known PPIs, the extent to which these co-associations spanned cell lines was not known, apart from CTCF/Cohesin. CAPs which bind heterochromatin, specifically H3K27me3-high facultative chromatin, were identified among the SRMs (**Fig. 1c, Extended Data Fig. 2b and 6b**). Interestingly when further comparing constitutive heterochromatin co-binding between cell lines in **Extended Data Fig. 6d**, we found HepG2 exhibited a different combination of CAPs present at putative H3K9me3-high modules compared to K562 modules, involving CAPs associated with RNA binding. While consistent with existing reports in yeast that RNA machinery can establish heterochromatin sites^55–57^, this finding warrants further investigation using techniques that enrich for heterochromatin as opposed to euchromatin-enriched ChIP-seq^58^. By investigating the regulatory activity and motif enrichment of all regions and SRM CAPs, respectively, we demonstrated that most regions bound by SRMs are cell type-specific. We provided examples for when this could be due to aCAPs or discordant motif recognition. Chromatin modifying complexes, such as the Polycomb repressor complex, are recruited to DNA via TF-independent mechanisms^59^, and therefore other mechanisms beyond TF cooperation and genomic accessibility may explain the differential binding localizations of SRM complexes replete of TFs such as the HetChrom SRM. Further studies exploring the hierarchies of SRM recruitment, transcriptional targets, and co-occupancies in additional cell lines would be extremely useful in further defining the conserved regulatory functions of SRMs. Additionally, as the cell lines explored in this analysis are both cancer cell lines, it would be worthwhile to expand future analyses to non-cancer contexts, as SRMs could represent cancer-specific epigenetic modalities.

The identification of CAP co-occupancy could be the result one of four conditions: 1) co-occupied CAPs cooperate with one another for regulatory functions, 2) co-occupied CAPs bind the same genomic window but do not cooperate for regulatory functions, 3) co-occupied CAPs bind to the same genomic window, but not simultaneously, and as such results are an artifact of piling up ChIP-seq datasets from a cell population, or 4) co-occupied CAPs within a module display a mixture of conditions 1-3. Although motif analysis, previous literature, and PPIs can be assessed to support CAP co-occupancy fitting one of the four proposed conditions, we chose to conduct *in silico* cooperation experiments using base-resolution neural network models of TF activity. As expanding this analysis beyond cooperation between two TFs would be exponentially more challenging, we chose to conduct motif scrambling experiments on 4,878 DNA-binding TF pairs present in the same regulatory module with passing quality prediction models. From this analysis we identified 592 TF pairs of different DBDs predicted to exhibit DNA recruitment cooperation, 91% of them having no known PPIs. Through this work we hope to provide examples of candidate TF pairs that require additional investigating in vitro as well as a new pipeline for investigating TF pairs genome-wide.

While the generation of regulatory modules provided meaningful information regarding CAP co-occupancy across cell lines, it is important to acknowledge the “greedy” nature of topic modeling. As a result of this method, co-bound regions do not need to be bound by every module CAP to be included in said module, explaining how several “promoter-” and “enhancer-” associated modules can be comprised of over 100 CAPs, but not be significantly enriched for HOT regions. Due to this feature, we conducted HOT site comparisons as a stand-alone analysis. From this we were able to address well-known doubts surrounding HOT site validity. For example, we provided substantial evidence suggesting HOT loci are not a result of highly accessible chromatin or high chromosomal copy numbers. We demonstrated that cell type-specific HOT sites are more likely to be in higher copy number regions, whereas shared HOT sites show no enrichment with higher copy number loci. Additionally, by comparing RNA-seq and STARR-seq data, we showed that both copy number and HOT site status contribute to gene expression levels. Finally, we determined that CAP enrichment at HOT site subtypes was cell type-specific and not dependent on differential CAP expression levels, demonstrating that HOT sites exhibit CAP specificity as opposed to being non-specific highly accessible loci.

This work aimed to 1) provide both a catalog of CAP co-occupancy patterns across two cell lines incorporating the largest number of ENCODE ChIP-seq datasets to date as well as 2) demonstrate examples of how this data can be used to generate new hypotheses surrounding CAP co-occupancy. Through analyzing CAP co-occupancy with chromatin state, cell agnostic CAP co-occupancy patterns, regulatory module interactions, candidate TF cooperators within regulatory modules, and HOT site subtypes, we provided a co-occupancy roadmap that can be used to generate new hypotheses for future work. Our study provides a valuable resource that can be referenced to and built upon.

## EXPERIMENTAL PROCEDURES

### ENCODE ChIP–seq

The protocols for ChIP–seq have been established and previously published on the ENCODE web portal (https://www.encodeproject.org/documents/). In brief, HepG2 and K562 cells were obtained from ATCC (HB-8065 and CCL-243, respectively), confirmed by morphological observation, and tested for negative mycoplasma status. Following cell crosslinking and chromatin shearing, samples were incubated with their respective antibodies and further processed to obtain the final DNA libraries. ENCODE has developed a set of antibody characterization standards which are published on the ENCODE web portal and consist of two validation steps. Raw fastq data was filtered for quality and PCR duplicates prior to alignment to the GRCh38 genome. For each CAP, peak calling was carried out using the SPP algorithm, with replicate consistency and peak ranking determined by irreproducible discovery rate (IDR) package^60,61^.

### ENCODE Experiment Selection

All available CAP-ChIP-seq datasets aligned to the human reference genome (GRCh38) were downloaded from the ENCODE portal on October 27th, 2022. CAP-ChIP-seq experiments were filtered for target redundancy, with experiments containing optimal IDR bed files, untagged cell lines, and recent experiments prioritized, resulting in 664 and 503 unique HepG2 and K562 TF ChIP-seq datasets, respectively. The CAP ChIP-seq final experiment list is available in **Extended Data Table 1**. Of note, at the time of publication K562 ENCSR000EGP had been revoked on ENCODE’s portal. We therefore removed ENCSR000EGP and the corresponding HepG2 ENCSR080UEM from all results, although these experiments were used (along with 540 other CAPs) to generate the regulatory modules. Related sequencing files including RNA-seq, STARR-seq, Hi-C, ATAC-seq, and histone-ChIP-seq are listed in **Extended Data Table 2**. To assess peak replicability across experiments, we used bedtools intersect (default parameters) and overlapped all peaks from HepG2 and K562 experiments listed in **Extended Data Table 1** with ChIP-seq experiments of the same target CAP in other ENCODE cell lines, when available (for HepG2 this was 391/664 experiments and for K562 this was 390/502 experiments).

### ATAC-seq coverage on subsampling of experiments

Experiments for shared CAPs of sufficient quality were repeatedly subsampled for 5, 10, and up to 180 (in increments of 5) experiments 50 times each. Sampled experiment peaks were merged, and the fraction of open base pairs based on ATAC-seq peaks was determined for each subsampling. Significant differences of coverage of open regions between the two cells was determined with the kruskal.test function in R.

### Regulatory Module Discovery and Module Pre-processing

Peak bed files were downloaded for each CAP ChIP-seq experiment and filtered to include peaks with a q-value of 0.05 or lower. For each cell line, a region-CAP matrix describing the distribution of CAP binding across co-bound regions was compiled from the 271 common CAP ChIP-seq datasets available for both HepG2 and K562 cells using the method described by Guo and Gifford^16^ and provided at http://groups.csail.mit.edu/cgs/gem/rmd/. Briefly, CAP ChIP-seq peak summit positions were expanded-/+ 150bp and overlapping peaks regions were merged until a final list of non-overlapping merged peaks was achieved. Only regions with at least 3 unique factors bound were kept for the Regulatory Module Discovery. To implement the Regulatory Module Discovery, the cell line matrix and filtered bed files were provided as input to the hierarchical Dirichlet processes (HDP) topic model using the parameters eta=0.1, init_topics=60, and max_iter=1000. Modules with fewer than 500 co-bound regions assigned were omitted from downstream analyses. CAPs were considered a member of a regulatory module if they passed the following thresholds. First, for each CAP, z-scores representing the number of regions bound per module were calculated. Second, for each module, z-scores representing the relative number of regions bound per CAP were calculated. For each module-CAP pair, a CAP was considered a member of the regulatory module if the calculated z-scores were greater than 0, the mean, for both conditions. If the module-CAP pair did not pass these thresholds, the module-CAP pair was assigned z-score values of 0 and was not included as a module constituent, thus eliminating lowly bound CAPs from modules. A Table summarizing the final module constituents identified in both cell lines is available in **Extended Data Table 3**. The full list of co-bound regions with their corresponding module assignment(s) are available in **Extended Data Tables 17 and 18**. Signal z-scores across module regions were calculated using the average signal value across the CAP peak present in that module region and adjusting for the mean and standard deviation of the averaged signal values across all module regions. TF family annotation was performed according to Lambert, SA et al.^62^ A549 ChIP-seq experiments used are listed in **Extended Data Table 1** and were processed as described above. TF ChIP-seq of untreated and treated samples were inputted as separate datasets for RMD. Principal component analysis was conducted using the stats R package.

### Module clustering and heatmap generation

To identify patterns amongst modules derived from K562 and HepG2 cells, we performed unsupervised, hierarchical clustering using CAP z-scores calculated independently of each cell line. CAPs were clustered according to Euclidean distance while modules were clustered according to Pearson correlation distance, as recommended by the RMD pipeline^16^. All heatmaps were generated using the ComplexHeatmap R package^63^.

### Chromatin state annotation

HepG2 and K562 combined genome segmentation files for GRCh38 were downloaded from the UCSC genome browser (http://genome.ucsc.edu/index.html). The V4 cCRE human dataset was provided in advance by the ENCODE Data Analysis Center and are available in ENC4079. Module regions were assigned to the nearest segmentation or cCRE locus using the ChIPSeeker R package^64^.

#### CAP contribution to chromatin state-associated modules

Module clusters corresponding to a specific chromatin state were determined using hierarchical clustering displayed in **Fig. 1c**. Modules derived from the same cell line and determined to be in the same chromatin state cluster were combined, and the percent of total regions bound by each CAP was calculated.

### Shared regulatory module (SRM) analysis

#### Classification

All regulatory modules, defined by CAP z-score values, were clustered by Pearson distance as shown in **Fig. 1c**. Pearson distance is defined as 1-r, where r is the Pearson correlation coefficient. Therefore modules with Pearson distances closer to 0 are more similar compare to modules with larger Pearson distances. Clusters whose Pearson distance was less than 0.3 and which contained at least one module from each cell line were classified as SRMs.

#### Distance Measurements

For each SRM group, regions from the corresponding K562 and HepG2 module were extracted and for each region, the closest region in the other cell line was calculated using AnnotatePeaks from the ChIPSeeker R package^64^.

#### Histone and ATAC-seq z-scores and regulatory ratio calculations

For all co-bound regions (the input data for regulatory module discovery) and SRM regions, the average H3K27ac, H3K4me^3^, H3K4me^1^, H3K27me^3^ and ATAC-seq signal values were calculated using the following strategy. For each histone mark, the global mean and standard deviation was calculated from all co-bound regions. For each module, the average signal values across all regions were then converted to z-scores using the global average and standard deviation. For regulatory ratios the difference between H3K4me^3^ and H3K4me^1^ or H3K27ac and H3K27me^3^ average z-scores were calculated for each SRM.

#### De Novo Motif pipeline

Peak files from DNA-bound TFs common amongst SRMs were overlapped with cell type-specific SRM regions and de novo motif analysis was conducted using MEME-ChIP^65^ and options-meme-nmotifs 5 -ccut 150 -streme-nmotifs 5 -meme-p 8 -fimo-skip -spamo-skip -centrimo-local. Motifs identified using meme and streme were combined and compared to existing motif databased using Tomtom. Tomtom results were filtered for peak centrality using centrimo enrichment and filtered for E values less than 0.05. Filtered motifs were then combined according to Jaspar motif clusters to remove redundant motifs^28^.

#### Known motif analysis and motif distance

For each SRM, regions were divided into HepG2 only, K562 only, or shared. Genomic regions were expanded by 500bp upstream and downstream and MEME XSTREME motif discovery was performed on each SRM subgroup. For motif distance calculations, motif positions were downloaded from https://www.vierstra.org/resources/ ^45^.

#### CAP vs CAP peak distance measurements

To determine the binding distance between two CAPs, the distance between peak summits was calculated for all co-bound regions and the distribution was plotted.

### Differential Expression Analysis

Differential expression analysis was performed using RNA-seq datasets **(Extended Data Table 2)** and the deseq2 package^66^ in R. Analysis was performed on all genes.

### Pathway Analysis

#### Regulatory Module pipeline

For Hi-C data, loop anchors were assigned to all genes within their genomic range. Gene sets were then calculated for GO pathway enrichment using the msigdbr R package^67^. For other regulatory module pathway enrichment, regions were mapped to genes using the following method. If module region was within 1.5 kb of a gene, the region was assigned to that gene. If the module region was not within 1.5 kb of any gene, the region was overlapped with PCHi-C “other end” peak list and then mapped to the corresponding “bait” gene.

#### SRM permutation analysis

For each SRM module-module pair, 10,000 permutation tests were conducted to determine the significance of SRM pairs distally regulating the same genes. We used the resampleRegions from the regioneR package^68^ to resample SRM module regions by chromosome using the universe of all regulatory module discovery co-bound regions for each cell line. For each permutation, resampled regions were assessed for their overlap of distally regulated genes, using cell line-specific PCHi-C defined interactions. All regions were assigned to PCHi-C “bait” genes if the regions overlapped PCHi-C “other ends” within 1.5 kbp. The empirical p-value was calculated as follows:

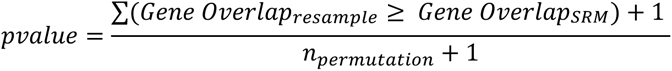

#### HOT site pipeline

HOT and cold sites were mapped to gene transcription start sites within 1kb. GO enrichment analysis was performed by providing gene lists for either HepG2-HOT, K562-Cold genes, or K562-HOT, HepG2-cold genes, as appropriate for the analysis, to the Gene Ontology web portal (http://geneontology.org/). A control list of genes which had HOT promoters in both cell types was provided.

### Hi-C loop and TAD analysis

#### Identification of Hi-C loop anchors, intra-loop span and TAD boundaries

HepG2 and K562 intact Hi-C experiments used for this analysis are listed in **Extended Data Table 2**. Bedpe loop files downloaded from ENCODE were used to identify chromatin loops. To identify module regions or CAP peaks that overlapped loop anchors, we used bedtools pairToBed under default parameters. For assessment of regions/peaks that overlap intra-loop span, we used bedtools pairToBed -type ispan, which subsets loops where the inner region overlaps the input bed file. For overlap analysis, any region/peak that overlapped the loop anchor and intra-loop span was classified as a loop anchor overlap and removed from the intra-loop span analysis. Contact domain bedpe files downloaded from ENCODE were used to denote TAD regions. The start of the most upstream “start” genomic position represented the first TAD boundary and the most downstream “end” genomic region represented the second TAD boundary for each TAD region. We then expanded those loci 5kb in each direction and classified that window as the TAD boundary (width = 10kb). The intra-TAD span was all loci within those expanded boundaries. Overlap with TAD boundaries and TAD intra-span regions was conducted using bedtools intersect. CAP signal across all regions of interest was calculated using deepTools multiBigwigSummary.

#### Classification of cell type-specific, shared loops, and active Hi-C loops

Hi-C loops that overlapped at both loop ends were classified as shared loops and identified using bedtools pairToPair -type both. Loops that did not overlap were considered cell type-specific. Hi-C loops where either loop end overlapped with H3K27ac peaks were classified as “active” loops. CAP enrichment at active loops was determined using the following equation:

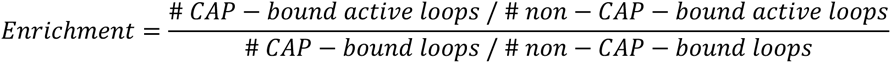

#### Aggregate peak analysis

Aggregate peak analysis was conducted using Juicer the APA function with normalized Hi-C data under default options^69^.

### Differentially Bound Modules

Differentially bound modules (DBMs) were defined as module pairs with a minimum Pearson distance value of 0.4 and at least 30% of overlapping regions between the two modules. For all HepG2-K562 module pairs we calculated the Pearson distance based on the CAP z-scores as described above. We then also calculated the ratio of overlapped regions, from both the K562 and HepG2 module perspective, and kept the maximum ratio between the two. Then all pairs that passed the cutoff values were classified as DBMs.

### Protein-protein Interactions

Protein-protein interactions were calculated using the StringDb homo sapiens database of physical links^70^. Any protein-protein interaction with a “combined score” greater than 0 was considered to have a known interaction.

### Promoter Capture Hi-C (PCHi-C)

We performed PCHi-C in-situ using the Arima PCHi-C Kit (catalog no. A301010, A302010, A303010), Arima Genomics Inc.). Briefly , 10^6^ cells per sample were cross-linked in PBS for 10 minutes at a final formaldehyde concentration of 2%. Subsequently, the crosslinked chromatin was digested with a restriction enzyme cocktail, and the digested ends were labeled with a biotinylated nucleotide by filling in the 5’-overhangs. The next steps were DNA proximity ligation, reverse crosslinking and DNA purification. The DNA was then fragmented to an average fragment size of 400bp using a Covaris S2 sonicator, followed by double sided bead-based size selection to 200-600bp (SPRI beads (catalog no. B23318, Beckman Coulter Inc.). The biotinylated fragments were then enriched using streptavidin Beads, and processed into Illumina-compatible sequencing libraries using the Arima library prep module.

To enrich for promotor regions the Hi-C libraries were hybridized with a set of custom RNA-probes that target the promoters of 23,711 genes. The final libraries were sequenced using an Illumina NextSeq sequencer to generate paired-end 150-bp reads.

#### Data preprocessing

Sequencing files were processed using the Arima Capture Hi-C pipeline (https://github.com/ArimaGenomics/CHiC) using a 5 kb resolution and according to the recommended guidelines. Briefly, for each replicate, raw reads were mapped and filtered to GRCh38 genome using HiCUP^13^ and significant loop interactions were called using CHiCAGO^14^. Loops identified in both replicates were used for all subsequent analyses.

### Module-module interaction analysis

#### Permutation Tests for loop end interactions

For each module-module pair (A,B), 100 permutation tests were conducted to determine the significance of module-module interactions at loop ends. We used the permTest function from the regioneR package^68^ and our random sampling universe were all of the regulatory module discovery co-bound regions for each cell line. We designed a custom evaluation function to determine the number of module-module interactions via PCHi-C loops. Module regions were overlapped with both PCHi-C loop ends (end1, end2). If Module A overlapped end1 and Module B overlapped end 2 within the same loop, and vice versa, then the loop was considered to be an overlap. If the same co-bound region was assigned to both modules, it was removed from Module B regions to avoid a high rate of false positive loop overlaps. For the resulting data, modules were considered to overlap if the p-value was <0.01 and the predicted z-score was greater than 0.

#### H3K27ac signal at PCHi-C loop anchors

For each module pair, module-module interactions via PCHi-C loops were determined as described above. Using multiBigwigSummary from deepTools, the H3K27ac signal at overlapping loop ends were calculated.

### TF combinatorial binding detection

We first created a matrix of TF pairs that are present in the same module for either K562 or HepG2. A CAP was determined to be a TF if it had a known DNA-binding motif from the Viestra database^45^. The Pearson correlation between ChIP-seq signals at co-bound sites was calculated for each TF pair. To limit the computational load for predictive analysis, only TFs whose signal correlation was greater than 0 were kept for further analysis. TF pairs that passed this threshold were then processed using our pipeline. First for a pair of TFs, TF_A_ and TF_B_, we determined the TF_A_ peaks that overlap TF_B_ using BEDTools^71^ intersect. These co-bound genomic regions were then scanned for the TF_B_ motif, using the coordinates available from the Viestra motif database^45^. The identified motifs were then scrambled, keeping the same nucleotide composition, to produce 5 uniquely perturbed motif sequences. These 5 perturbed sequences and the unmodified control sequence were then used for TF_A_ ChIP-seq signal predictions using the base-resolution models provided by the Kundaje lab and available on ENCODE (ENCIDs are listed in **Extended Data Table 9**). The base-resolution models were quality checked using two metrics: 1) primary motif enrichment based on existing literature and 2) a counts Pearson correlation coefficient of 0.5 or greater. For each TF, five prediction models were developed. Therefore five predictions were computed for each sequence, resulting in 25 total motif-scrambled signal predictions and 5 total control signal predictions for each initial sequence. For each TF_A_, we performed a paired t-test between the read counts of each control and motif-scrambled predictions and then calculated the effect size as the fold change of the median read count of motif-scrambled sequences over the median read count of control sequences. TF pairs were considered significant if the read count fold change >= 1.2 or <= 0.83 and the p-value <= 0.001. As expected, TF pairs showed dependency on each other’s motifs as the majority of fold changes were less than 1 (log2 fold change < 0), suggesting a reduction in predicted read counts at motif-deleted regions. Code for this analysis can be found at https://github.com/theBigTao/HepG2-K562-Toolkits. Note HepG2 ZNF143 experiment was included in this analysis when applicable.

### HOT site analysis

#### HOT site definition

Because HOT sites are defined by a certain percent of CAPs bound among a set, we first removed experiments with few peaks or low quality peaks. For every experiment, we restricted to peaks with a q-value of 0.01 or lower. Experiments with fewer than 500 peaks remaining were removed. We identified CAPs with an experiment of sufficient quality in both HepG2 and K562 which produced 190 CAPs from which to identify HOT sites. We then restricted peaks to +/-50 base pairs surrounding the peak center, and merged peaks across experiments into regions. Any region which was made up of peaks from at least 1/3 of the experiments (rounded down) in this set was defined as a HOT site. Regions that were not HOT sites in either cell line were categorized as “non-HOT”.

#### Cold sites Definition

After identifying HOT sites which appeared to be unique in a cell type, we defined cold sites in the alternate cell type as those regions which had 10 or fewer CAPs bound in a region which was HOT in a given cell type.

#### CAP Enrichment in HOT sites

For each CAP, we created a table whose first column was the number of HOT sites lacking the CAP of interest and the number of HOT sites containing the CAP of interest in HepG2. The second column was the same information, but in K562. We performed a chi-squared analysis on this table using the chisq function in R version 3.6.1. The enrichment value displayed in graphs is:

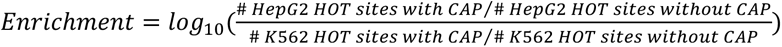

#### Dependency Analysis

Dependency data was download from DepMap https://depmap.org/portal/^53^. For dependency comparisons, we broke genes into a focal group and a comparison group. To control for differences in expression, we iteratively went through our focal group and identified which gene in the comparison group had the closest expression level. Once identified, we removed that control gene from the comparison pool. We continued this process until the comparison pool was depleted or we had a paired comparison for every focal gene. Unpaired genes were discarded.

#### Karyotype data

Information on copy number of different chromosomal regions was identified for HepG2^72^ and for K562^73^. To confirm that enrichment of HOT sites at higher chromosomal copy number regions was not an artefact, we compared the reads mapping to genomic regions in a given ChIP-seq dataset compared to its control. We broke the genome into 2000 bp bins and determined the number of reads mapping to each bin and its ploidy. We then determined the total number of reads mapping to each region in both the dataset and the control bams. Bins where both the dataset and the control had 0 normalized reads were removed. We then normalized these numbers by the following formula:

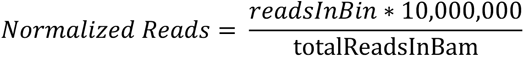

#### STARR-seq analysis

K562 STARR-seq data was acquired from the ENCODE portal under accessions ENCFF338YDG, ENCFF817HOX, and ENCFF906QND. A custom script was used to determine the average STARR-seq activity over the promoter region for each replicate. We then took the average of the three reps.

### Visualization and statistical analysis

All heatmaps were generated using the Complex Heatmap R package. Upset plots were generated using the Upset R package^74^. TF ChIP-seq signal plots were generated using computeMatrix and plotProfile commands from the deepTools suite^75^. Hi-C contact matrices were generated using Juicebox available at https://aidenlab.org/juicebox/ and IGV^76^. Unless otherwise stated, the remaining plots were generated using the ggplot2 R package^77^. All operations were performed under R version 3.6.

## REFERENCES

1. Heinz, S. et al. Simple Combinations of Lineage-Determining Transcription Factors Prime cis-Regulatory Elements Required for Macrophage and B Cell Identities. Molecular Cell 38, 576–589 (2010).

2. Yosef, N. et al. Dynamic regulatory network controlling TH17 cell differentiation. Nature 496, 461–468 (2013).

3. Corces, M. R. et al. Lineage-specific and single-cell chromatin accessibility charts human hematopoiesis and leukemia evolution. Nat Genet 48, 1193–1203 (2016).

4. Visel, A. et al. ChIP-seq accurately predicts tissue-specific activity of enhancers. Nature 457, 854–858 (2009).

5. Chen, X. et al. Integration of External Signaling Pathways with the Core Transcriptional Network in Embryonic Stem Cells. Cell 133, 1106–1117 (2008).

6. Zanconato, F. et al. Genome-wide association between YAP/TAZ/TEAD and AP-1 at enhancers drives oncogenic growth. Nat Cell Biol 17, 1218–1227 (2015).

7. Robertson, G. et al. Genome-wide profiles of STAT1 DNA association using chromatin immunoprecipitation and massively parallel sequencing. Nat Methods 4, 651–657 (2007).

8. Johnson, D. S., Mortazavi, A., Myers, R. M. & Wold, B. Genome-Wide Mapping of in Vivo Protein-DNA Interactions. Science 316, 1497–1502 (2007).

9. Partridge, E. C. et al. Occupancy maps of 208 chromatin-associated proteins in one human cell type. Nature 583, 720–728 (2020).

10. Yip, K. Y. et al. Classification of human genomic regions based on experimentally determined binding sites of more than 100 transcription-related factors. Genome Biol 13, R48 (2012).

11. Ramaker, R. C. et al. Dissecting the regulatory activity and sequence content of loci with exceptional numbers of transcription factor associations. Genome Res. 30, 939–950 (2020).

12. 12. The ENCODE Project Consortium et al. Expanded encyclopaedias of DNA elements in the human and mouse genomes. Nature 583, 699–710 (2020).

13. Neph, S. et al. Circuitry and Dynamics of Human Transcription Factor Regulatory Networks. Cell 150, 1274– 1286 (2012).

14. Xie, D. et al. Dynamic trans-Acting Factor Colocalization in Human Cells. Cell 155, 713–724 (2013).

15. Gerstein, M. B. et al. Architecture of the human regulatory network derived from ENCODE data. Nature 489, 91–100 (2012).

16. Guo, Y. & Gifford, D. K. Modular combinatorial binding among human trans-acting factors reveals direct and indirect factor binding. BMC Genomics 18, 45 (2017).

17. Avsec, Ž., et al. Base-resolution models of transcription-factor binding reveal soft motif syntax. Nat Genet 53, 354–366 (2021).

18. Lozzio, C. & Lozzio, B. Human chronic myelogenous leukemia cell-line with positive Philadelphia chromosome. Blood 45, 321–334 (1975).

19. Aden, D. P., Fogel, A., Plotkin, S., Damjanov, I. & Knowles, B. B. Controlled synthesis of HBsAg in a differentiated human liver carcinoma-derived cell line. Nature 282, 615–616 (1979).

20. Teh, Y. W., Jordan, M. I., Beal, M. J. & work(s):, D. M. B. R. Hierarchical Dirichlet Processes. Journal of the American Statistical Association 101, 1566–1581 (2006).

21. Lou, S. et al. TopicNet: a framework for measuring transcriptional regulatory network change. Bioinformatics 36, i474–i481 (2020).

22. Wang, M. et al. Topic model-based mass spectrometric data analysis in cancer biomarker discovery studies. BMC Genomics 17, 545 (2016).

23. Vockley, C. M. et al. Direct GR Binding Sites Potentiate Clusters of TF Binding across the Human Genome. Cell 166, 1269–1281.e19 (2016).

24. Zhang, Y., An, L., Yue, F. & Hardison, R. C. Jointly characterizing epigenetic dynamics across multiple human cell types. Nucleic Acids Res 44, 6721–6731 (2016).

25. Heintzman, N. D. et al. Histone modifications at human enhancers reflect global cell-type-specific gene expression. Nature 459, 108–112 (2009).

26. Dixon, J. R. et al. Chromatin architecture reorganization during stem cell differentiation. Nature 518, 331– 336 (2015).

27. Dixon, J. R. et al. Topological domains in mammalian genomes identified by analysis of chromatin interactions. Nature 485, 376–380 (2012).

28. Castro-Mondragon, J. A. et al. JASPAR 2022: the 9th release of the open-access database of transcription factor binding profiles. Nucleic Acids Research 50, D165–D173 (2022).

29. Elagib, K. E. et al. RUNX1 and GATA-1 coexpression and cooperation in megakaryocytic differentiation. Blood 101, 4333–4341 (2003).

30. Reizel, Y. et al. Collapse of the hepatic gene regulatory network in the absence of FoxA factors. Genes Dev. 34, 1039–1050 (2020).

31. Lee, C. S., Friedman, J. R., Fulmer, J. T. & Kaestner, K. H. The initiation of liver development is dependent on Foxa transcription factors. Nature 435, 944–947 (2005).

32. Gruber, S., Haering, C. H. & Nasmyth, K. Chromosomal Cohesin Forms a Ring. Cell 112, 765–777 (2003).

33. Wendt, K. S. et al. Cohesin mediates transcriptional insulation by CCCTC-binding factor. Nature 451, 796– 801 (2008).

34. Grubert, F. et al. Landscape of cohesin-mediated chromatin loops in the human genome. Nature 583, 737– 743 (2020).

35. Vos, E. S. M. et al. Interplay between CTCF boundaries and a super enhancer controls cohesin extrusion trajectories and gene expression. Molecular Cell 81, 3082–3095.e6 (2021).

36. Parelho, V. et al. Cohesins Functionally Associate with CTCF on Mammalian Chromosome Arms. Cell 132, 422–433 (2008).

37. Kagey, M. H. et al. Mediator and cohesin connect gene expression and chromatin architecture. Nature 467, 430–435 (2010).

38. Shen, M. et al. Hepatic ARID3A facilitates liver cancer malignancy by cooperating with CEP131 to regulate an embryonic stem cell-like gene signature. Cell Death Dis 13, 732 (2022).

39. Liao, T.-T. et al. let-7 Modulates Chromatin Configuration and Target Gene Repression through Regulation of the ARID3B Complex. Cell Reports 14, 520–533 (2016).

40. Rhee, C. et al. Arid3a is essential to execution of the first cell fate decision via direct embryonic and extraembryonic transcriptional regulation. Genes Dev. 28, 2219–2232 (2014).

41. Qian, M. et al. Whole-transcriptome sequencing identifies a distinct subtype of acute lymphoblastic leukemia with predominant genomic abnormalities of *EP300* and *CREBBP*. Genome Res. 27, 185–195 (2017).

42. Ortabozkoyun, H. et al. CRISPR and biochemical screens identify MAZ as a cofactor in CTCF-mediated insulation at Hox clusters. Nat Genet 54, 202–212 (2022).

43. Xiao, T., Li, X. & Felsenfeld, G. The Myc-associated zinc finger protein (MAZ) works together with CTCF to control cohesin positioning and genome organization. Proc Natl Acad Sci USA 118, e2023127118 (2021).

44. Fulco, C. P. et al. Activity-by-contact model of enhancer–promoter regulation from thousands of CRISPR perturbations. Nat Genet 51, 1664–1669 (2019).

45. Vierstra, J. et al. Global reference mapping of human transcription factor footprints. Nature 583, 729–736 (2020).

46. Ramalingam, Vivekanandan, Kundaje, Anshul. MODISCO [Manuscript in preparation]. TBD.

47. The ENCODE Project Consortium. ENCODEv4 Flagship [Manuscript in preparation]. Nature.

48. Eferl, R. & Wagner, E. F. AP-1: a double-edged sword in tumorigenesis. Nat Rev Cancer 3, 859–868 (2003).

49. Alder, O. et al. Hippo Signaling Influences HNF4A and FOXA2 Enhancer Switching during Hepatocyte Differentiation. Cell Reports 9, 261–271 (2014).

50. Cebola, I. et al. TEAD and YAP regulate the enhancer network of human embryonic pancreatic progenitors. Nat Cell Biol 17, 615–626 (2015).

51. Stein, C. et al. YAP1 Exerts Its Transcriptional Control via TEAD-Mediated Activation of Enhancers. PLoS Genet 11, e1005465 (2015).

52. Li, H., Liu, F., Ren, C., Bo, X. & Shu, W. Genome-wide identification and characterisation of HOT regions in the human genome. BMC Genomics 17, 733 (2016).

53. Broad. DepMap. (2022).

54. Wang, R. et al. Hierarchical cooperation of transcription factors from integration analysis of DNA sequences, ChIP-Seq and ChIA-PET data. BMC Genomics 20, 296 (2019).

55. Volpe, T. A. et al. Regulation of Heterochromatic Silencing and Histone H3 Lysine-9 Methylation by RNAi.Science 297, 1833–1837 (2002).

56. Bayne, E. H. et al. Stc1: A Critical Link between RNAi and Chromatin Modification Required for Heterochromatin Integrity. Cell 140, 666–677 (2010).

57. Bühler, M., Verdel, A. & Moazed, D. Tethering RITS to a Nascent Transcript Initiates RNAi- and Heterochromatin-Dependent Gene Silencing. Cell 125, 873–886 (2006).

58. Becker, J. S. et al. Genomic and Proteomic Resolution of Heterochromatin and Its Restriction of Alternate Fate Genes. Molecular Cell 68, 1023–1037.e15 (2017).

59. Perino, M. et al. MTF2 recruits Polycomb Repressive Complex 2 by helical-shape-selective DNA binding. sNat Genet 50, 1002–1010 (2018).

60. Landt, S. G. et al. ChIP-seq guidelines and practices of the ENCODE and modENCODE consortia. Genome Research 22, 1813–1831 (2012).

61. Li, Q., Brown, J. B., Huang, H. & Bickel, P. J. Measuring reproducibility of high-throughput experiments. Ann. Appl. Stat. 5, (2011).

62. Lambert, S. A. et al. The Human Transcription Factors. Cell 172, 650–665 (2018).

63. Gu, Z., Eils, R. & Schlesner, M. Complex heatmaps reveal patterns and correlations in multidimensional genomic data. Bioinformatics 32, 2847–2849 (2016).

64. Yu, G., Wang, L.-G. & He, Q.-Y. ChIPseeker: an R/Bioconductor package for ChIP peak annotation, comparison and visualization. Bioinformatics 31, 2382–2383 (2015).

65. Bailey, T. L., Johnson, J., Grant, C. E. & Noble, W. S. The MEME Suite. Nucleic Acids Res 43, W39–W49 (2015).

66. Anders, S. & Huber, W. Differential expression of RNA-Seq data at the gene level – the DESeq package. 24.

67. Subramanian, A. et al. Gene set enrichment analysis: A knowledge-based approach for interpreting genome-wide expression profiles. Proc. Natl. Acad. Sci. U.S.A. 102, 15545–15550 (2005).

68. Gel, B. et al. regioneR: an R/Bioconductor package for the association analysis of genomic regions based on permutation tests. Bioinformatics 32, 289–291 (2016).

69. Durand, N. C. et al. Juicer Provides a One-Click System for Analyzing Loop-Resolution Hi-C Experiments. Cell Systems 3, 95–98 (2016).

70. Szklarczyk, D. et al. STRING v10: protein–protein interaction networks, integrated over the tree of life. Nucleic Acids Research 43, D447–D452 (2015).

71. Quinlan, A. R. & Hall, I. M. BEDTools: a flexible suite of utilities for comparing genomic features. Bioinformatics 26, 841–842 (2010).

72. Zhou, B. et al. Haplotype-resolved and integrated genome analysis of the cancer cell line HepG2. Nucleic Acids Research 47, 3846–3861 (2019).

73. Zhou, B. et al. Comprehensive, integrated, and phased whole-genome analysis of the primary ENCODE cell line K562. Genome Res. 29, 472–484 (2019).

74. Conway, J. R., Lex, A. & Gehlenborg, N. UpSetR: an R package for the visualization of intersecting sets and their properties. Bioinformatics 33, 2938–2940 (2017).

75. Ramírez, F. et al. deepTools2: a next generation web server for deep-sequencing data analysis. Nucleic Acids Res 44, W160–W165 (2016).

76. Thorvaldsdottir, H., Robinson, J. T. & Mesirov, J. P. Integrative Genomics Viewer (IGV): high-performance genomics data visualization and exploration. Briefings in Bioinformatics 14, 178–192 (2013).

77. Wickham, H. *Ggplot2: Elegant Graphics for Data Analysis*. (Springer New York, New York, NY, 2009). doi:10.1007/978-0-387-98141-3.

