## Supplementary Figures for "Comparative atlas of genome-wide chromatin-associated protein co-occupancy"

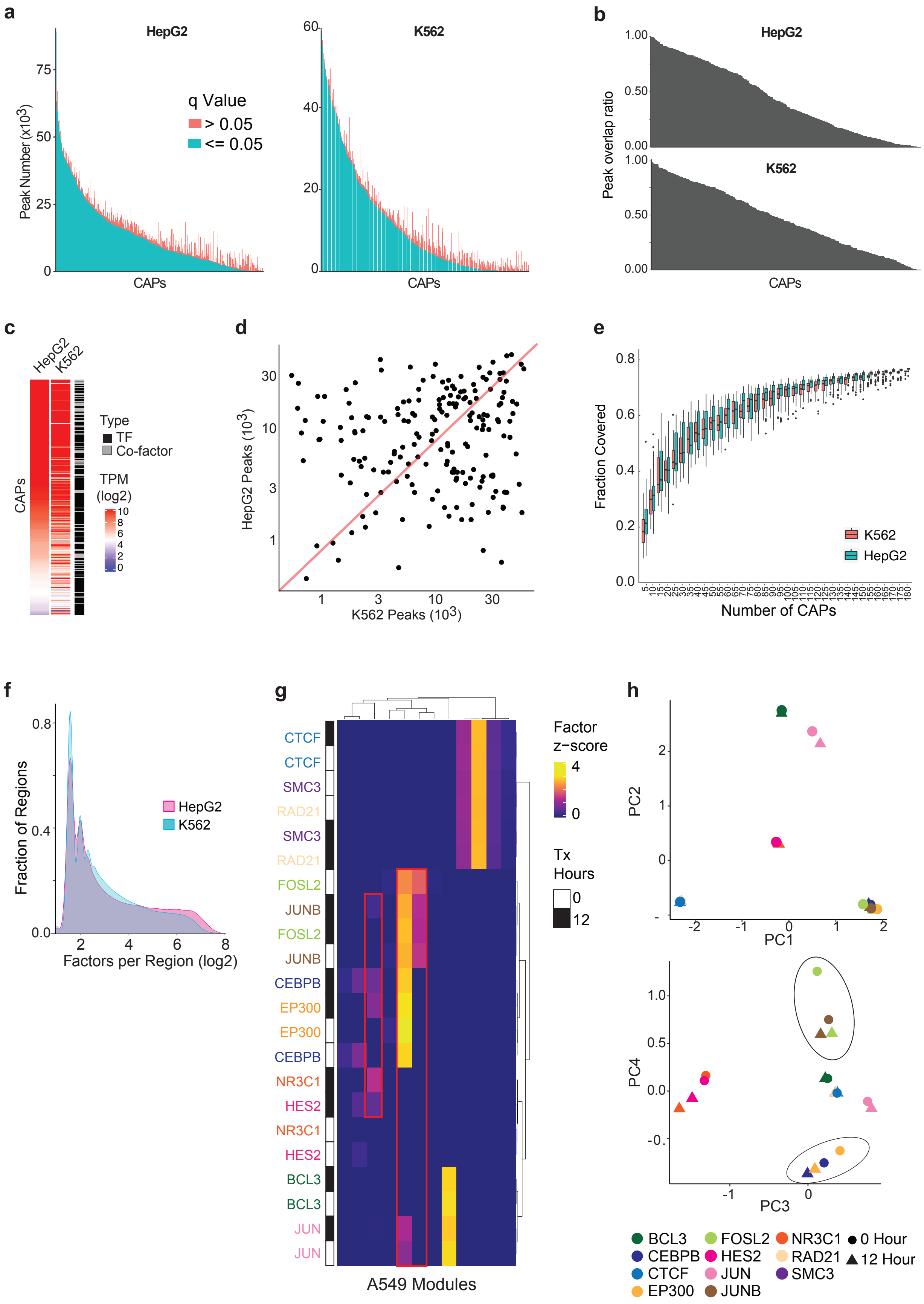
a
HepG2
K562
Peak Number (x10^3)
q Value
> 0.05
<= 0.05
CAPs
b
Peak overlap ratio
c
Type
TF
Co-factor
TPM
(log2)
d
HepG2 Peaks (10^3)
K562 Peaks (10^3)
e
Fraction Covered
Number of CAPs
f
Fraction of Regions
Factors per Region (log2)
g
Factor
z−score
Tx
Hours
A549 Modules
CTCF
SMC3
RAD21
FOSL2
JUNB
CEBPB
EP300
NR3C1
HES2
BCL3
JUN
h
PC1
PC2
PC3
PC4
0 Hour
12 Hour

**Extended Data Figure 1. ChIP-seq dataset and co-binding characteristics.**

a) Stacked bar plot depicting HepG2 (left) and K562 (right) ChIP-seq experiments used in this study. Each bar is colored according to the number of peaks that are above and below the indicated q-values.

b) The fraction of HepG2 (top) and K562 (bottom) ChIP-seq peaks replicated in other available ENCODE ChIP-seq experiments for the same TF. Note several ChIP-seq datasets used in this study are only replicated in other cell lines which are likely to have cell type-specific binding events.

c) Heatmap depicting the expression levels (TPM) in HepG2 and K562 across all 271 CAPs used in this analysis.

d) Peak number (q-value $< 0.01$) scatterplot for experiments of shared TFs in HepG2 and K562. Red line shows $x=y$. Peak numbers are loosely correlated between experiments (Spearman's Rho= 0.13, $p = 0.064$), and neither cell type consistently has more peaks than the other (k-s test $p= 0.51$).

e) ATAC-seq peak coverage in HepG2 and K562 as a function of experiment subsampling. HepG2 generally has higher coverage of open ATAC-seq regions than K562 (Kruskal-Wallis Rank-sum test $p = 0.000052$).

f) Distribution of the factors bound per co-bound region in HepG2 and K562 cell lines. This refers to the input region dataset for regulatory module discovery.

g) Heatmap depicting RMD modules identified for A549 dexamethasone or control treated ChIP-seq experiments. Modules with co-association differences following treatment are highlighted.

h) Scatterplots of principal components highlighting the differences observed in CAP co-association before and after dexamethasone treatment in A549 cells.

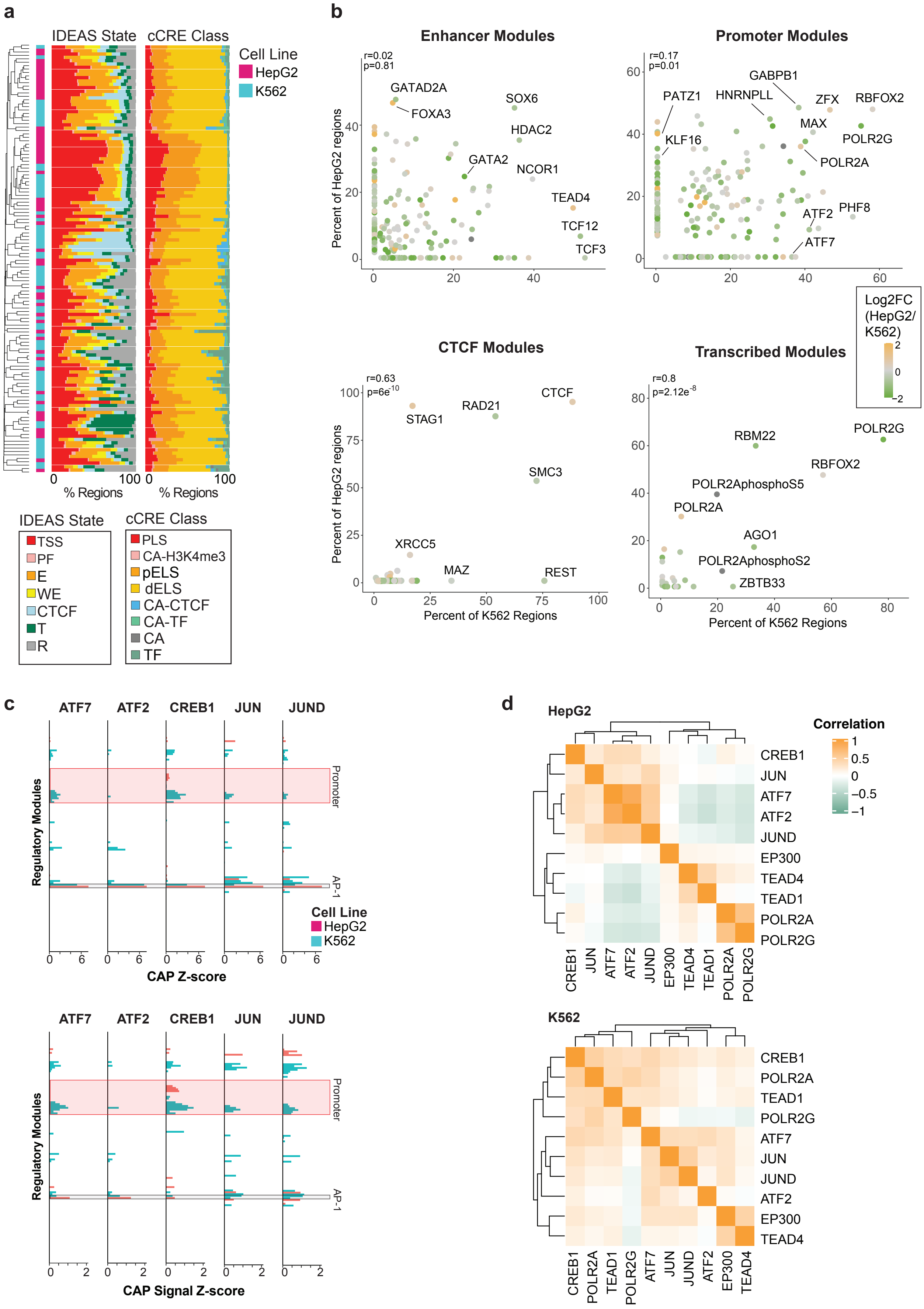
a
IDEAS State
cCRE Class
Cell Line
HepG2
K562
% Regions
% Regions
IDEAS State
TSS
PF
E
WE
CTCF
T
R
cCRE Class
PLS
CA-H3K4me3
pELS
dELS
CA-CTCF
CA-TF
CA
TF
b
Enhancer Modules
r=0.02
p=0.81
GATAD2A
FOXA3
SOX6
HDAC2
GATA2
NCOR1
TEAD4
TCF12
TCF3
Promoter Modules
r=0.17
p=0.01
GABPB1
PATZ1
HNRNPLL
ZFX
RBFOX2
KLF16
MAX
POLR2G
POLR2A
ATF2
PHF8
ATF7
Percent of HepG2 regions
Log2FC (HepG2/ K562)
CTCF Modules
r=0.63
p=6e-10
CTCF
RAD21
STAG1
SMC3
XRCC5
MAZ
REST
Transcribed Modules
r=0.8
p=2.12e-8
POLR2G
RBM22
RBFOX2
POLR2AphosphoS5
POLR2A
AGO1
POLR2AphosphoS2
ZBTB33
Percent of K562 Regions
Percent of K562 Regions
c
ATF7
ATF2
CREB1
JUN
JUND
Regulatory Modules
Promoter
AP-1
Cell Line
HepG2
K562
CAP Z-score
CAP Signal Z-score
d
HepG2
Correlation
CREB1
JUN
ATF7
ATF2
JUND
EP300
TEAD4
TEAD1
POLR2A
POLR2G
K562
CREB1
POLR2A
TEAD1
POLR2G
ATF7
JUN
JUND
ATF2
EP300
TEAD4

**Extended Data Figure 2. Regulatory state characteristics of co-binding modules.**

a) Distribution of IDEAS chromatin state (left) and cCRE annotation (right) across regulatory modules. Modules are ordered the same as in Fig. 1c.

b) Scatterplots showing the percent of regions bound in module clusters enriched for the indicated chromatin states. Dot colors correspond to the log2FC expression values comparing HepG2 and K562 normalized counts. Pearson correlation was used to calculate correlation and p-values.

c) Distribution of number of regions bound across modules for the indicated TFs. Either binding z-score (top) or ChIP-seq signal z-score (bottom) values are plotted.

d) Heatmaps displaying the binary (bound or not bound) co-binding Pearson correlations for the indicated CAP pairs across all co-bound regions provided as input to RMD.

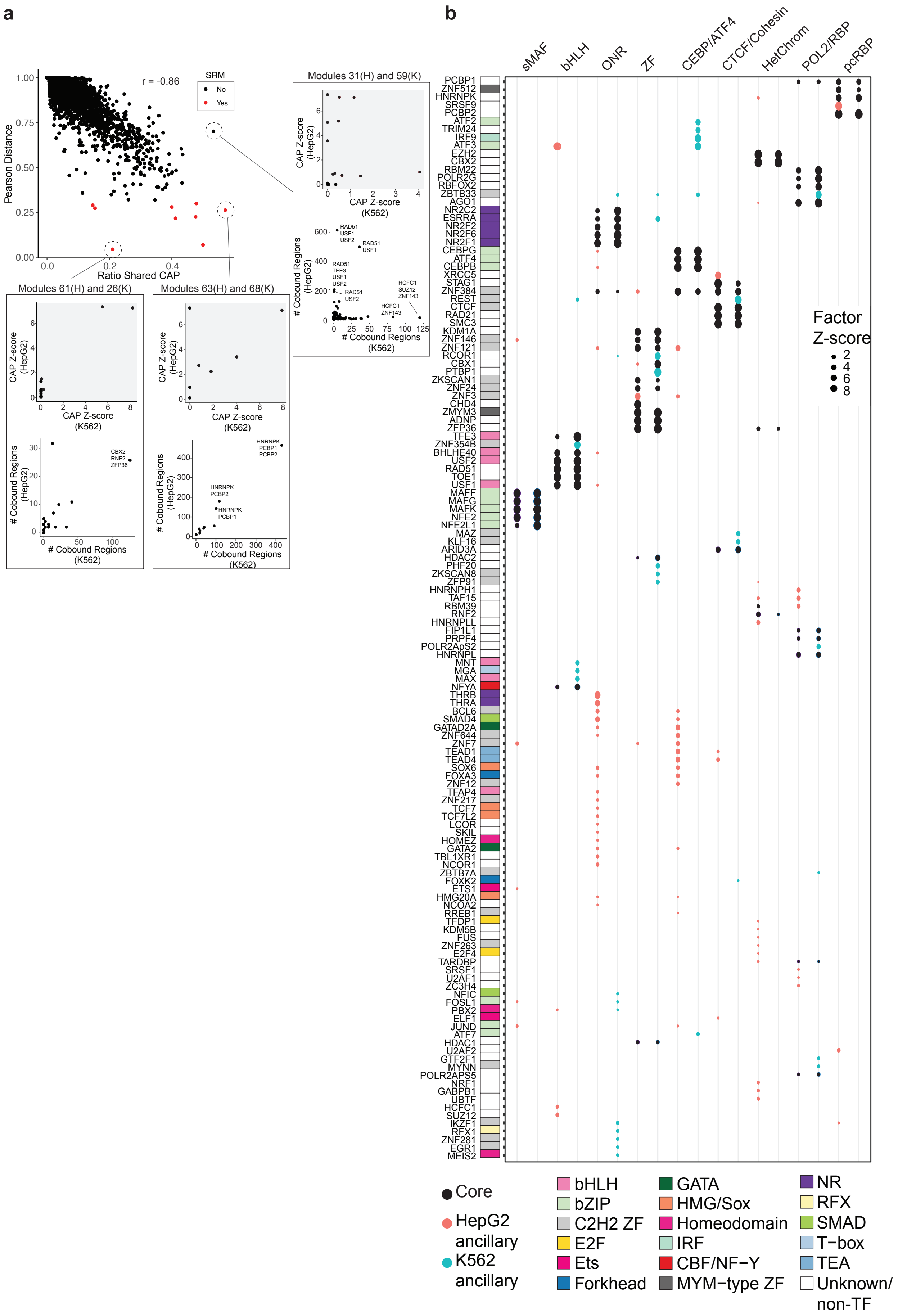

a
b
r = -0.86
SRM
No
Yes
Pearson Distance
Ratio Shared CAP
Modules 31(H) and 59(K)
Modules 61(H) and 26(K)
Modules 63(H) and 68(K)
CAP Z-score (HepG2)
CAP Z-score (K562)
### Cobound Regions (HepG2)
### Cobound Regions (K562)
RAD51 USF1 USF2
RAD51 USF1
RAD51 TFE3 USF1 USF2
RAD51 USF2
HCFC1 SUZ12 ZNF143
HCFC1 ZNF143
CBX2 RNF2 ZFP36
HNRNPK PCBP1 PCBP2
HNRNPK PCBP2
HNRNPK PCBP1
sMAF
bHLH
ONR
ZF
CEBP/ATF4
CTCF/Cohesin
HetChrom
POL2/RBP
pcRBP
PCBP1
ZNF512
HNRNPK
SRSF9
PCBP2
ATF2
TRIM24
IRF9
ATF3
EZH2
CBX2
RBM22
POLR2G
RBFOX2
ZBTB33
AGO1
NR2C2
ESRRA
NR2F2
NR2F6
NR2F1
CEBPG
ATF4
CEBPB
XRCC5
STAG1
ZNF384
REST
CTCF
RAD21
SMC3
KDM1A
ZNF146
ZNF121
RCOR1
CBX1
PTBP1
ZKSCAN1
ZNF24
ZNF3
CHD4
ZMYM3
ADNP
ZFP36
TFE3
ZNF354B
BHLHE40
USF2
RAD51
TOE1
USF1
MAFF
MAFG
MAFK
NFE2
NFE2L1
MAZ
KLF16
ARID3A
HDAC2
PHF20
ZKSCAN8
ZFP91
HNRNPH1
TAF15
RBM39
RNF2
HNRNPLL
FIP1L1
PRPF4
POLR2ApS2
HNRNPL
MNT
MGA
MAX
NFYA
THRB
THRA
BCL6
SMAD4
GATAD2A
ZNF644
ZNF7
TEAD1
TEAD4
SOX6
FOXA3
ZNF12
TFAP4
ZNF217
TCF7
TCF7L2
LCOR
SKIL
HOMEZ
GATA2
TBL1XR1
NCOR1
ZBTB7A
FOXK2
ETS1
HMG20A
NCOA2
RREB1
TFDP1
KDM5B
FUS
ZNF263
E2F4
TARDBP
SRSF1
U2AF1
ZC3H4
NFIC
FOSL1
PBX2
ELF1
JUND
ATF7
HDAC1
U2AF2
GTF2F1
MYNN
POLR2APS5
NRF1
GABPB1
UBTF
HCFC1
SUZ12
IKZF1
RFX1
ZNF281
EGR1
MEIS2
Factor Z-score
2
4
6
8
Core
HepG2 ancillary
K562 ancillary
bHLH
bZIP
C2H2 ZF
E2F
Ets
Forkhead
GATA
HMG/Sox
Homeodomain
IRF
CBF/NF−Y
MYM−type ZF
NR
RFX
SMAD
T−box
TEA
Unknown/ non-TF

**Extended Data Figure 3. Shared regulatory modules classification and composition.**

a) Scatterplot depicting the Pearson distance and the ratio of shared CAPs between two module pairs from each cell line. Boxes highlight specific module pairs and plot the CAP z-scores within modules as well as the number of co-bound regions for core CAP subgroups. Note, module pairs with high ratio of shared CAPs and low Pearson distance have similar CAP subgroups whereas module pairs with high ratio of shared CAPs and high Pearson distance exhibit differing CAP subgroups.

b) Dot plot displaying the CAP z-scores for all CAPs, core and ancillary, present at common regulatory modules. TF family annotation is indicated. Data is related to Fig 2a.

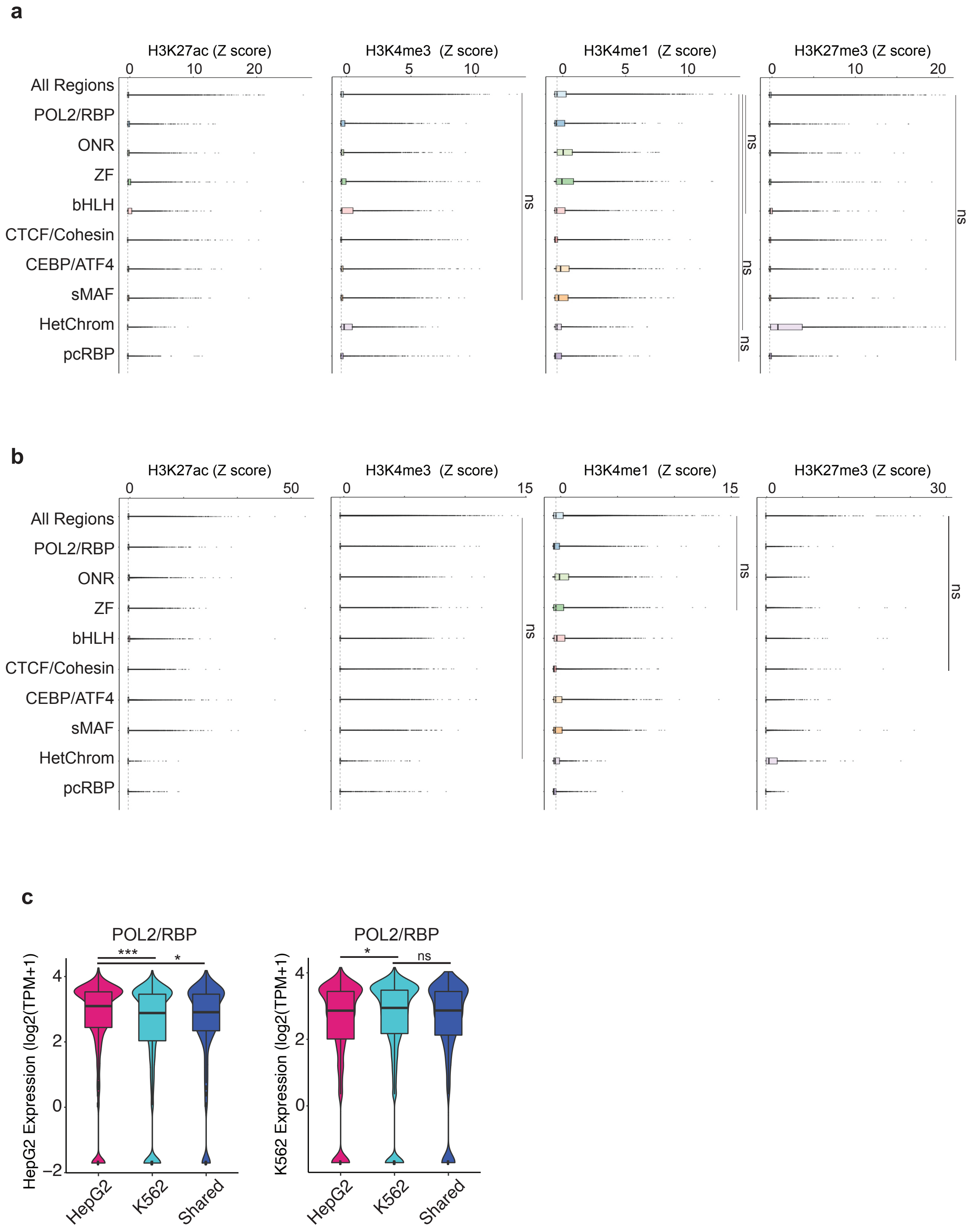


**Extended Data Figure 4. Shared regulatory modules functional state.**

a-b) Boxplots displaying the H3K27ac, H3K4me3, H3K4me1, and H3K27me3 signal distributions across all SRM modules for HepG2 (a) and K562 (b) cell lines. Note unless otherwise stated, all p values are less than 0.005 (Wilcoxon rank sum test).

c) Violin plots displaying the expression level of POL2/RBP HepG2-specific, K562-specific, or shared regulated genes in HepG2 (left) or K562 (right) cells. *** = $p < 0.0005$; * = $p < 0.05$; ns = not significant. All p-values were determined using the Wilcoxon rank sum test.

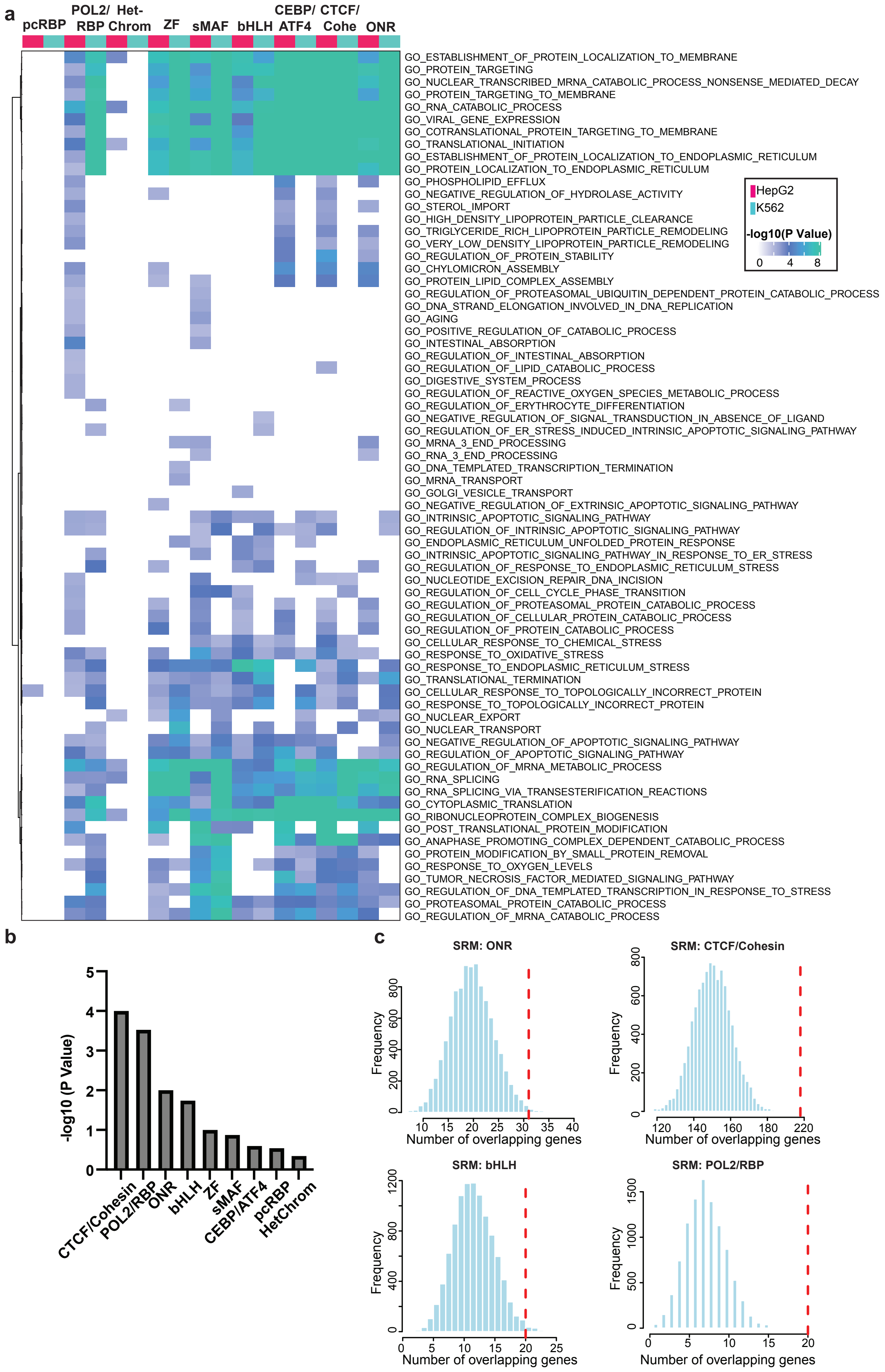
a
pcRBP
POL2/ RBP
Het- Chrom
ZF
sMAF
bHLH
CEBP/ ATF4
CTCF/ Cohe
ONR
GO_ESTABLISHMENT_OF_PROTEIN_LOCALIZATION_TO_MEMBRANE
GO_PROTEIN_TARGETING
GO_NUCLEAR_TRANSCRIBED_MRNA_CATABOLIC_PROCESS_NONSENSE_MEDIATED_DECAY
GO_PROTEIN_TARGETING_TO_MEMBRANE
GO_RNA_CATABOLIC_PROCESS
GO_VIRAL_GENE_EXPRESSION
GO_COTRANSLATIONAL_PROTEIN_TARGETING_TO_MEMBRANE
GO_TRANSLATIONAL_INITIATION
GO_ESTABLISHMENT_OF_PROTEIN_LOCALIZATION_TO_ENDOPLASMIC_RETICULUM
GO_PROTEIN_LOCALIZATION_TO_ENDOPLASMIC_RETICULUM
GO_PHOSPHOLIPID_EFFLUX
GO_NEGATIVE_REGULATION_OF_HYDROLASE_ACTIVITY
GO_STEROL_IMPORT
GO_HIGH_DENSITY_LIPOPROTEIN_PARTICLE_CLEARANCE
GO_TRIGLYCERIDE_RICH_LIPOPROTEIN_PARTICLE_REMODELING
GO_VERY_LOW_DENSITY_LIPOPROTEIN_PARTICLE_REMODELING
GO_REGULATION_OF_PROTEIN_STABILITY
GO_CHYLOMICRON_ASSEMBLY
GO_PROTEIN_LIPID_COMPLEX_ASSEMBLY
GO_REGULATION_OF_PROTEASOMAL_UBIQUITIN_DEPENDENT_PROTEIN_CATABOLIC_PROCESS
GO_DNA_STRAND_ELONGATION_INVOLVED_IN_DNA_REPLICATION
GO_AGING
GO_POSITIVE_REGULATION_OF_CATABOLIC_PROCESS
GO_INTESTINAL_ABSORPTION
GO_REGULATION_OF_INTESTINAL_ABSORPTION
GO_REGULATION_OF_LIPID_CATABOLIC_PROCESS
GO_DIGESTIVE_SYSTEM_PROCESS
GO_REGULATION_OF_REACTIVE_OXYGEN_SPECIES_METABOLIC_PROCESS
GO_REGULATION_OF_ERYTHROCYTE_DIFFERENTIATION
GO_NEGATIVE_REGULATION_OF_SIGNAL_TRANSDUCTION_IN_ABSENCE_OF_LIGAND
GO_REGULATION_OF_ER_STRESS_INDUCED_INTRINSIC_APOPTOTIC_SIGNALING_PATHWAY
GO_MRNA_3_END_PROCESSING
GO_RNA_3_END_PROCESSING
GO_DNA_TEMPLATED_TRANSCRIPTION_TERMINATION
GO_MRNA_TRANSPORT
GO_GOLGI_VESICLE_TRANSPORT
GO_NEGATIVE_REGULATION_OF_EXTRINSIC_APOPTOTIC_SIGNALING_PATHWAY
GO_INTRINSIC_APOPTOTIC_SIGNALING_PATHWAY
GO_REGULATION_OF_INTRINSIC_APOPTOTIC_SIGNALING_PATHWAY
GO_ENDOPLASMIC_RETICULUM_UNFOLDED_PROTEIN_RESPONSE
GO_INTRINSIC_APOPTOTIC_SIGNALING_PATHWAY_IN_RESPONSE_TO_ER_STRESS
GO_REGULATION_OF_RESPONSE_TO_ENDOPLASMIC_RETICULUM_STRESS
GO_NUCLEOTIDE_EXCISION_REPAIR_DNA_INCISION
GO_REGULATION_OF_CELL_CYCLE_PHASE_TRANSITION
GO_REGULATION_OF_PROTEASOMAL_PROTEIN_CATABOLIC_PROCESS
GO_REGULATION_OF_CELLULAR_PROTEIN_CATABOLIC_PROCESS
GO_REGULATION_OF_PROTEIN_CATABOLIC_PROCESS
GO_CELLULAR_RESPONSE_TO_CHEMICAL_STRESS
GO_RESPONSE_TO_OXIDATIVE_STRESS
GO_RESPONSE_TO_ENDOPLASMIC_RETICULUM_STRESS
GO_TRANSLATIONAL_TERMINATION
GO_CELLULAR_RESPONSE_TO_TOPOLOGICALLY_INCORRECT_PROTEIN
GO_RESPONSE_TO_TOPOLOGICALLY_INCORRECT_PROTEIN
GO_NUCLEAR_EXPORT
GO_NUCLEAR_TRANSPORT
GO_NEGATIVE_REGULATION_OF_APOPTOTIC_SIGNALING_PATHWAY
GO_REGULATION_OF_APOPTOTIC_SIGNALING_PATHWAY
GO_REGULATION_OF_MRNA_METABOLIC_PROCESS
GO_RNA_SPLICING
GO_RNA_SPLICING_VIA_TRANSESTERIFICATION_REACTIONS
GO_CYTOPLASMIC_TRANSLATION
GO_RIBONUCLEOPROTEIN_COMPLEX_BIOGENESIS
GO_POST_TRANSLATIONAL_PROTEIN_MODIFICATION
GO_ANAPHASE_PROMOTING_COMPLEX_DEPENDENT_CATABOLIC_PROCESS
GO_PROTEIN_MODIFICATION_BY_SMALL_PROTEIN_REMOVAL
GO_RESPONSE_TO_OXYGEN_LEVELS
GO_TUMOR_NECROSIS_FACTOR_MEDIATED_SIGNALING_PATHWAY
GO_REGULATION_OF_DNA_TEMPLATED_TRANSCRIPTION_IN_RESPONSE_TO_STRESS
GO_PROTEASOMAL_PROTEIN_CATABOLIC_PROCESS
GO_REGULATION_OF_MRNA_CATABOLIC_PROCESS
HepG2
K562
-log10(P Value)
0 4 8
b
-log10 (P Value)
0 1 2 3 4 5
CTCF/Cohesin
POL2/RBP
ONR
bHLH
ZF
sMAF
CEBP/ATF4
pcRBP
HetChrom
c
SRM: ONR
Frequency
0 200 400 600 800
10 15 20 25 30 35 40
Number of overlapping genes
SRM: CTCF/Cohesin
Frequency
0 200 400 600 800
120 140 160 180 220
Number of overlapping genes
SRM: bHLH
Frequency
0 400 800 1200
0 5 10 15 20 25
Number of overlapping genes
SRM: POL2/RBP
Frequency
0 500 1000 1500
0 5 10 15 20
Number of overlapping genes

**Extended Data Figure 5. Shared regulatory modules pathway enrichment.**

a) Heatmap showing GO pathways enriched across all SRMs in HepG2 and K562 cells. For each SRM, the top 500 expressed genes were used for pathway enrichment analysis and only pathways with significant p values ($p < 0.01$) are plotted.

b) Bar plot depicting the p values from 10,000 permutation tests measuring the significant of HepG2 and K562 module regions targeting the same gene promoter via distal interactions. To do this HepG2 and K562 SRM module regions were overlapped with PCHi-C "other" ends, mapped to "bait" genes, and the number of target genes that overlapped were measured. This was compared to randomly selected, same-sized CAP-bound regions instead of SRM regions (see Methods).

c) Histograms depicting the significant results from the permutation tests described in (b). The histogram shows the distribution of the number of overlapping target genes in random HepG2 and K562 CAP-bound regions compared to the observed number in each SRM (red dotted line).

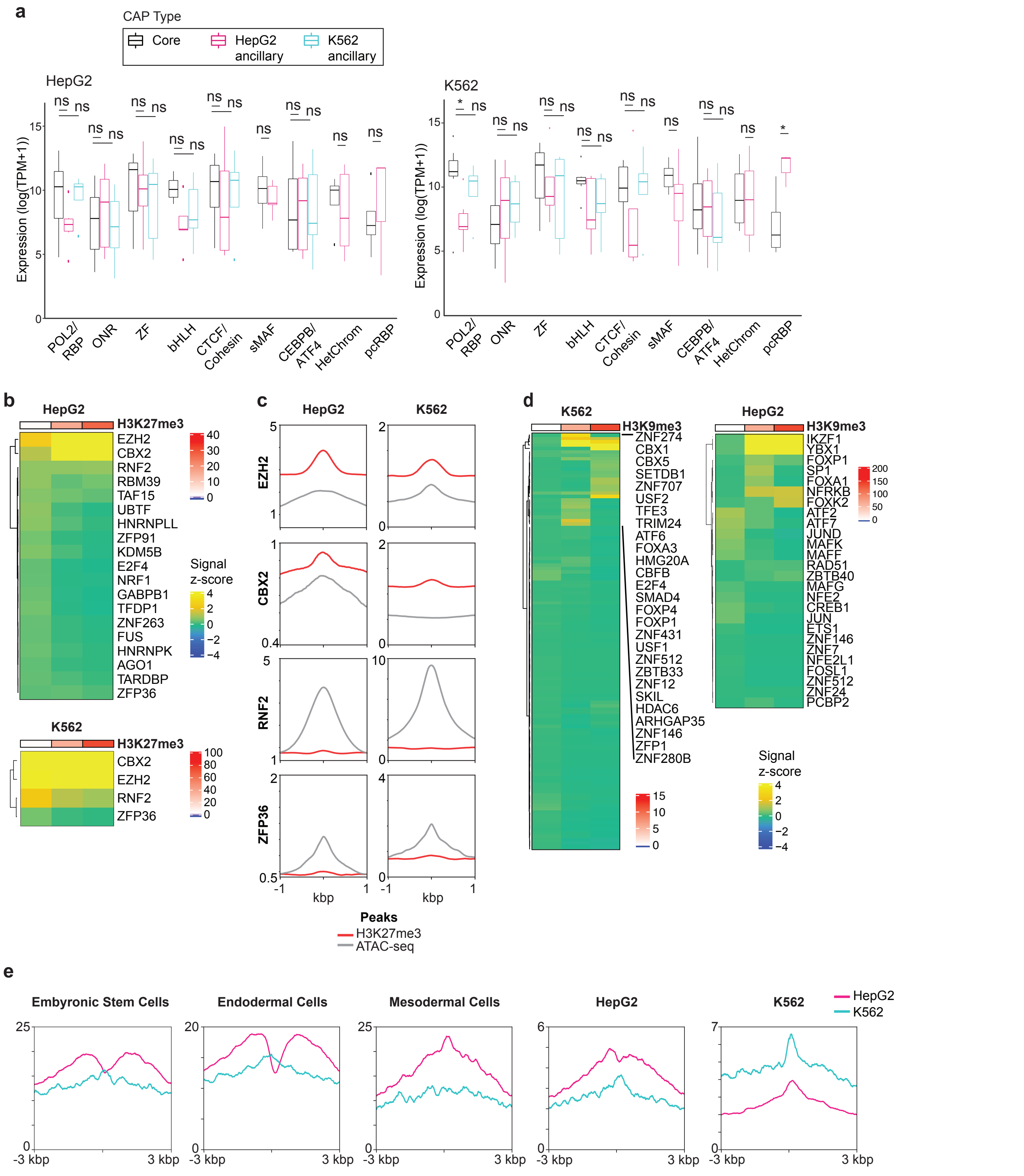


**Extended Data Figure 6. Shared regulatory modules CAP comparison and HetChrom lineage analysis.**

a) Boxplots displaying the expression levels of core and ancillary CAPs in HepG2 (left) and K562 (right) cell lines. ns = not significant. All p-values were determined using the Wilcoxon rank sum test.

b) Heatmap of ChIP-seq signal z-score for the indicated HetChrom SRM CAPs in HepG2 (top) and K562 (bottom) cell lines. H3K27me3+ SRM regions were binned into 3 groups and average CAP signal z-score was measured.

c) ChIP-seq signal profile plots of the indicated HetChrom core CAPs at HetChrom SRM regions compared to all ATAC-seq regions in HepG2 and K562 cell lines.

d) Heatmap of ChIP-seq signal z-score for the indicated CAPs in HepG2 (top) and K562 (bottom) cell lines. H3K9me3+ SRM regions were binned into 3 groups and average CAP signal z-score was measured.

e) H3K27me3 profile plots at either HepG2 or K562 HetChrom SRM regions in the indicated cell types.

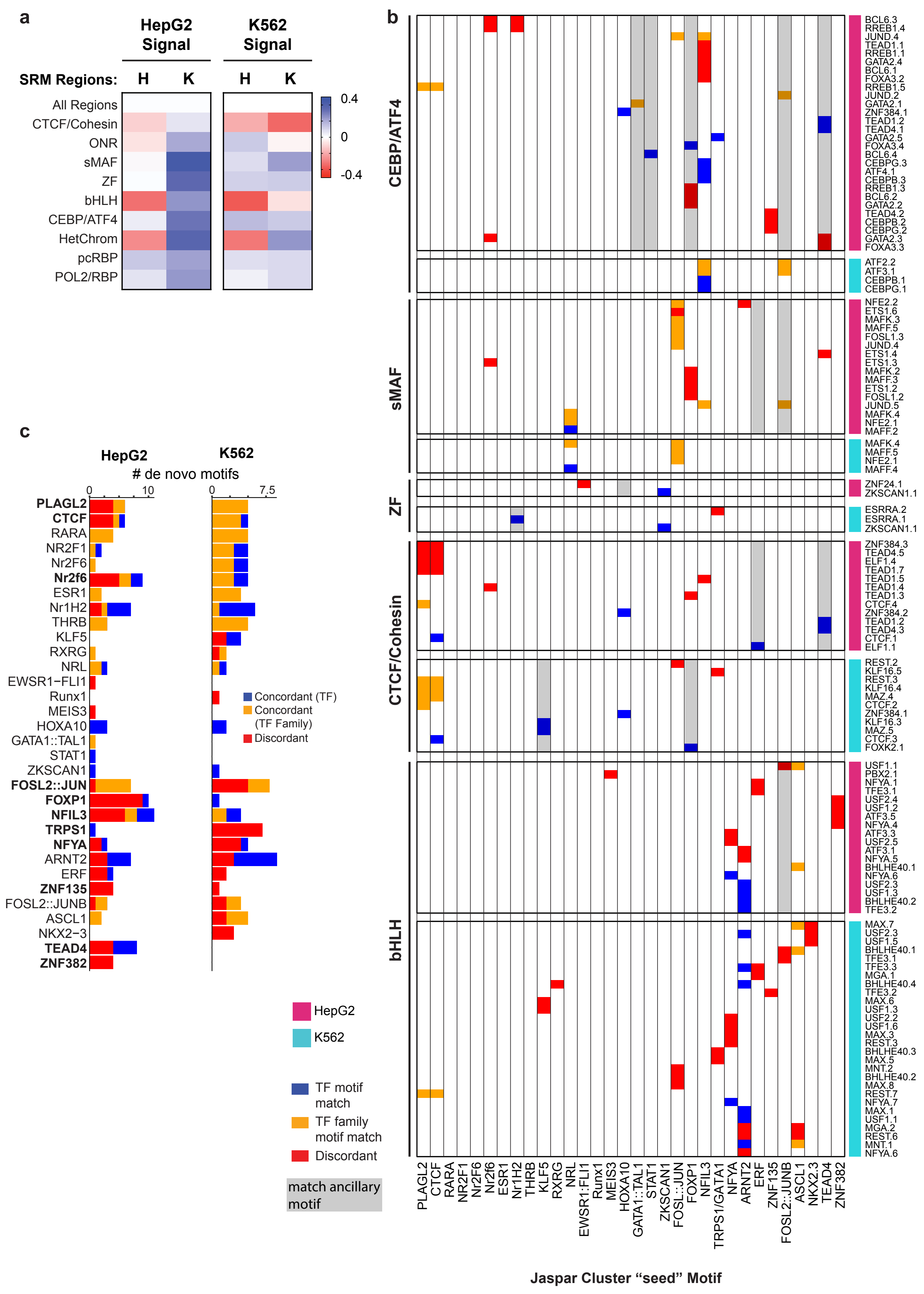
a
HepG2 Signal
K562 Signal
SRM Regions:
H
K
H
K
All Regions
CTCF/Cohesin
ONR
sMAF
ZF
bHLH
CEBP/ATF4
HetChrom
pcRBP
POL2/RBP
0.4
0
-0.4
b
CEBP/ATF4
sMAF
ZF
CTCF/Cohesin
bHLH
BCL6.3
RREB1.4
JUND.4
TEAD1.1
RREB1.1
GATA2.4
BCL6.1
FOXA3.2
RREB1.5
JUND.2
GATA2.1
ZNF384.1
TEAD1.2
TEAD4.1
GATA2.5
FOXA3.4
BCL6.4
CEBPG.3
ATF4.1
CEBPB.3
RREB1.3
BCL6.2
GATA2.2
TEAD4.2
CEBPB.2
CEBPG.2
GATA2.3
FOXA3.3
ATF2.2
ATF3.1
CEBPB.1
CEBPG.1
NFE2.2
ETS1.6
MAFK.3
MAFF.5
FOSL1.3
JUND.4
ETS1.4
ETS1.3
MAFK.2
MAFF.3
ETS1.2
FOSL1.2
JUND.5
MAFK.4
NFE2.1
MAFF.2
MAFK.4
MAFF.5
NFE2.1
MAFF.4
ZNF24.1
ZKSCAN1.1
ESRRA.2
ESRRA.1
ZKSCAN1.1
ZNF384.3
TEAD4.5
ELF1.4
TEAD1.7
TEAD1.5
TEAD1.4
TEAD1.3
CTCF.4
ZNF384.2
TEAD1.2
TEAD4.3
CTCF.1
ELF1.1
REST.2
KLF16.5
REST.3
KLF16.4
MAZ.4
CTCF.2
ZNF384.1
KLF16.3
MAZ.5
CTCF.3
FOXK2.1
USF1.1
PBX2.1
NFYA.1
TFE3.1
USF2.4
USF1.2
ATF3.5
NFYA.4
ATF3.3
USF2.5
ATF3.1
NFYA.5
BHLHE40.1
NFYA.6
USF2.3
USF1.3
BHLHE40.2
TFE3.2
MAX.7
USF2.3
USF1.5
BHLHE40.1
TFE3.1
TFE3.3
MGA.1
BHLHE40.4
TFE3.2
MAX.6
USF1.3
USF2.2
USF1.6
MAX.3
REST.3
BHLHE40.3
MAX.5
MNT.2
BHLHE40.2
MAX.8
REST.7
NFYA.7
MAX.1
USF1.1
MGA.2
REST.6
MNT.1
NFYA.6
PLAGL2
CTCF
RARA
NR2F1
Nr2f6
Nr2f6
ESR1
Nr1H2
THRB
KLF5
RXRG
NRL
EWSR1:FLI1
Runx1
MEIS3
HOXA10
GATA1::TAL1
STAT1
ZKSCAN1
FOSL::JUN
FOXP1
NFIL3
TRPS1/GATA1
NFYA
ARNT2
ERF
ZNF135
FOSL2::JUNB
ASCL1
NKX2.3
TEAD4
ZNF382
Jaspar Cluster “seed” Motif
HepG2
K562
TF motif match
TF family motif match
Discordant
match ancillary motif
c
HepG2
K562
### de novo motifs
0
10
0
7.5
PLAGL2
CTCF
RARA
NR2F1
Nr2F6
Nr2f6
ESR1
Nr1H2
THRB
KLF5
RXRG
NRL
EWSR1−FLI1
Runx1
MEIS3
HOXA10
GATA1::TAL1
STAT1
ZKSCAN1
FOSL2::JUN
FOXP1
NFIL3
TRPS1
NFYA
ARNT2
ERF
ZNF135
FOSL2::JUNB
ASCL1
NKX2−3
TEAD4
ZNF382
Concordant (TF)
Concordant (TF Family)
Discordant

**Extended Data Figure 7. *De novo* motif discovery at cell type-specific SRM regions.**

a) Heatmap displaying the average ATAC-seq z-scores across the SRM regions of the indicated cell type. Note HepG2 ATAC-seq signal decreases at all K562 SRM regions compared to the corresponding HepG2 SRM regions. H: HepG2, K: K562.

b) Heatmap clustered by cell line and SRM displaying the significant Tomtom *de novo* motifs for the indicated TFs. All motif matches were aggregated according to JASPAR motif cluster before plotting. Heatmap is colored according to whether the motif matches the exact TF motif, a motif within the same TF family, or a discordant motif. Motif clusters matching ancillary CAPs for each SRM are highlighted in gray.

c) Bar plot displaying the number of significant Tomtom *de novo* identified for each JASPAR motif cluster and colored according to TF concordance, TF family concordance, or discordant. Note the JASPAR motif cluster is named based on the "seed" motif.

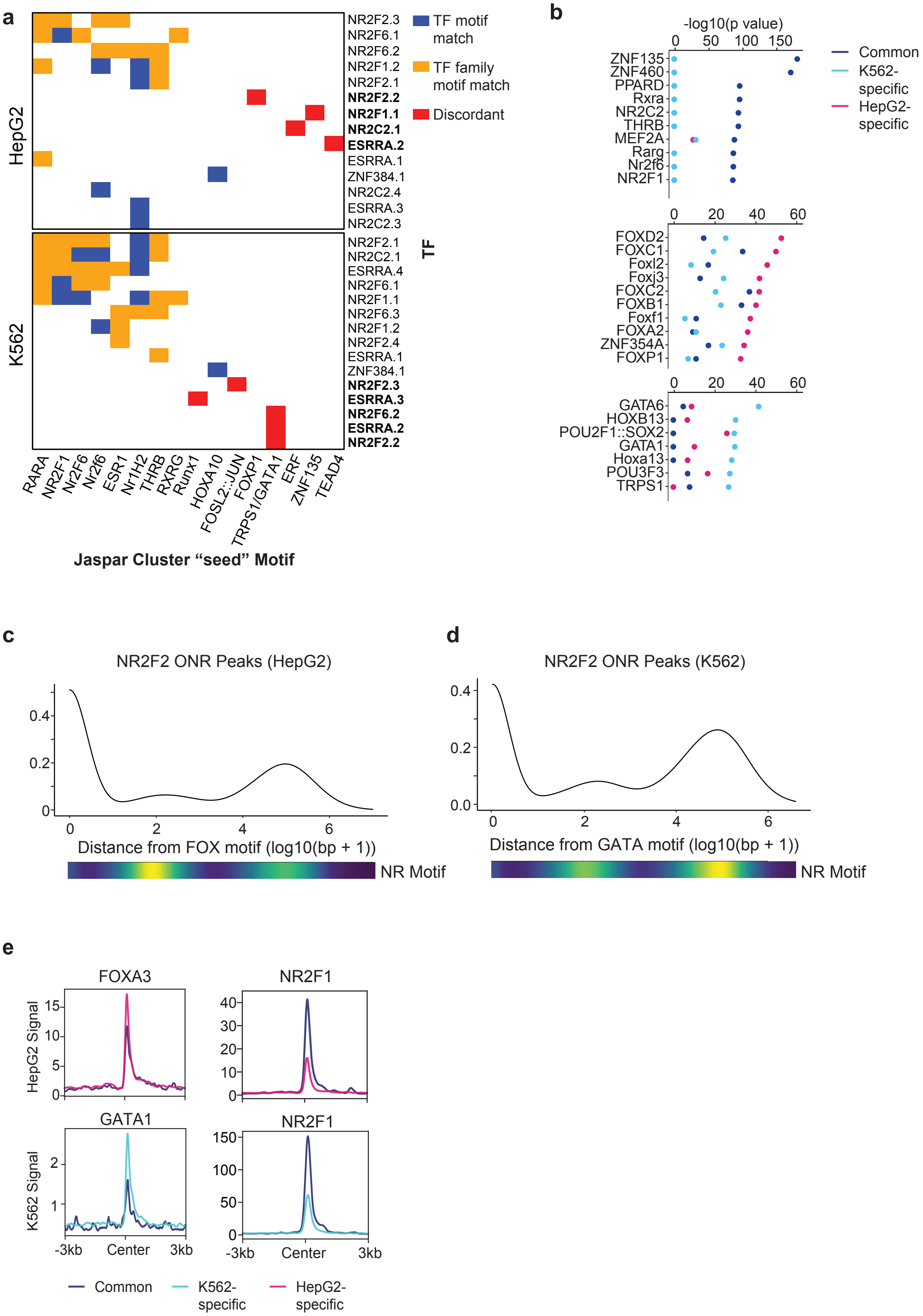


**Extended Data Figure 8. Shared and cell type-specific motif discovery at ONR modules .**

a) Heatmap displaying the significant Tomtom *de novo* motifs discovered from TF ChIP-seq peaks present within the orphan nuclear receptor SRM. Heatmap is colored according to whether the motif matches the exact TF motif, a motif within the same TF family, or a discordant motif. TFs with discordant motif matches are bolded.

b) XSTREME motif discovery enrichment at ONR shared and cell type-specific regions.

c-d) Density plots displaying the peak position of the indicated TF (top) or motif (bottom) relative to the FOX (c) or GATA (d) motif positions at ONR module regions.

e) Regions from ONR module were grouped into "Common", "K562-specific", or "HepG2-specific" indicating whether the regions were present in ONR modules in both cell lines or only a single cell line. Density plots and corresponding box plots displaying the indicated TF signals at common and cell type-specific ONR regions.

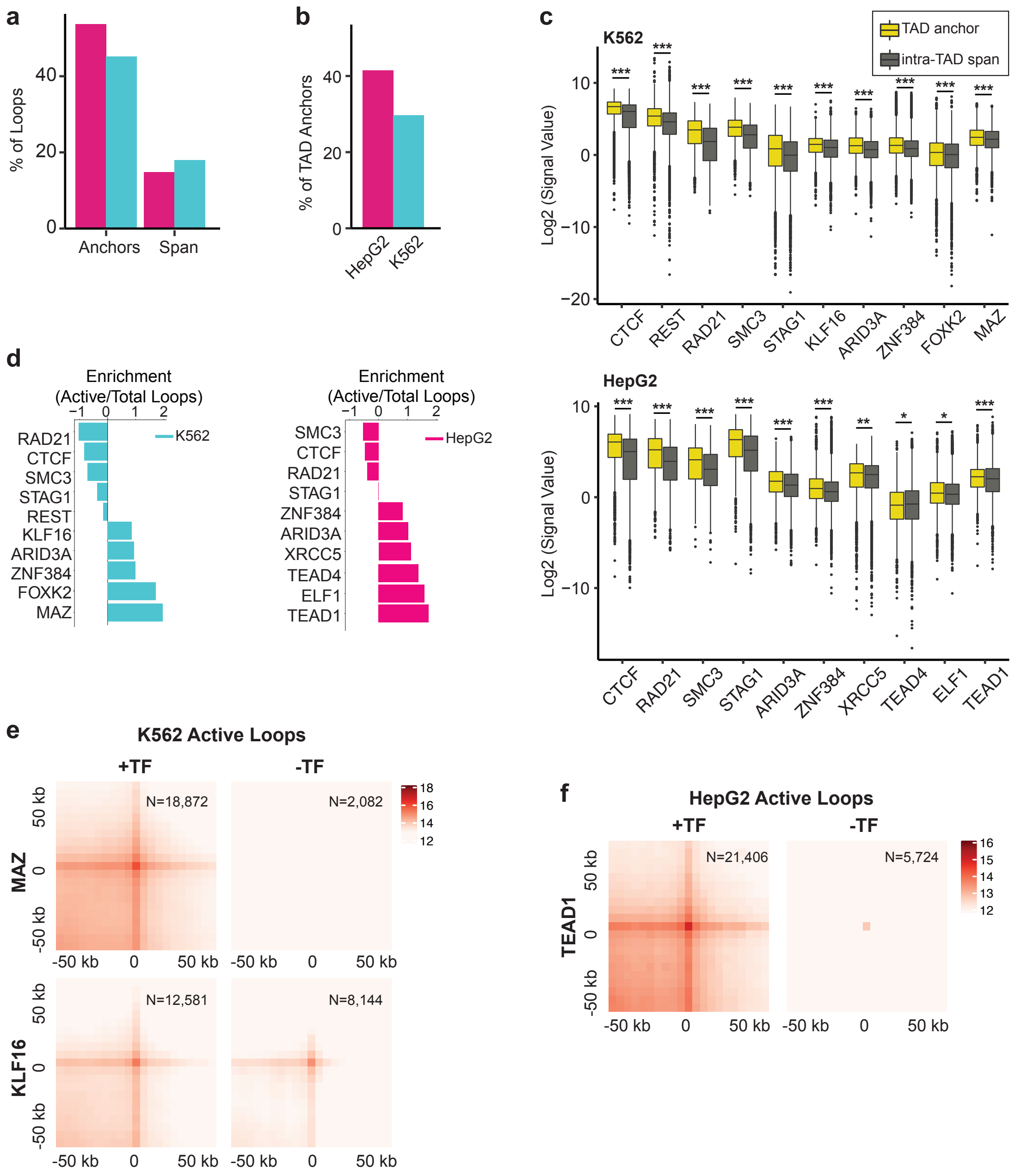


**Extended Data Figure 9. Characteristics of CTCF/Cohesin SRMs.**

a-b) Bar plot displaying the percent of Hi-C loop anchors or intra-loop regions (a) or TAD boundaries (b) bound by CTCF/Cohesin SRM regions in HepG2 and K562 cells.

c) Boxplots displaying the ChIP-seq signal value for all CAPs constituting K562 (top) or HepG2 (bottom) CTCF/Cohesin SRMs at module bound TAD boundaries or intra-TAD regions. *** = $p < 0.0005$; ** = $p < 0.005$; * = $p < 0.05$; ns = not significant. All p-values were determined using the Wilcoxon rank sum test.

d) Bar plot showing the enrichment of active (H3K27ac peak at either or both loop ends) Hi-C loops bound over total Hi-C loops for all CAPs identified at K562 or HepG2 CTCF/Cohesin SRMs. Note KLF16 and MAZ are enriched at K562 active loops compared to all loops and TEAD1 is enriched at HepG2 active loops.

e-f) APA plots displaying aggregated signals across active loops containing or lacking the indicated CAPs in K562 (e) or HepG2 (f) cells. The number of loops used for each plot is indicated in the top right corner.

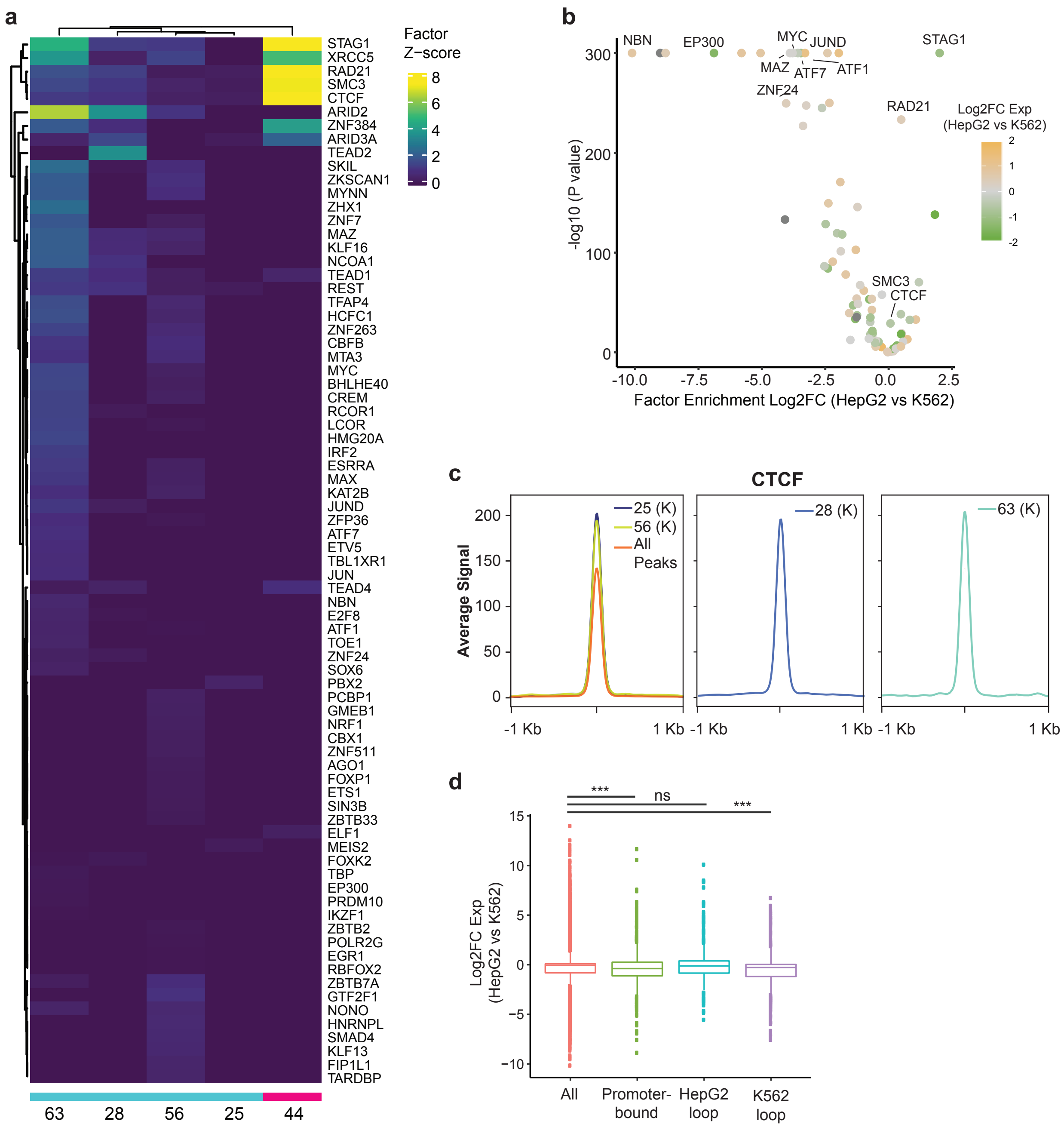


**Extended Data Figure 10. Dynamics of differentially bound modules.**

a) Heatmap displaying CAP z-scores for indicated CAPs across all HepG2 and K562 DBMs.

b) Scatterplot of CAP enrichment at DBMs. Points are colored according to CAP expression (TPM) log2FC in HepG2 compared to K562 cells.

c) ChIP-seq signal profile plots of CTCF at the CTCF peaks present in the indicated DBMs or all CTCF peaks in K562 cells.

d) Box plot showing log2FC expression levels of either all expressed genes ("All"), genes where DBM regions overlap promoters ("Promoter-bound"), or genes where DBM regions overlap distal interactors (PCHi-C "other" ends) in each cell line ("HepG2 loop" or "K562 loop"). *** = $p < 0.0005$; ns = not significant. All P-values were determined using the Wilcoxon rank sum test.

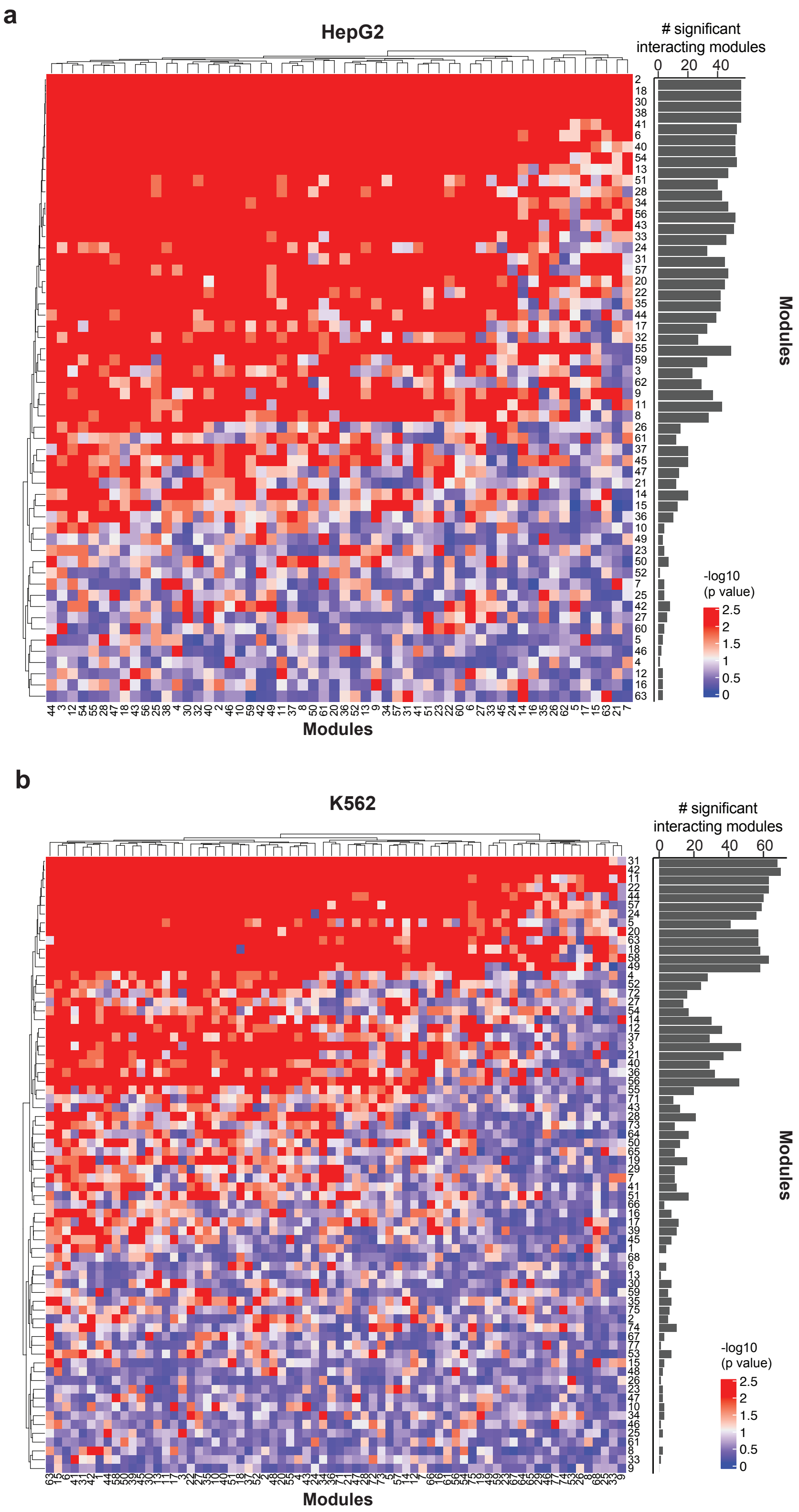


**Extended Data Figure 11. PCHi-C module-module interactions.**

a-b) Heatmap (left) displaying the p-values following 100 permutation tests measuring the number of module-module interacting PCHi-C loops compared to random overlap across same-sized regions (see Methods) and a bar plot (right) showing the number of modules each module was found to interact with via PCHi-C loops ($p < 0.01$). Analyses were conducted using HepG2 (a) and K562 (b) regulatory modules and PCHi-C data.

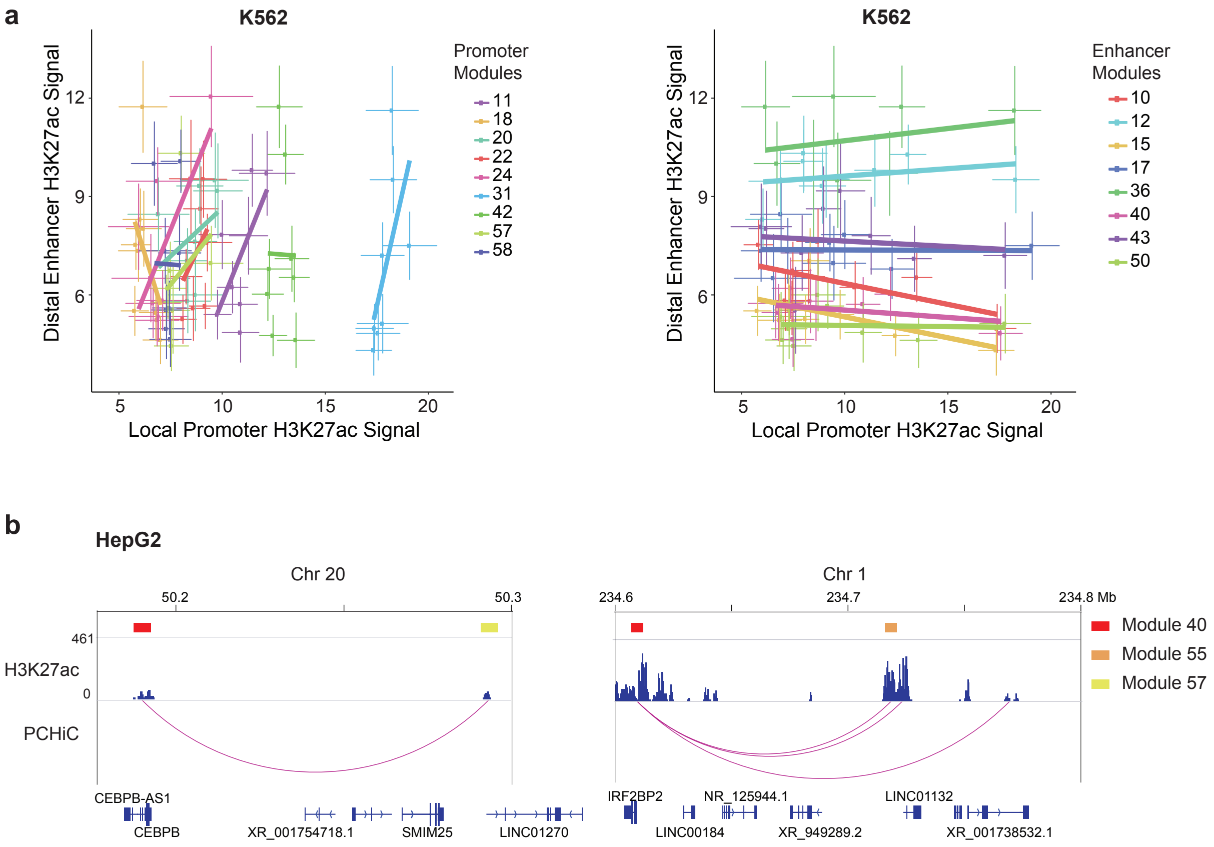


**Extended Data Figure 12. PCHi-C interactions between promoter-associated and enhancer-associated modules.**

a) Scatterplot of the K562 H3K27ac signal value at bait/promoter regions (x axis) and other end/distal regions (y axis) for interacting module pairs. Lines indicate standard error values. Module interactions and linear trend lines are colored by promoter-associated modules (left) or enhancer-associated modules (right).

b) Diagram displaying PCHi-C interactions amongst HepG2 module pairs 40-55 and 40-57. Module 40 represents a promoter module and modules 55 and 57 represent two enhancer modules with varying H3K27ac signal at module 40 interactions. Note module pair 40-55 has a higher H3K27ac signal at both bait (module 40 bound) and other end (module 55 bound) compared to the module 40-57 interaction shown.

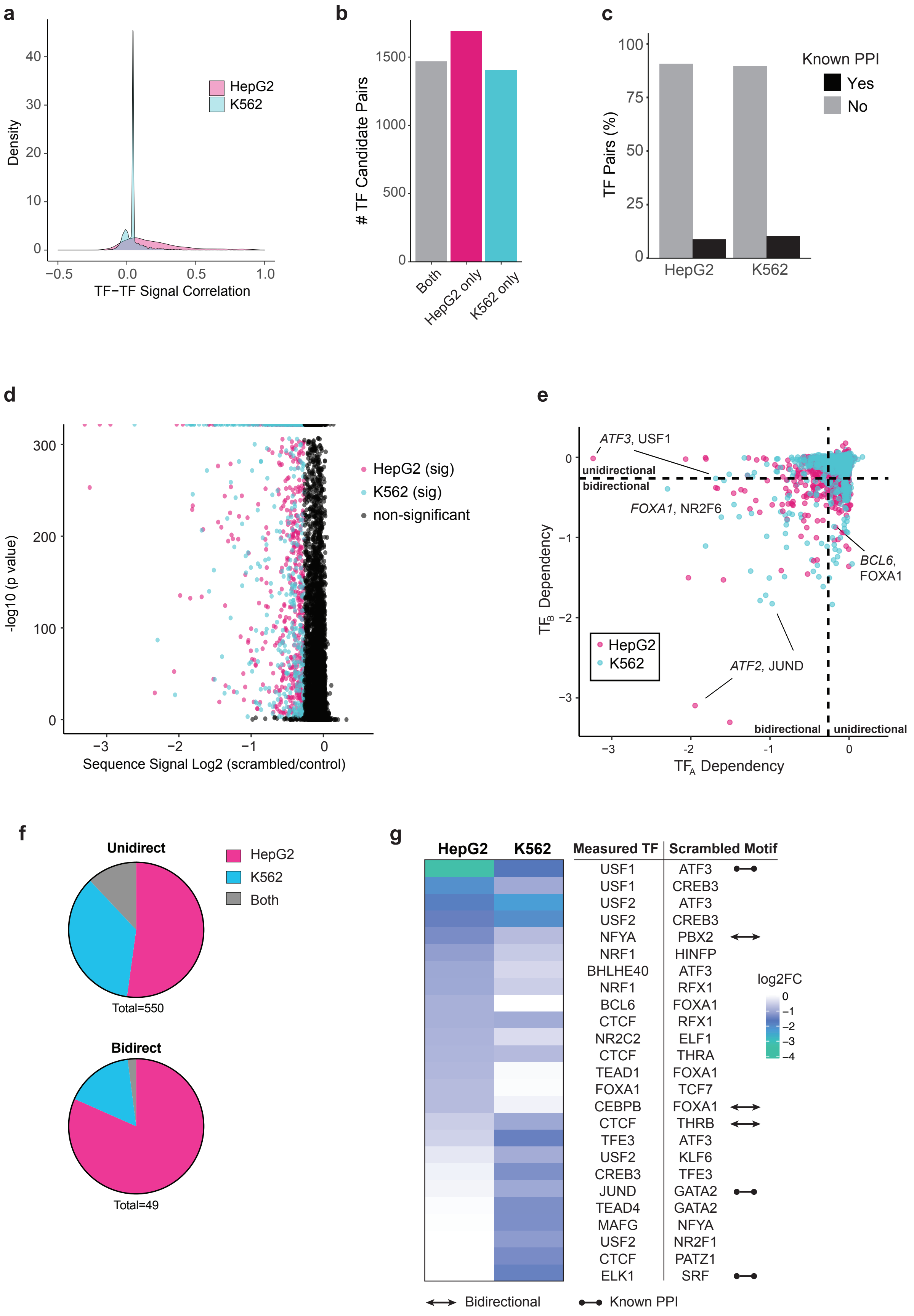


**Extended Data Figure 13. Assessment of TF pairs as candidate cooperators.**

a) Density plots displaying the correlation between signal values candidate TF-TF pairs at overlapping regions in the indicated cell lines.

b) The number of candidate TF pairs that are shared or unique to each cell line based on their presence in regulatory modules and positive ChIP-seq signal correlations at overlapping regions.

c) Bar plot displaying the percent of candidate TF pairs with known protein-protein interactions.

d) Volcano plot depicting the log2 fold change (predicted read count at motif-scrambled sequence / predicted read count at unmodified sequence) versus p-value (paired t-test). Blue or pink points depict TF pairs with significant cooperative prediction results. Significant values were defined as -1.2 < log2 (fold change) or log2 (fold change) > 1.2 with p values < 0.001.

e) Scatter plot displaying the log2 fold change (predicted read count at motif-scrambled sequence / predicted read count at unmodified sequence) of $TF_B$ signal (i.e. $TF_A$ dependency, x-axis) vs $TF_A$ signal (i.e. $TF_B$ dependency, y-axis). TF pairs with significant dependencies in both directions ($TF_A$ signal at $TF_B$ motif-modified sequences and $TF_B$ signal at $TF_A$ motif-modified sequences) are labeled as bidirectional whereas TF pairs with significant dependencies in one direction are labeled as unidirectional. $TF_A$ = italic, $TF_B$ = regular font. *ATF2,* JUN and *JUND*, CREB1 represent positive control TF pairs.

f) Pie chart displaying the ratio of significant TF pairs found in either cell line or both cell lines as well as if the TF pairs displayed bidirectional or unidirectional dependencies. Only TF pairs with different DNA binding domains are plotted.

g) Table displaying the top 15 significant TF pairs identified in either HepG2, K562, or both cell lines. For a complete table of significant TF pairs see **Extended Data Table 10.** Only TF pairs with different DNA binding domains were plotted.

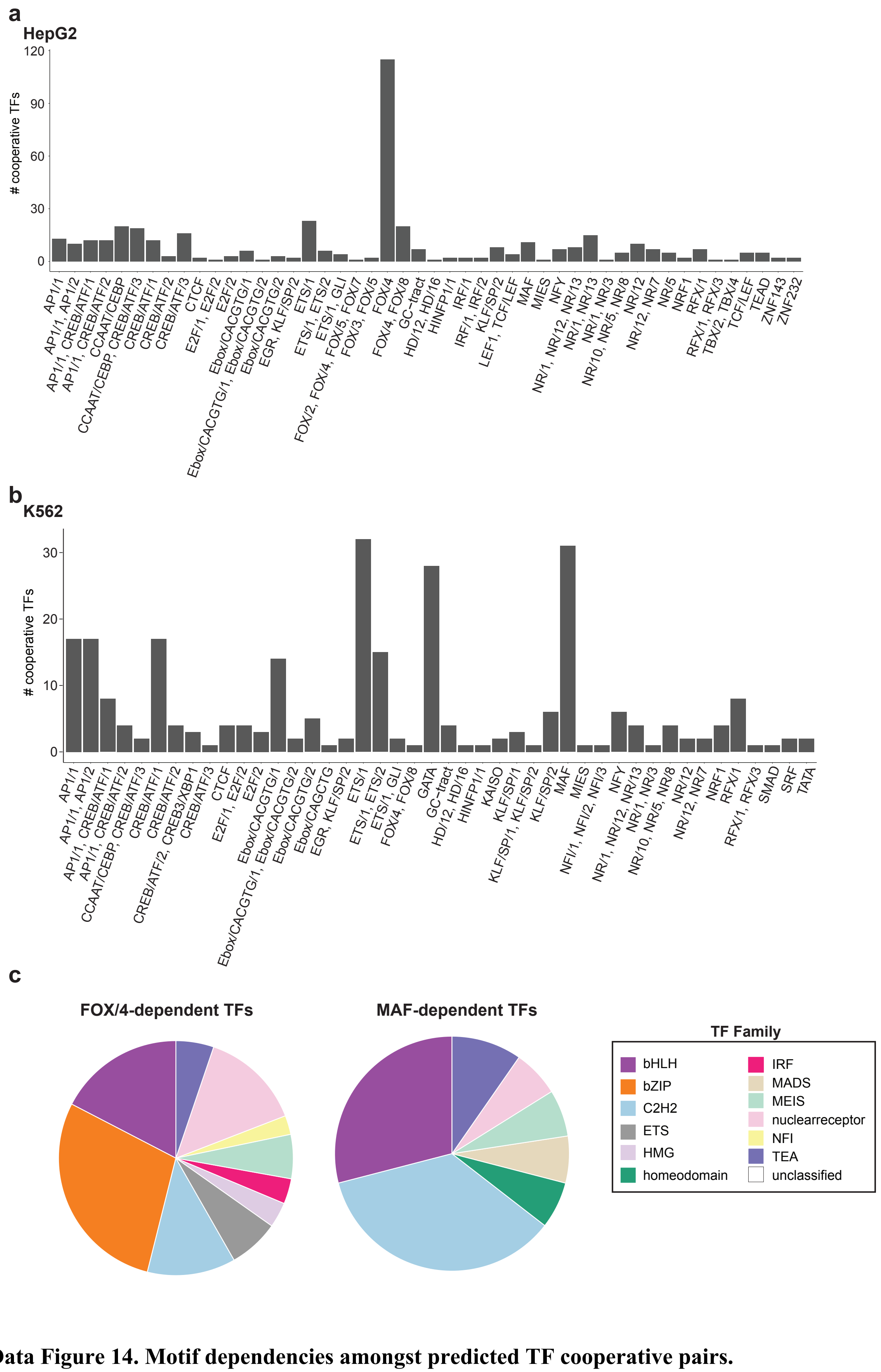


**Extended Data Figure 14. Motif dependencies amongst predicted TF cooperative pairs.**

a-b) Bar plots displaying the number of TFs predicted to display significant cooperative activity when the focus motif (x-axis) was scrambled in HepG2 (a) or K562 (b) ChIP-seq models. Focus motifs (x-axis) were aggregated according to their Viestra motif cluster. Only TF pairs with different DNA binding domains were plotted.

c) Pie plots displaying the distribution of TF DNA-binding domain families exhibiting predicted dependencies on FOX/4 motifs in HepG2 (left) or MAF motifs in K562 (right). Only TF pairs with different DNA binding domains were plotted.

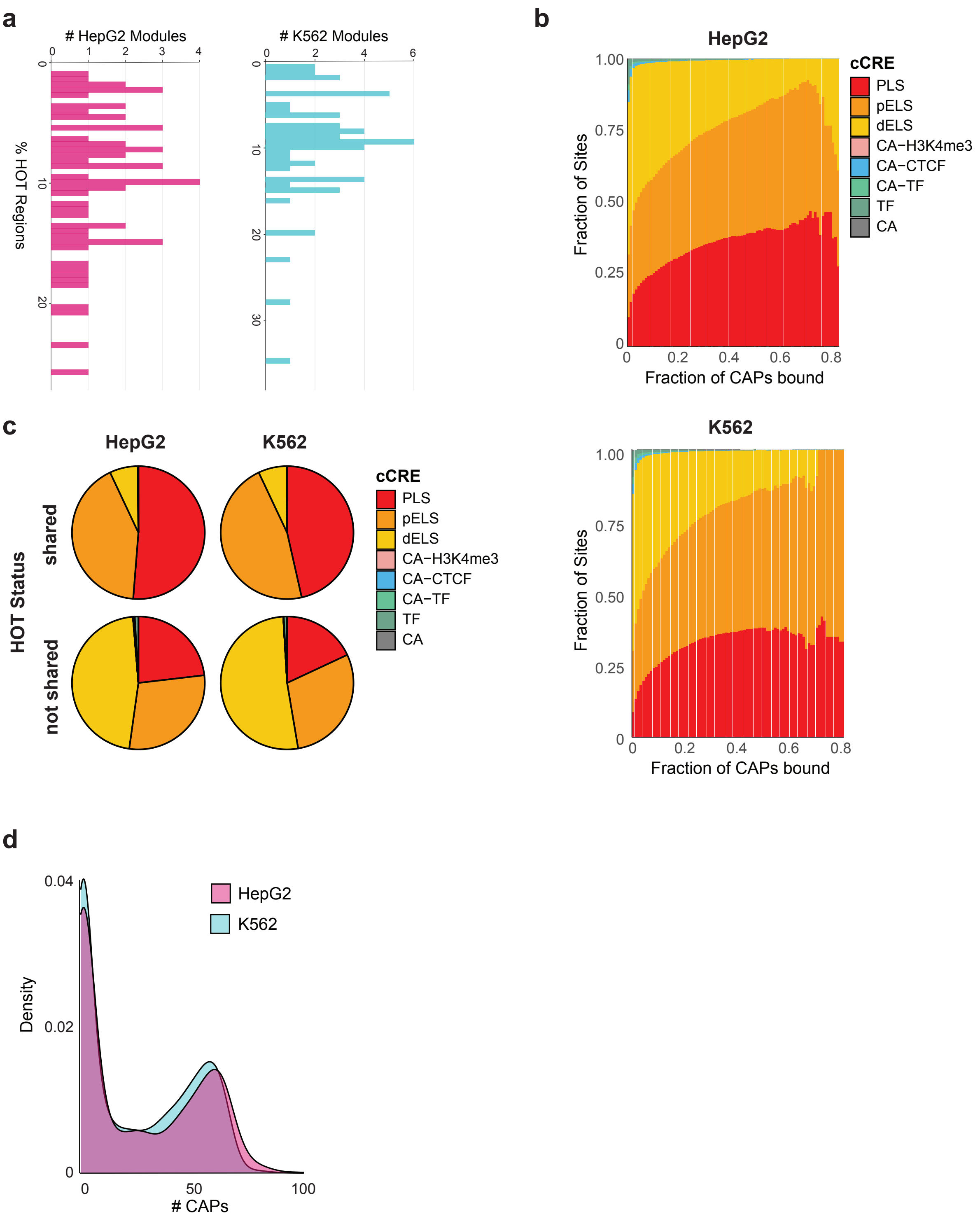


**Extended Data Figure 15. HOT site Characteristics across cell lines.**

a) Histograms displaying the percentage of HOT sites present within regulatory modules for HepG2 and K562 cell lines.

b) Fraction of HOT sites occurring in a particular kind of cCRE in HepG2 (left) or K562 (right) as the threshold for "HOTness" is increased.

c) HOT-site segmentation at shared or cell type-specific HOT sites for the either HepG2 (left) or K562 (right) cells.

d) Density plot of the number of CAPs bound at regions that are not-HOT for the plotted cell line but are HOT for the alternate cell line. For example, the plot labeled “HepG2” shows the number of factors bound in HepG2 at regions that are only HOT in K562.

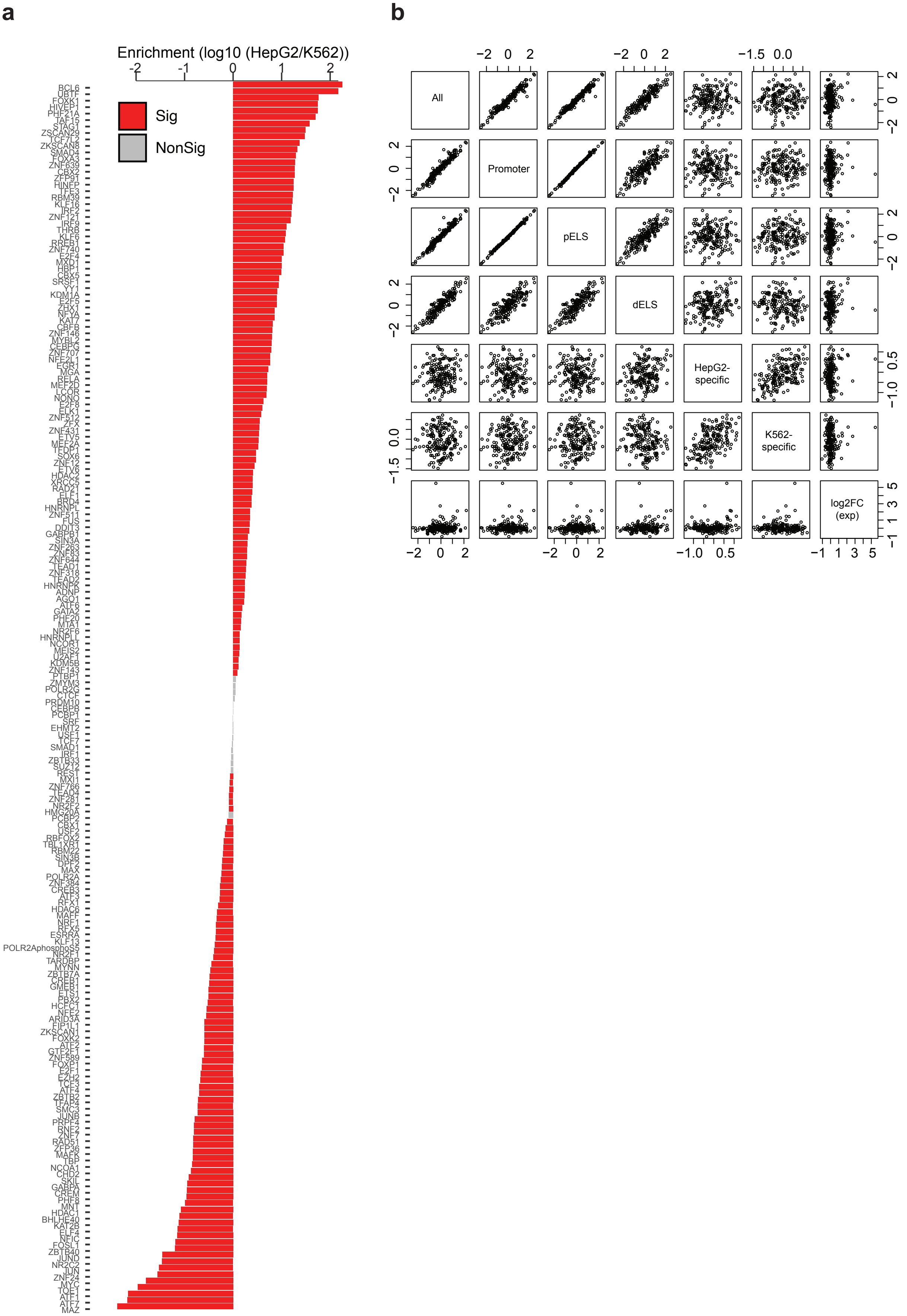


**Extended Data Figure 16. CAP enrichment at HOT sites.**

a) Log10 fold enrichment of factors in HOT sites of one cell type compared to HOT sites in the opposite cell type, ordered from highest to lowest. Positive numbers show enrichment for HepG2 while negative numbers show enrichment in K562. Enrichment was significant if the p-value was <= 0.005 (Chi-squared test). Sig: significant enrichment; NonSig: non-significant enrichment.

b) Log2 enrichment of TFs in HOT sites of different classes. "All" refers to looking at factor enrichment among all HOT sites between HepG2 and K562. "Promoter" is the same, but restricted to HOT sites overlapping with PLS cCRE annotations. "pELS" is the same, but restricted to HOT sites overlapping with pELS cCRE annotations. "dELS" is the same, but restricted to HOT sites overlapping with dELS cCRE annotations. "HepG2-specific" show enrichment for factors in HepG2 HOT sites which are cold in K562 versus HepG2 HOT sites which are neither cold nor HOT in K562. "K562-specific" show enrichment for factors in K562 HOT sites which are cold in HepG2 versus K562 HOT sites which are neither cold nor HOT in HepG2. Finally, "log2FC (exp)" refers only to the logRatio of expression in HepG2 over K562.

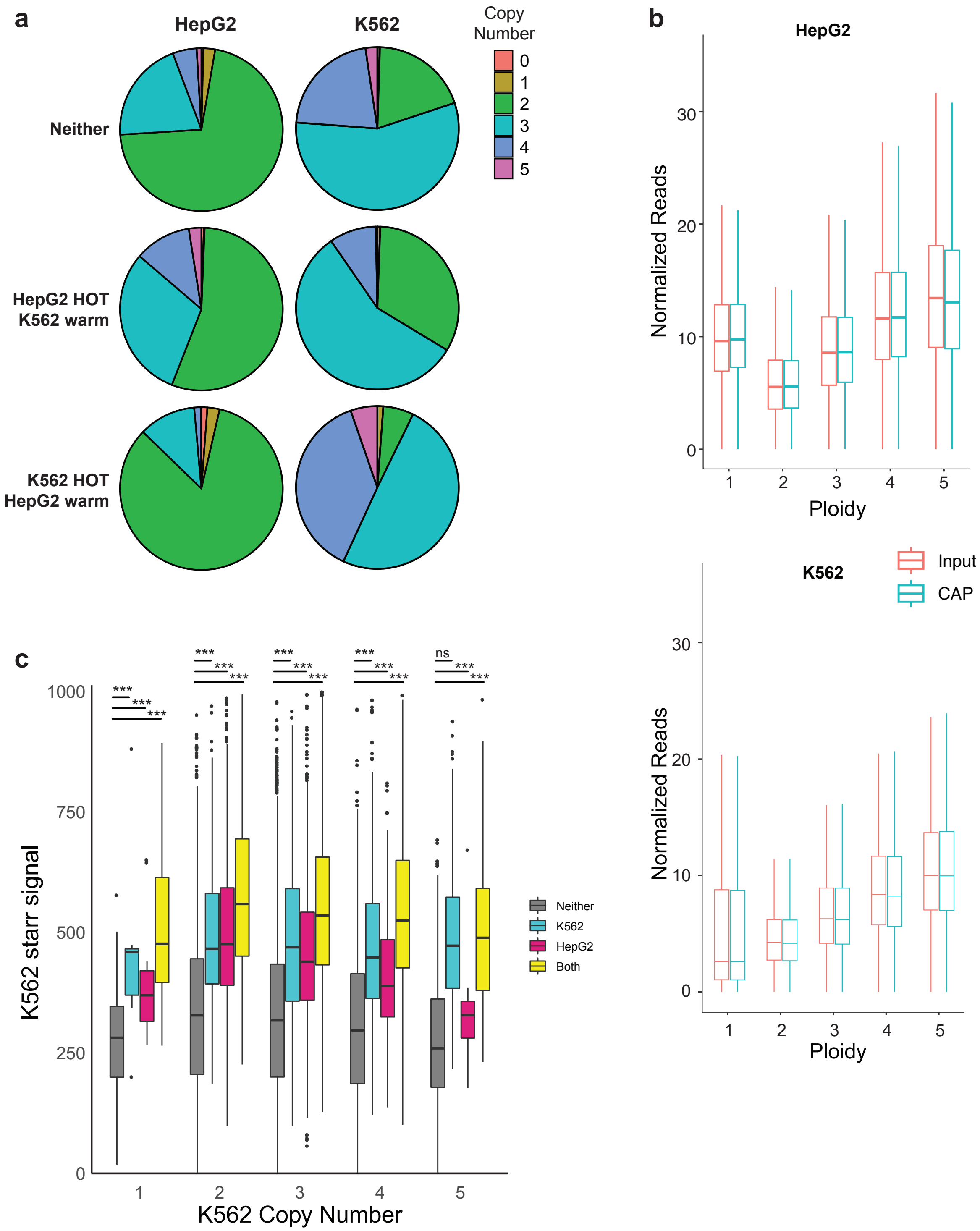


**Extended Data Figure 17. Chromosomal ploidy and HOT site specificity.**

a) Pie plots displaying the HepG2 and K562 copy number distribution for the indicated HOT site categories. “Warm” HOT sites refer to sites that are HOT in one cell type and contain 11-62 bound CAPs in the other cell line.

b) Box plots showing normalized read counts across chromosomal copy number for experiment input control and CAP experiment bam files in HepG2 (top) and K562 (bottom) cell lines.

c) STARR-seq activity levels for K562 genes whose promoters are HOT in either cell type, both, or neither (non-HOT), binned by copy number. ns = not significant; *** = $p < 0.0005$. All p-values were determined using the Wilcoxon rank sum test.
